# Crimean-Congo Hemorrhagic Fever Survivors Elicit Protective Non-Neutralizing Antibodies that Target 11 Overlapping Regions on Viral Glycoprotein GP38

**DOI:** 10.1101/2024.03.02.583110

**Authors:** Olivia S. Shin, Stephanie R. Monticelli, Christy K. Hjorth, Vladlena Hornet, Michael Doyle, Dafna Abelson, Ana I. Kuehne, Albert Wang, Russell R. Bakken, Akaash Mishra, Marissa Middlecamp, Elizabeth Champney, Lauran Stuart, Daniel P. Maurer, Jiannan Li, Jacob Berrigan, Jennifer Barajas, Stephen Balinandi, Julius J. Lutwama, Leslie Lobel, Larry Zeitlin, Laura M. Walker, John M. Dye, Kartik Chandran, Andrew S. Herbert, Noel T. Pauli, Jason S. McLellan

**Author notes:** Deceased (Leslie Lobel). These authors contributed equally to the work (Olivia S. Shin, Stephanie R. Monticelli, Christy K. Hjorth). Correspondence (J.S.M.), (A.S.H.), (N.T.P.).

## Abstract

Crimean-Congo hemorrhagic fever virus can cause lethal disease in humans yet there are no approved medical countermeasures. Viral glycoprotein GP38, unique to *Nairoviridae*, is a target of protective antibodies, but extensive mapping of the human antibody response to GP38 has not been previously performed. Here, we isolated 188 GP38-specific antibodies from human survivors of infection. Competition experiments showed that these antibodies bind across five distinct antigenic sites, encompassing eleven overlapping regions. Additionally, we reveal structures of GP38 bound with nine of these antibodies targeting different antigenic sites. Although GP38-specific antibodies were non-neutralizing, several antibodies were found to have protection equal to or better than murine antibody 13G8 in two highly stringent rodent models of infection. Together, these data expand our understanding regarding this important viral protein and inform the development of broadly effective CCHFV antibody therapeutics.

## INTRODUCTION

Crimean-Congo hemorrhagic fever virus (CCHFV) is a member of the family *Nairoviridae* (*Orthonairovirus* genus) of the *Bunyavirales* order. Although infection by CCHFV is often asymptomatic in humans, severe hemorrhagic disease with fatality rates of 5–40%—and sometimes as high as 80%—have been documented^1,2^. Transmission of CCHFV to humans, as well as to domesticated and wild animals, occurs primarily through the bite of *Hyalomma* ticks^3–5^. Direct contact with infected tissues, primarily due to contact with blood from infected livestock, can also result in transmission^6,7^, and, though less common, nosocomial infections have been reported^8,9^. The broad geographic range of *Hyalomma* ticks contributes to widespread outbreaks of CCHFV across at least three continents, including Europe, Asia, and Africa, where CCHFV is endemic^2,6,10,11^.

Proportionate to its extensive distribution, CCHFV exhibits considerable genetic diversity among geographically distinct isolates^6,12^. Historically, CCHFV isolates were classified into six genotypes, or clades: I–III (endemic in Africa), IV (Asia), V (Europe I), and VI (Europe II)^12–19^. However, Clade VI genotypes were recently reclassified into a separate and distinct species, Aigai virus, which infrequently causes severe disease^20^. CCHFV has been recognized for its pandemic potential and, as of 2017, the World Health Organization has designated it a priority pathogen^21^. Despite this designation, no specific approved medical countermeasures are currently available, apart from the off-label use of the broad-spectrum antiviral ribavirin, but evidence for its efficacy against CCHFV is lacking^22^.

CCHFV has a tri-segmented negative-sense RNA genome. The genomic RNA segments are termed S (small), M (medium), and L (large), encoding for the nucleoprotein, the glycoprotein precursor complex (GPC), and the viral polymerase, respectively^23^. The GPC undergoes a series of proteolytic cleavages and maturation to generate multiple structural glycoproteins (Gc and Gn) and non-structural glycoproteins (GP38, GP85, GP160, and mucin-like domain)^24,25^. GP38 is unique to members of the *Nairoviridae* family and is thought to play a crucial role in CCHFV pathogenesis and the maturation of viral particles^26^. Crystal structures of CCHFV GP38 resolved in prior studies have shown the protein to have a novel fold consisting of an N-terminal 3-helix bundle followed by a β-sandwich^27,28^. Some evidence points to GP38 localizing to the membrane of virus particles and the surface of infected cells^29^. However, the specific function of GP38 and its role in pathogenesis remain unresolved.

Gc-specific neutralizing antibodies and GP38-specific non-neutralizing antibodies have been shown to be protective in animal models of infection^27–30^. 13G8, a non-neutralizing GP38-specific antibody of murine origin, has been characterized for its ability to protect mice against CCHFV-induced mortality and liver and spleen pathologies in both pre- and post-exposure studies^29^. Furthermore, 13G8 has shown varied prophylactic potential against diverse isolates of CCHFV, including IbAr10200, Afg09, and Turkey2004^27–29^. Investigations of the landscape of antibody responses to GP38 are limited, but two prior studies showed that antibodies target five discrete antigenic sites on CCHFV GP38^27,29^. These include seven human GP38-specific antibodies, one of which was structurally characterized and determined to compete with 13G8, but it was shown to be poorly protective compared to 13G8^27^. Given the unknown role of GP38 in viral pathogenesis and the limited understanding of epitopes contributing to protection, an evaluation of an extensive panel of human antibodies against GP38 is needed to investigate its function and develop effective antibody therapeutics.

Here, the B-cell repertoires of three human CCHF-convalescent donors from Uganda were mined for monoclonal antibodies specific for CCHFV GP38. A panel of 188 GP38-specific antibodies was isolated, binned into competition groups, and characterized for binding across several clinical isolates and for neutralization potency. Structural studies of select antibodies targeting each antigenic site were conducted to define epitopes across the surface of GP38. Subsequent animal challenge studies were performed to correlate protection with antigenic sites and gain insight into surfaces of GP38 that may be functionally important for pathogenesis.

## RESULTS

### Isolation of GP38-reactive antibodies from CCHF-convalescent donors

Peripheral blood mononuclear cells (PBMCs) were isolated from three human CCHF-convalescent donors from Uganda between 3- and 46-months post-infection (**Table 1**). All donors had detectable serum titers to GP38, relative to naïve controls (**Supplementary Figure S1A**). To sort memory B cells (MBCs) expressing GP38-reactive B cell receptors, PBMCs were stained with fluorescently conjugated recombinant IbAr10200 GP38 (rGP38), expressed from a stably transfected Schneider 2 cell line, and a panel of fluorescently conjugated antibodies to cell-surface markers.

**Table 1.** Patient metadata of CCHF-convalescent donors. Date of infection, blood donation, and hospitalization time are approximate.

| Donor ID | Sex | Birth Year | Date of Infection | Date of Blood Donation | Time Post-Infection (months) | Diagnostic Method | Hospitalization Time (days) | District |
| --- | --- | --- | --- | --- | --- | --- | --- | --- |
| 1 | M | 1949 | Aug 2013 | Jun 2017 | 46 | PCR | 14 | Agago |
| 5 | M | 1982 | Nov 2015 | Nov 2017 | 24 | PCR | 14 | Nakaseke |
| 6 | M | 1987 | Aug 2017 | Nov 2017 | 3 | PCR | >28 | Nakaseke |

Of the total population of switched immunoglobulin (SwIg) B cells, 0.35%, 0.14%, and 0.11% were rGP38-reactive for donors 1, 5, and 6, respectively (**Supplementary Figure S1B**). Flow analysis demonstrated that 63–91% of GP38-reactive B cells from these donors were class-switched (**Figure 1A**), indicative of a MBC response and of class-switch recombination dynamics consistent with the Gc-specific CCHFV response^30^. Of this class-switched, GP38-reactive population, 50.0%, 15.8%, and 25.0% of the cells were CD27^+^ for Donors 1, 5, and 6, respectively (**Supplementary Figure S1C** and **D**), consistent with the varying levels of CD27 expression observed in the human MBC compartment^31–34^. Because the majority of the GP38-reactive B cells were IgM^−^ IgD^−^, only these SwIg B cells were isolated for further downstream analysis (**Figure 1A** and **Supplementary Figure S1E**). Isolated antibody genes from sorted B cells were amplified using V_H_ and V_κ_ or V_λ_ single-cell PCR. In total, 254 paired V_H_/V_L_ antibody genes were successfully cloned into an IgG1 isotype in a proprietary, engineered *S. cerevisiae* strain.

**Figure 1.**
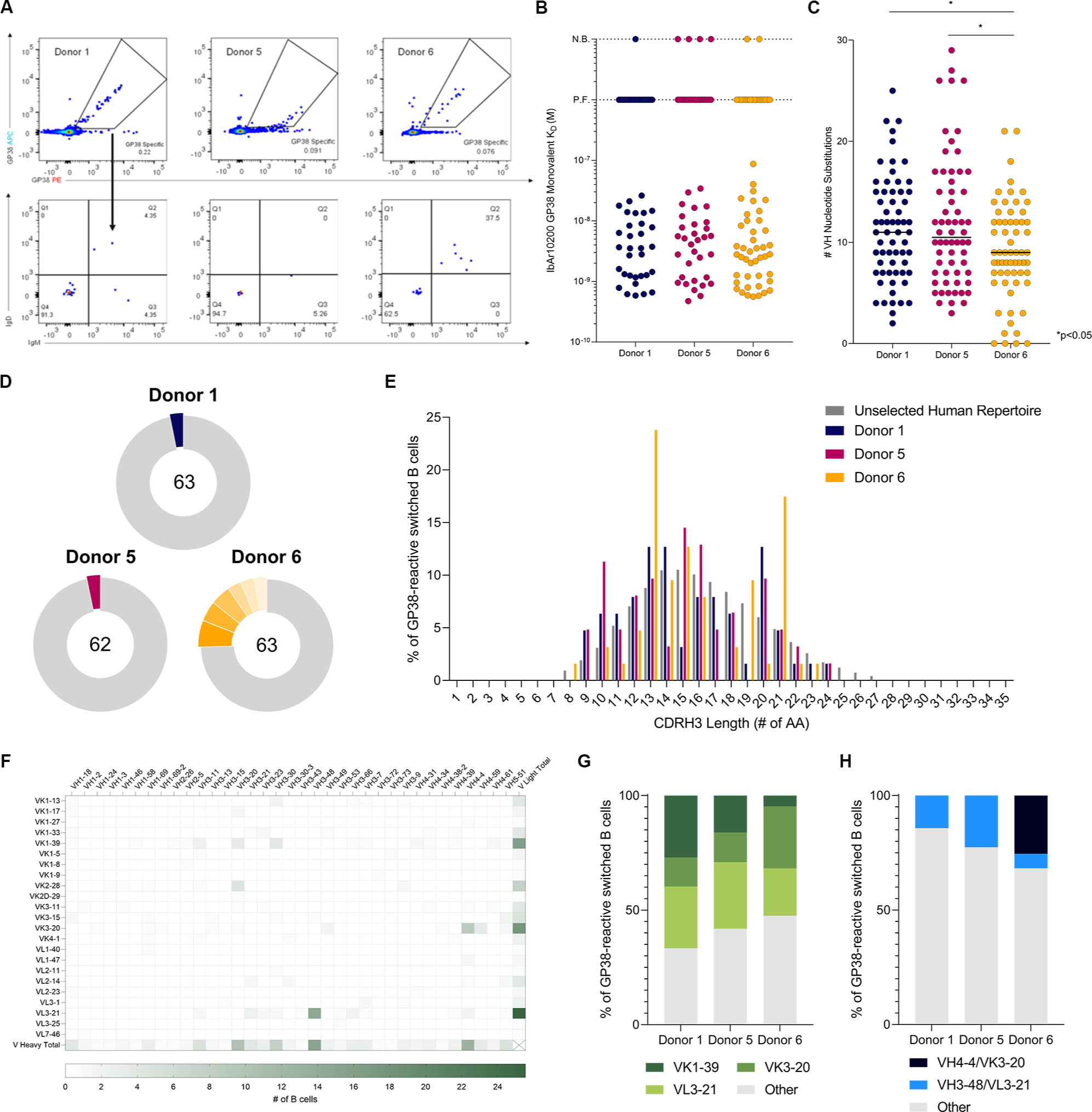
Isolation of monoclonal antibodies and genetic signatures of the B cell repertoire. (A) Flow cytometric analysis of avid-rGP38 binding of B cells (top panel) and IgM and IgD expression on the surface of rGP38-reactive B cells (bottom panel). Donor 1 PBMCs were gated on CD3^−^CD8^−^CD14^−^CD16^−^PI^−^CD19^+^ lymphocytes; Donor 5 and 6 PMBCs were gated on CD3^−^CD8^−^CD14^−^CD16^−^PI^−^CD19^+^CD20^+^ lymphocytes. (B) Single concentration BLI binding analysis of 188 antibodies to IbAr10200 rGP38 protein. Dotted horizontal lines indicate antibodies for which no binding (N.B.) was detected or for which poor fits (P.F.) to the binding model were obtained (C) Analysis of VH nucleotide substitutions of each of the mAbs. Statistical comparison was performed using the Mann-Whitney Test (*p<0.05). (D) Clonal lineage analysis of B cells from Donors 1, 5, and 6. B cells with antibody sequences that had the same V heavy and V light germline gene usage and CDRH3s of the same length with >80% nucleotide sequence identity were considered to be clonally related. Colored slices represent the percentage of clones from each donor that are related. Total number of isolated mAbs from each donor are indicated in each corresponding circular diagram. (E) Analysis of CDRH3 lengths of mAbs from the three donors. (F) Heatmap of VH and VL germline gene usage across mAbs from the three donors; shades of green represent the number of B cells that used a certain germline gene pairing. (G) Analysis of VK1-29, VK3-20, and VL3-21 germline gene usage broken down by donor. (H) Analysis of VH4-4/VK3-20 and VH3-48/VL3-21 germline gene usage broken down by donor.

After expression and purification of this panel of monoclonal antibodies (mAbs), we assessed binding of the full-length IgGs to IbAr10200 rGP38 using biolayer interferometry (BLI). We found that 188 of the 254 purified mAbs bound to rGP38 in this assay (**Supplementary Figure S2A** and **B**). To better understand the human immune response against GP38, we determined the affinities of these antigen-specific IgGs via BLI. 181 mAbs had detectable monovalent binding to IbAr10200 rGP38 (**Figure 1B**). Of the 107 monovalent binders for which a 1:1 binding model could be fit, 78.5% (n=84) had affinities better than 10 nM (**Figure 1B** and **Supplementary Figure S2C**). Antibodies isolated from Donors 1, 5, and 6, displayed single-digit nanomolar median binding affinities against IbAr10200 rGP38 with median affinities of 3.5 nM, 4.4 nM, and 2.8 nM, respectively (**Supplementary Figure 2C**). Taken together, these data indicate that convalescent CCHFV-infected donors can generate high-affinity, long-lived, GP38-specific antibody responses.

### Genetic signatures of GP38-specific antibodies

We next assessed the specific genetic signatures associated with CCHF-convalescent donor antibody responses to GP38. Previous work has described CCHFV Gc-specific antibody responses, as well as genetic signatures typically observed in antibodies elicited by other primary viral infections or vaccinations^30,35–37^. Somatic hypermutation (SHM)—a hallmark of affinity maturation—and clonal diversity are important metrics in the assessment of the quality of an antigen-specific antibody response following infection or immunization^38,39^. Antibodies from the three donors had median values of SHM between 9 and 11 heavy-chain nucleotide substitutions (**Figure 1C**), and in general, samples collected from donors with longer times between infection and blood donation contained B cells with higher levels of SHM (**Table 1** and **Figure 1C**). Paired heavy- and light-chain analyses demonstrated high levels of clonal diversity (3–25% clonal relatedness) amongst antibodies cloned from all three donors (**Figure 1D**), similar to levels of diversity seen amongst B cells isolated from survivors of Ebola virus and SARS-CoV-2 infections^34,36^. Interestingly, the higher clonal relatedness (25%) amongst GP38-reactive B cells cloned from Donor 6 is in contrast with what was seen amongst Gc-specific MBCs (0% clonal relatedness) from the same donor^30^. GP38-specific mAbs from all three donors had a similar distribution of heavy-chain complementarity-determining region three (CDRH3) lengths as compared to the unselected human repertoire^40^ (**Figure 1E**). However, the Donor 6 B cell response appears to be skewed toward clones with CDRH3 lengths of 13 and 21 amino acids, consistent with data showing that most of these clones arose from two distinct clonal expansions (**Supplementary Figure S3**).

We next sought to determine if specific V-genes were preferentially enriched in GP38 antibodies collected from these donors. Across all donors, sorted GP38-reactive B cells utilized V_K_1-39, V_K_3-20, and V_L_3-21 light chain V-genes most often, at a frequency of 16%, 17%, and 26%, respectively (**Figure 1F** and **Supplementary Figure S4**). For each individual donor, greater than 50% of all sorted GP38-reactive B cells utilized these three light chain V-genes (**Figure 1G** and **Supplementary Figure S4**). Heavy chain V-gene usage was less skewed than light chain V-gene usage, however, 13% of all cloned GP38-specific antibodies used V_H_3-48 and 15% used V_H_4-4 V-genes (**Figure 1F** and **Supplementary Figure S4**). V_H_3-48 predominantly paired with V_L_3-21 and V_H_4-4 paired with V_K_3-20 (**Figure 1F**). Although the V_H_3-48/V_L_3-21 pairing was seen across all donors, the V_H_4-4/V_K_3-20 pairing was a unique feature of the Donor 6 response (**Figure 1H**). Collectively, our analysis shows that this isolated panel of GP38-specific antibodies is derived from a diverse population of B cells with a preference toward specific heavy and light chain V-genes.

### GP38-specific antibodies recognize 11 overlapping antigenic regions

We conducted binding-competition assays to better understand where on GP38 the isolated antibodies bound. Because we lacked the capacity to cross-bin 188 mAbs (i.e., a 188 x 188 matrix), we down-selected our repertoire to 19 high-affinity clones with disparate V_H_/V_L_ germline pairings and CDRH3 sequences to perform multiple cross-competition experiments (**Supplementary Figure S5**). From these experiments, we discovered seven high-affinity mAbs (ADI-46120, ADI-46146, ADI-46152, ADI-46158, ADI-46172, ADI-46174, and ADI-58048) that, when cross-binned in yeast-based competition assays, revealed the presence of five-non-overlapping bins (**Figure 2A**), as has been described previously^27^.

**Figure 2.**
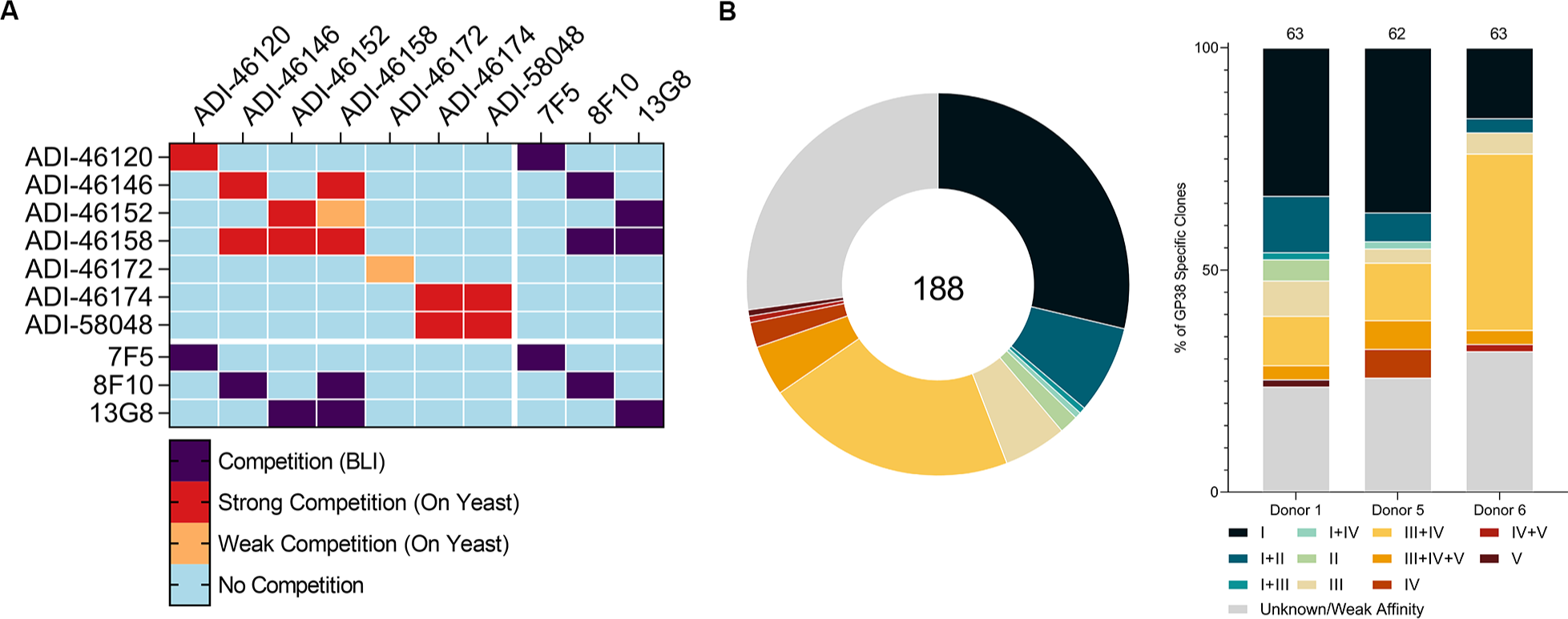
Competition-binning profile of GP38 antibodies. (A) Matrix of competition-binning experiments. For on-yeast competition experiments (top left quadrant), results are displayed with surface-presented IgGs on the y-axis and competitive pre-complexed Fabs on the x-axis. For BLI competition assays (the other three quadrants), binning was performed in an IgG vs. IgG format. (B) Binning analysis of on-yeast competition assays of all 188 antibodies; each color represents one of 11 overlapping bins and the Unknown/Weak Affinity mAbs are shown in gray (**Supplementary Table 1**). Distribution of overlapping bins of the antibody panel (left) and broken down by donor (right). Total number of mAbs is indicated in the circular diagram and total mAbs from each donor are indicated above the bar graph.

To gain a more granular understanding of the immunogenic surface of GP38, we performed a binning assay with our entire panel of 188 GP38-specific antibodies. We chose one antibody from each of the five non-overlapping antigenic sites to be run in competition against all 188 antibodies (i.e., a 188 x 5 matrix): ADI-46120, ADI-46146, ADI-46152, ADI-46158, and ADI-58048. The highest affinity antibody from each of the five non-overlapping bins was selected to provide the assay with the greatest discriminatory power. The results revealed that our panel of 188 GP38 mAbs fell into 11 overlapping bins (**Figure 2B**). Antibodies that only competed with one of the five representative antibodies were labeled as bin I (ADI-46120 competitor), II (ADI-58048 competitor), III (ADI-46146 competitor), IV (ADI-46158 competitor), or V (ADI-46152 competitor) and antibodies that competed with one or more of the five representative antibodies were labeled with two or more roman numerals (i.e., bin III+IV antibodies compete with both ADI-46146 and ADI-46158) (**Supplementary Table S1**). Across all donors, the immune response consisted primarily of antibodies from bin I (n=54) and bin III+IV (n=40) (**Figure 2B**). Fifty of the 188 antibodies (26.6%) did not appear to compete with any of the five selected competitor antibodies (**Figure 2B**). Many of these antibodies likely appear non-competitive in yeast-based competition assays because of their weak affinity for IbAr10200 GP38; however, a subset did bind to IbAr10200 GP38 and may recognize unique antigenic sites (**Supplementary Figure S6**). We also conducted cross-competition assays with three previously characterized murine mAbs (7F5, 8F10, and 13G8). These experiments revealed that 7F5 is a bin I mAb as it competes with ADI-46120, 8F10 is a bin III+IV mAb as it competes with both ADI-46146 and ADI-46158, and 13G8 is a bin IV+V mAb as it competes with both ADI-46158 and ADI-46152 (**Figure 2A**). Collectively, these studies identify 11 overlapping regions on the GP38 surface targeted by human and murine antibodies.

### GP38-specific antibodies are broadly reactive

Our initial binding studies used GP38 derived from CCHFV IbAr10200 (**Figure 1** and **Supplementary Figure S2**), a clade III virus. However, this is a highly laboratory-passaged virus with little clinical relevance. Most confirmed reported cases of human infection are attributed to isolates from clades III, IV (Afg09, Oman, and China), and V (Turkey2004 and Hoti)^1,6,18,41^, and over the past few years, new strains have emerged from areas where these clades are endemic^42,43^. Therefore, we chose five clinically relevant isolates (Afg09, Turkey2004, Oman, Hoti, and M18-China) in addition to IbAr10200 to determine the extent to which the 188 GP38-specific antibodies bind to multiple clinically relevant and diverse isolates. The GP38s of the aforementioned CCHFV isolates exhibit between 70–92% amino acid sequence similarity with IbAr10200 (**Figure 3A**). Sequence alignment of the six isolates reveals that much of the variation occurs in variable loops 1 (residues 322–341) and 2 (residues 377–394) (**Supplementary Figure S7**). First, we used BLI to assess the monovalent affinity of each of the 188 mAbs at a single concentration to each of the six GP38 variants. mAbs for which the recorded response was greater than 0.05 response units (RUs) were considered to bind to the respective rGP38 protein. These experiments revealed that 87% of the 188 GP38-specific mAbs bound GP38 derived from all six tested isolates and 8% across five of six; the remaining 5% of mAbs bound GP38 derived from 4 or fewer isolates (**Figure 3B**). These high levels of cross-reactivity are comparable to those seen in the Gc-specific responses from the same donors^30^. The single-concentration BLI data were used to select high-affinity, cross-reactive clones with varying germline usage from discrete bins (**Supplementary Table S2**). Antibody-drug developability metrics (i.e. polyreactivity, hydrophobic interaction chromatography, thermostability; **Supplementary Table S3**)^44^ were then run on these clones of interest and lead candidates were established for further study: ADI-58026 (bin I), ADI-58062 (bin I+II), ADI-58048 (bin II), ADI-63530 (bin III+IV), ADI-46138 (bin III+IV+V) and ADI-63547 (bin IV+V).

**Figure 3.**
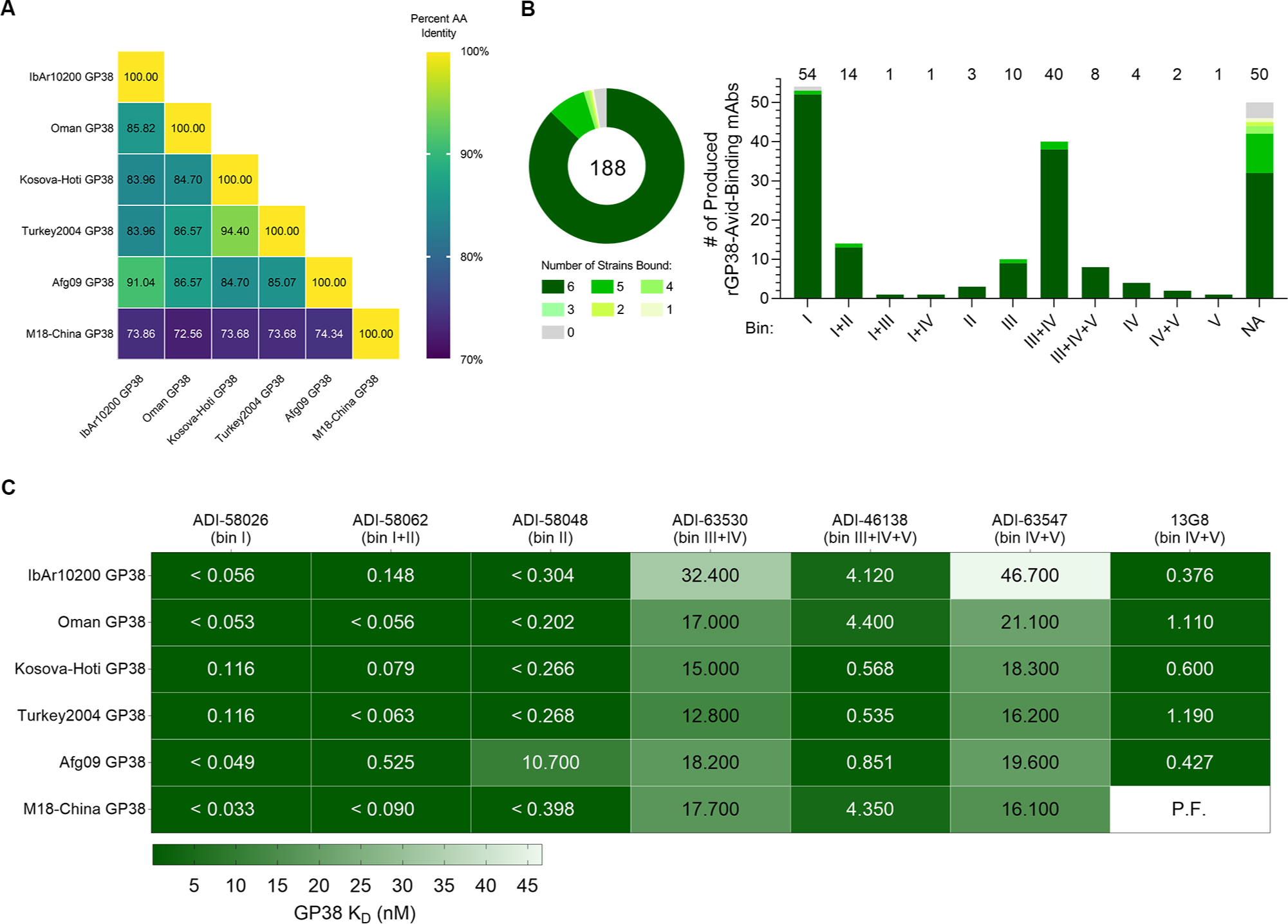
Analysis of antibody binding to GP38 proteins derived from six CCHFV isolates. (A) Matrix of percent sequence identity of GP38 amino acid residues across six CCHFV isolates. (B) Single concentration BLI binding analysis of 188 antibodies to the six rGP38 proteins as a whole panel (left) and broken down by bin (right). Shades of green represent the number of rGP38 proteins bound by a single antibody. Total number of mAbs is indicated in the circular diagram and total mAbs from each donor are indicated above the bar graph. (C) Carterra system HT-SPR binding analysis of six lead antibody candidates binding to six rGP38 proteins. The highest apparent binding affinities are in dark green and the lowest apparent binding affinities are in white. Calculated K_D_s appear in each rectangle of the heat map; for samples that were off-rate limited, K_D_s are denoted as < the calculated K_D_. The one interaction for which a curve could not be fit is denoted as P.F.

To gain a more nuanced understanding of the cross-clade binding dynamics, we used the Carterra system to carry out multipoint *K*_D_ measurements for the six lead antibodies as well as the previously described murine mAb 13G8. ADI-58026 (bin I) and ADI-58062 (bin I+II) bound to all six GP38 variants derived from clinical isolates with affinities better than 530 pM, and ADI-58048 (bin II) bound with an affinity less than 398 pM to five of six GP38 variants but had an approximately 27-fold reduction in binding to Afg09-derived GP38 (**Figure 3C** and **Supplementary Table S4**). Each mAb from bins III–V (ADI-63530, ADI-46138, and ADI-63547) bound to the six tested GP38 variants with affinities of 12.8–32.4 nM, 0.54–4.4 nM, and 16.2–46.7 nM, respectively (**Figure 3C** and **Supplementary Table S4**). These three mAbs all bound the six GP38 variants with affinities that were within 10-fold of their affinity to IbAr10200 GP38. Of these three antibodies, ADI-46138 exhibited the highest binding affinities, which were 3- to 30-fold higher than those determined for ADI-63530 and ADI-63547 (**Figure 3C** and **Supplementary Table S4**). Additionally, ADI-46138 (bin III+IV+V) and 13G8 (bin IV+V) bound to five GP38 variants with affinities within 11-fold of one another (**Figure 3C** and **Supplementary Table S4**). Taken together, 95% of the 188 isolated GP38-specific antibodies bound to five or six GP38 variants derived from clinically relevant CCHFV isolates spanning diverse clades, and antibodies ADI-58026 (bin I) and ADI-58062 (bin I+II) bound these GP38 variants with picomolar affinities.

### Antibodies targeting GP38 are non-neutralizing

The six lead GP38-specific mAbs were tested in a microneutralization assay utilizing transcription- and entry-competent virus-like particles (tecVLPs) bearing IbAr10200 GPC-derived proteins^30,45^. None of the GP38-specific antibodies neutralized the tecVLPs in this assay (**Figure 4A**). Neutralization assays were also performed with authentic CCHFV, including the prototype IbAr10200 (clade III; **Figure 4B**) and clinically relevant isolates Afg09 (clade IV; **Figure 4C**), Turkey2004 (clade V; **Figure 4D**), and Oman (clade IV; **Figure 4E**) in SW-13 cells, a cell line relevant for CCHFV-infection that exhibits epithelial morphology^46^. Again, none of the GP38-specific mAbs exhibited significant neutralization potency against the tested authentic viruses (**Figure 4**), consistent with previous reports^27,29,30,47^. To determine whether neutralization potency was cell-type specific, a microneutralization assay was also conducted in VeroE6 cells with authentic viruses. Comparable to the results obtained in SW-13 cells, none of the GP38 mAbs afforded significant neutralization potency against any of the CCHFV isolates tested in VeroE6 cells (**Supplementary Figure S8**). ADI-36121, a Gc-specific monoclonal antibody previously shown to afford significant cross-clade neutralization efficacy against CCHFV^30^, was utilized as a positive control and, as anticipated, potently neutralized tecVLPs (**Figure 4A**) and all isolates of authentic CCHFV tested in both SW-13 (**Figure 4B-E**) and VeroE6 cells (**Supplementary Figure S8**). Consistent with previously reported studies, our panel of GP38-specific antibodies was non-neutralizing under the conditions tested^27,29,30^.

**Figure 4.**
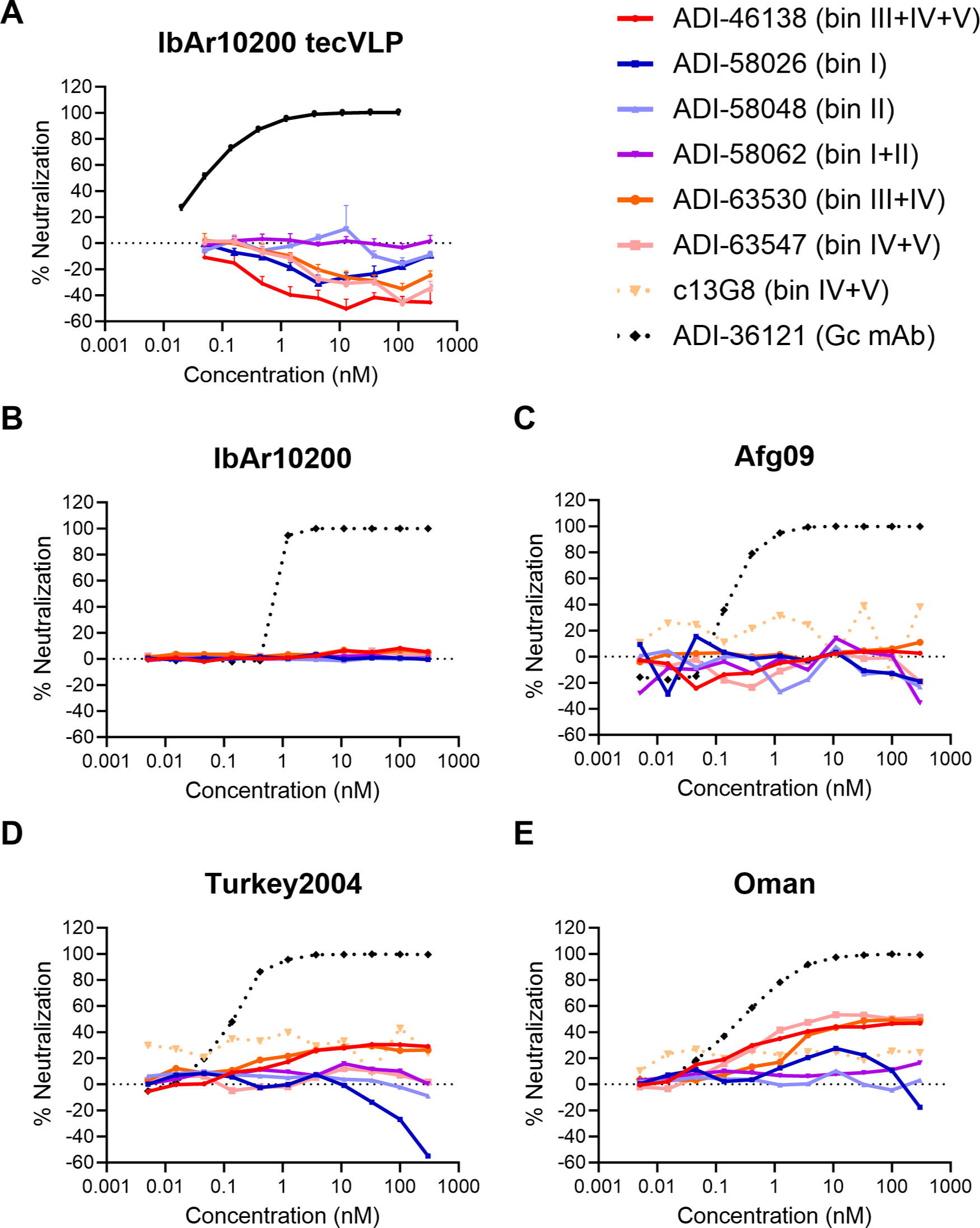
CCHFV tecVLPs and authentic virus neutralization assays of GP38 antibodies. (A) Individual neutralization curves for CCHFV IbAr10200 tecVLPs, as measured by the reduction in luciferase activity compared to no-antibody treatment on Vero cells. (B-E) Neutralization curves of the indicated mAbs against authentic (B) CCHFV IbAr10200, (C) CCHFV Afg09, (D) CCHFV Turkey2004, and (E) CCHFV Oman as measured by the reduction in infection compared to no-antibody treatment on SW-13 cells. The average of n=3 replicates each from two independent experiments (n=6 total) is shown for all neutralization curves.

### Epitope mapping reveals two predominantly targeted regions on GP38

We set out to map the location of the antigenic sites on GP38 to correlate certain epitopes with protection and function. We employed a yeast surface display (YSD)-based mapping and structural characterization strategy utilizing select GP38 antibodies. A YSD library of GP38 single-amino-acid variants was generated to compare antibody binding between mutant and wild-type GP38. Nine antibodies representing seven of the eleven overlapping bins successfully underwent YSD mapping to reveal critical residues on GP38 necessary for retaining antibody binding (**Figure 5A**). Critical residues that disrupted antibody binding by 75% or more were mapped onto the surface of IbAr10200 GP38 (PDB ID: 6VKF) to represent the five discrete antigenic sites (**Figure 5B** and **Supplementary Figure S9**). These studies were complemented with structural studies of select antibodies to further characterize the antigenic sites.

**Figure 5.**
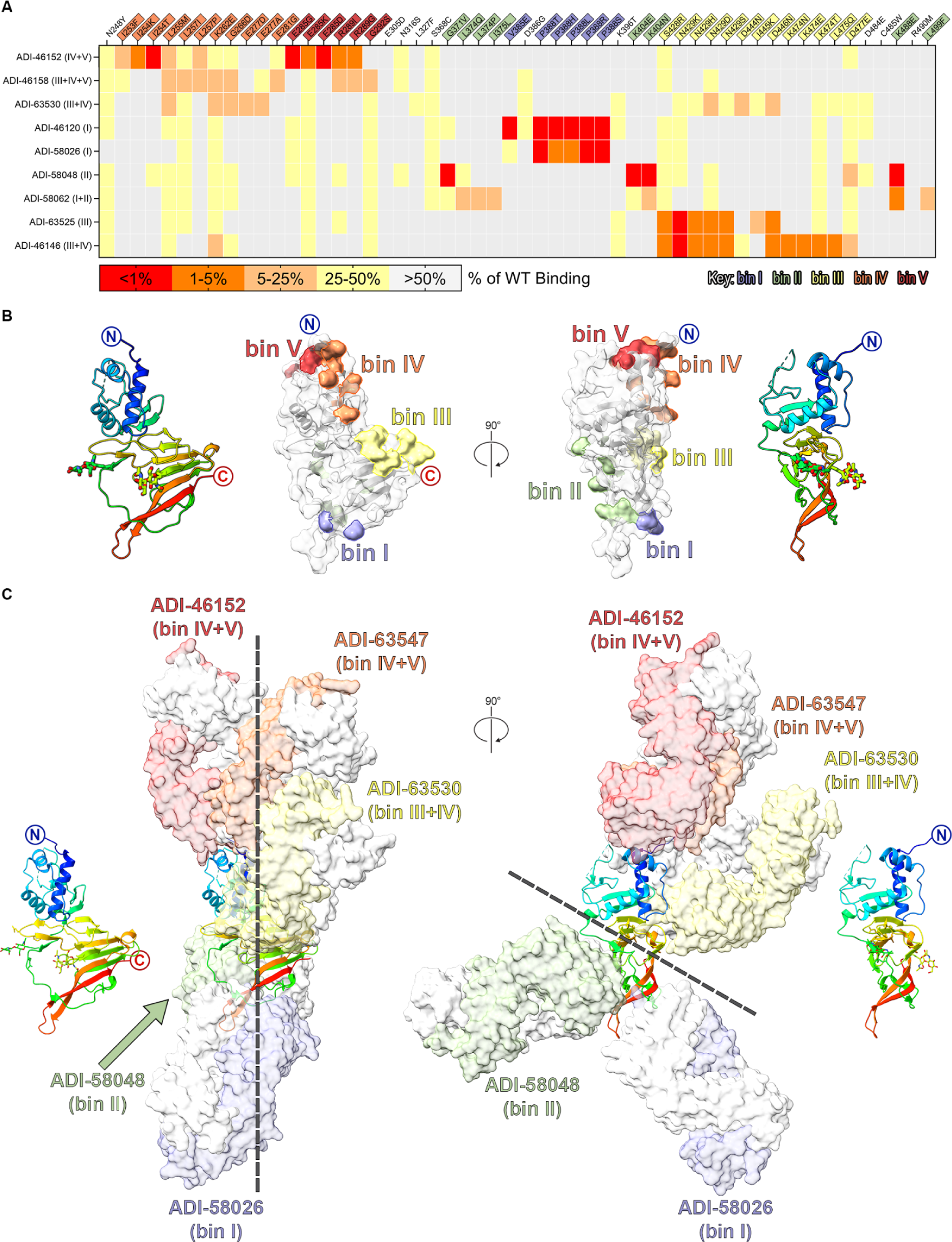
Structural characterization of GP38 antibodies. (A) Yeast-based mapping strategy of select antibodies to identify critical binding residues on GP38. The percentage of antibody binding retained by each GP38 variant is colored according to the key. Critical residues are defined as mutations that led to a binding disruption of 75% or more and are colored by the assigned antigenic site. (B) Yeast-based critical residues mapped on the surface of GP38 (PDB ID: 6VKF): bin I (blue, residues Val385, Pro388), bin II (green, residues Gly371, Leu374, Ile375, Lys404, Lys488, Leu499), bin III (yellow, residues Ser428–Ala429, Asp444–Asp446, Lys474– Leu475, Asp477), bin IV (orange, residues Ile253–Leu255, Leu257, Lys262, Gly266, Glu277, and Glu281), bin V (red, residues Glu285, Arg289, and Gly292). (C) Composite structure of GP38 (PDB ID: 6VKF) bound with representative antibodies. GP38 is shown as a rainbow ribbon, and Fabs as molecular surfaces. Heavy chains are colored to represent the 5 non-overlapping bins, and light chains are white. Black dashed lines highlight the vertical alignment of Fabs along one plane (left) and the opposing binding directions to another plane (right).

To map the epitope of bin I antibodies, we determined a 5.0 Å resolution cryo-EM structure of ADI-58026 Fab (bin I) and ADI-63547 Fab (bin IV+V) bound to GP38 (**Supplementary Figure S10** and **Supplementary Table 5**). Due to the resolution of the cryo-EM reconstruction, we docked AlphaFold2 models of the Fabs into the maps to assess the epitopes. The docked ADI-58026 Fab binds near the second variable loop and C-terminal β-hairpin, in excellent agreement with bin I YSD critical residues Val385 and Pro388 (**Figure 5C**). To further characterize the bin I epitope, we complexed ADI-46143 Fab (bin I) to GP38 and determined a 2.6 Å resolution crystal structure (R_work_/R_free_ = 0.177/0.217), which revealed that ADI-46143 Fab binds primarily to the second variable loop, with additional contacts to the C-terminal β12-β13 hairpin, similar to ADI-58026 (**Figure 6A**, **Supplementary Figures S10**, **Supplementary Table S6**). Pro388—a YSD-identified critical residue of bin I antibodies—is at the center of the ADI-46143 epitope (**Figure 6A**).

**Figure 6.**
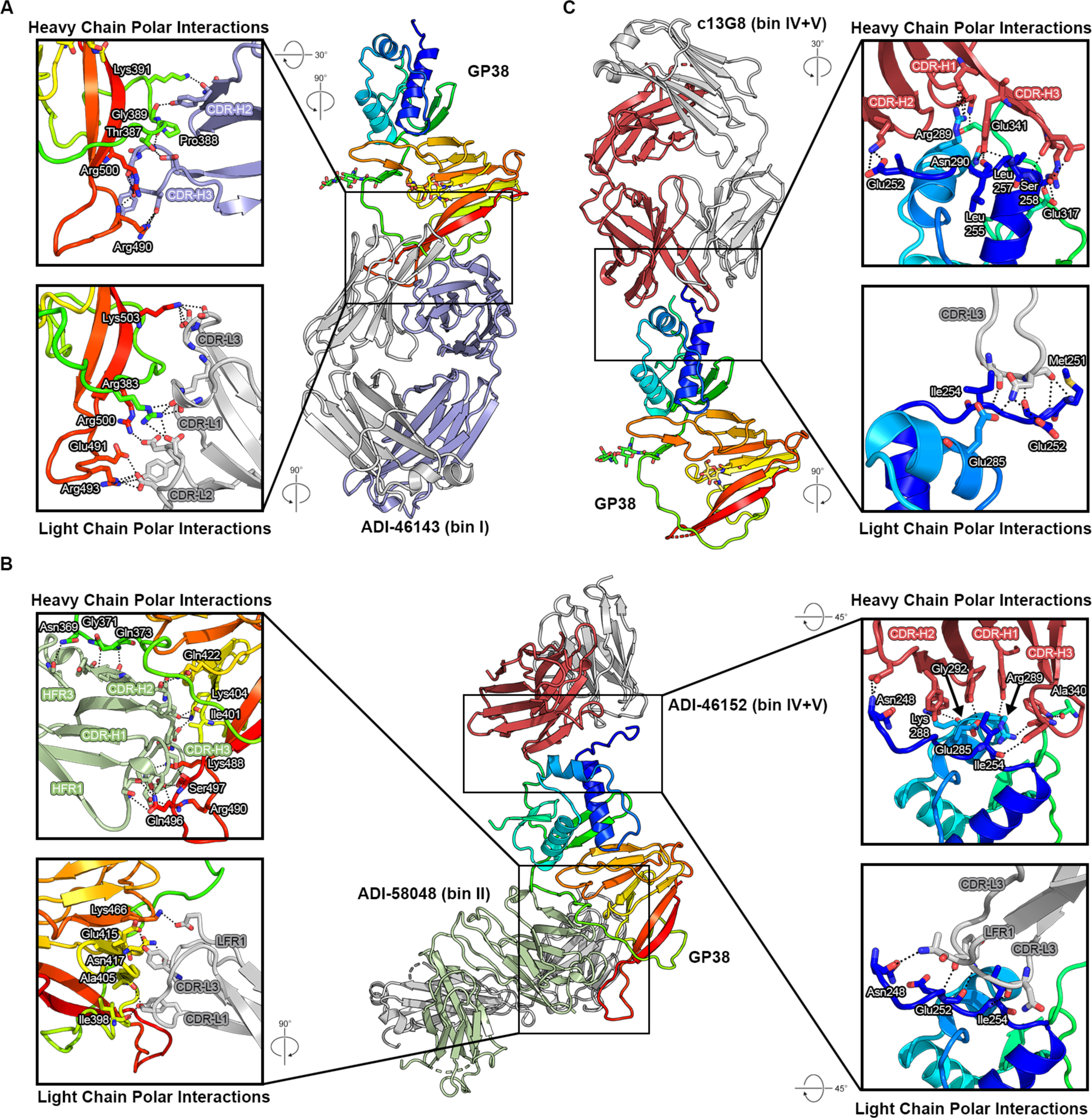
High-resolution structures of GP38–antibody complexes. (A) Crystal structure of GP38 bound with ADI-46143 (bin I, blue) with heavy chain interactions (top) and light chain interactions (bottom). (B) Cryo-EM structure of GP38 bound with ADI-58048 (bin II, green, left) and ADI-46152 (bin IV+V, red, right). Heavy chain interactions (top left, top right) and light chain interactions (bottom left, bottom right) are shown in the insets. (C) Crystal structure of GP38 bound with c13G8 (bin IV+V, red) with heavy chain interactions (top) and light chain interactions (bottom). For all panels, hydrogen bonds are indicated by black dashed lines. GP38 residues are labeled in white text with a black outline.

To map the epitope of bin II antibodies, a complex of GP38 bound with ADI-58048 Fab (bin II) and ADI-46152 Fab (bin IV+V) was generated and a 3.8 Å resolution cryo-EM structure of the complex was determined (**Figure 6B**, **Supplementary Figure S11** and **Supplementary Table S5**). ADI-58048 binds the β-sandwich, including residues in the long loop connecting the C-terminal β12-β13 hairpin (**Figures 5C** and **6B**, **Supplementary Figure S11**). Three bin II YSD critical residues on GP38 (Gly371, Lys404, Lys488) are at the interface with the ADI-58048 heavy chain (**Figure 6B**). Lys404 and Lys488 are at the interface with the ADI-58048 CDRs, whereas Gly371 is located in the first variable loop of GP38 and rests against the side of the V_H_ domain. We also determined a 5.1 Å resolution cryo-EM structure of GP38 in complex with ADI-58062 Fab (bin I+II) and ADI-63530 Fab (bin III+IV), which revealed that ADI-58062 binds to a similar epitope as ADI-58048 (**Supplementary Figures S11** and **S12**), and the antibodies would sterically clash, as expected for two bin II competitors.

To map the epitopes of antibodies that competed across bins III–V, we analyzed the aforementioned cryo-EM structures as well as determined a 5.8 Å resolution cryo-EM of GP38 in complex with ADI-46158 (bin III+IV+V) and ADI-46143 (bin I) (**Figure 5C**, **Supplementary Figures S10–13**). Consistent with bin IV critical residues, ADI-46158 and ADI-63547 (bin IV+V) Fabs bind the 3-helix bundle, primarily the first several N-terminal residues and the beginning of α-helix 1 (**Figure 5C**, **Supplementary Figures S10** and **S13**). Their epitopes predominantly target bin IV YSD critical residues while also contacting bin V YSD critical residues Glu285 and Arg289 on α-helix 2. The ADI-63530 (bin III+IV) epitope spans both the 3-helix bundle and β-sandwich, consistent with bin III YSD critical residues Ser428–Ala429, Asp444–Asp446, Lys474–Leu475, and Asp477, which are in loops connecting strands β6-β7, β8-β9, and β11-β12 (**Figure 5C** and **Supplementary Figure S12**).

To further map the epitope of bin V antibodies, we selected two bin IV+V antibodies for structural studies: ADI-46152 and a humanized chimeric variant of 13G8 (c13G8)^28^. From the 3.8 Å resolution cryo-EM structure of GP38 bound with ADI-58048 Fab (bin II) and ADI-46152 Fab, the ADI-46152 heavy chain makes several contacts on α-helix 2, N-terminal residues preceding α-helix 1, and variable loop 1 while the ADI-46152 light chain contacts N-terminal residues Asn248, Glu252, and Ile254, consistent with bin IV and V YSD residues (**Figures 5C** and **6B**, **Supplementary Figure S11**, and **Supplementary Table S5**). To resolve the epitope of 13G8, c13G8 Fab was complexed to GP38, and we determined a 1.8 Å resolution crystal structure (R_work_/R_free_ = 0.200/0.215) (**Figure 6C** and **Supplementary Table S6**). The structure revealed that c13G8 binds to the N-terminal 3-helix bundle of GP38, consistent with the 3.6 Å structure determined by^27^. YSD critical residues identified on GP38 (Ser258, Arg289, and Asn290) interact with the c13G8 heavy chain and YSD critical residue Ile254 is also at the antibody interface (**Figure 6C**). Epitopes of c13G8 and ADI-46152 are highly overlapping and share two YSD critical residues (Ile254 and Arg289) (**Supplementary Figure S14**). Compared to ADI-63547 (bin IV+V), ADI-46152 and c13G8 have shifted angles of approach that extend contacts to residues Glu317 and Ala340 within the bin V epitope.

To visualize the overall antigenic landscape, we generated a composite view of GP38 bound with Fabs ADI-58026 (bin I), ADI-58048 (bin II), ADI-63530 (bin III+IV), ADI-63547 (bin IV+V) and ADI-46152 (bin IV+V) (**Figure 5C**). These antibodies are representative of the five antigenic sites based on both YSD-based mapping and structural studies. The composite structure reveals that the antibodies approach GP38 along a similar plane. Furthermore, the antibodies bind predominately to two general regions: an N-terminal region containing bins III–V comprising the 3-helix bundle and loops connecting adjacent β-strands, and a region containing bins I and II comprising the second variable loop and C-terminal β-hairpin. These restricted binding modes may result in part from how GP38 is oriented on the virion or in complex with other proteins from the GPC.

### Antibodies targeting epitope bins III, IV, and V afford partial therapeutic protection against a lethal CCHFV-IbAr10200 challenge

We next evaluated the therapeutic potential our six lead GP38-specific antibodies in an immunocompromised rodent model of lethal CCHFV challenge: ADI-58026 (bin I), ADI-58048 (bin II), ADI-58062 (bin I+II), ADI-63530 (bin III+IV), ADI-63547 (bin IV+V), and ADI-46138 (bin III+IV+V). c13G8 (bin IV+V) was included as a benchmark for comparison to previously published studies. Type I interferon ɑ/β R^−/−^ (IFNAR1^−/−^) mice^48,49^ were challenged with 100 PFU of CCHFV-IbAr10200 and subsequently treated with 1 mg of mAb per animal 1- and 4-days post-challenge (2 mg/mouse total), to replicate previous conditions testing 13G8 efficacy^27,29^. As previously described, c13G8 afforded partial protection (40%) (**Figure 7A-C**). Antibodies targeting GP38 epitope bins I (ADI-58026), II (ADI-58048), or I+II (ADI-58062) were minimally protective (20-30% survival), and less so than that of c13G8. In contrast, antibodies targeting epitope bins III+IV+V (ADI-46138) and IV+V (ADI-63547) were similarly protective as c13G8 (40% survival). Moreover, antibody ADI-63530, targeting GP38 epitope bins III+IV, exhibited substantial protection (70%), which was greater than that observed for c13G8. Collectively, these data indicate that antibodies targeting GP38 epitope bins I and II are minimally protective, whereas antibodies targeting GP38 epitope bins III, IV, and V are most protective against a CCHFV-IbAr10200 lethal challenge. Interestingly, although antibodies targeting GP38 epitope bins III, IV, and V were more protective than the bin I and II targeting antibodies, the bin III, IV, and V specific antibodies displayed lower affinities compared to the bin I and II specific antibodies (**Figure 3** and **Supplementary Figure S15**). These findings indicate that human monoclonal antibodies targeting bins III, IV, and V on GP38 are equally, if not more efficacious than, the previously described murine mAb 13G8 against a lethal CCHFV-IbAr10200 challenge.

**Figure 7.**
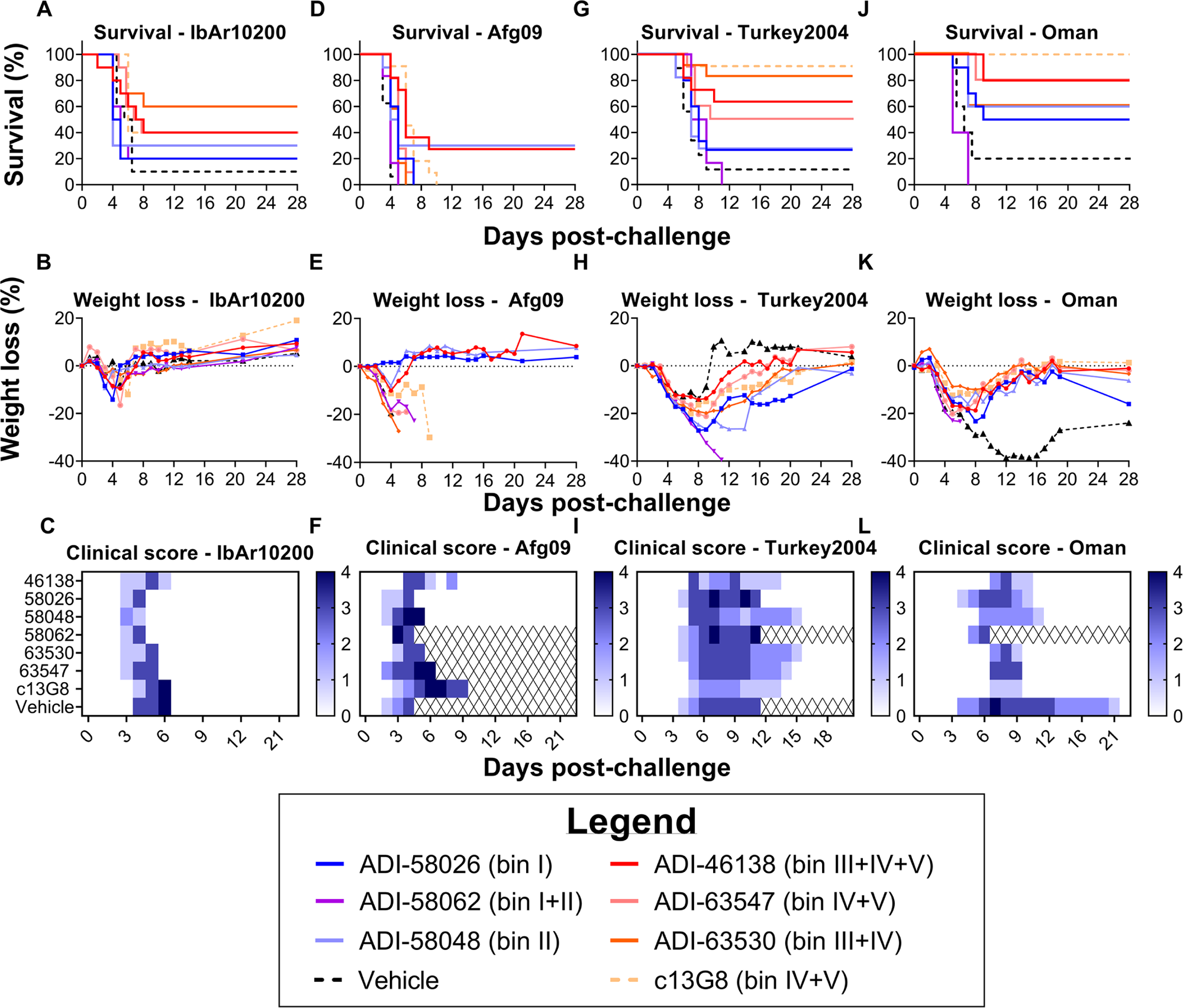
Protective efficacy of lead mAbs in two murine models of lethal CCHFV challenge. (A-C) IFNAR1^−/−^ mice were treated with the indicated mAbs at 1 mg/mouse 1- and 4-days post-challenge (2 mg total; n=10 mice per group). (A) Survival curves (vehicle versus test mAb), (B) associated mean weight loss, and (C) clinical score data are shown. (D-L) STAT1^−/−^ mice were challenged with (D-F) CCHFV-Afg09, (G-I) CCHFV-Turkey2004, or (J-L) CCHFV-Oman and then treated with 0.2 mg/mouse of mAb or vehicle 30 minutes post-exposure (n=5–6 mice per study; represented by 2 replicate studies). (D, G, J) Survival curves, (E, H, K) associated mean weight loss, and (F, I, L) clinical scores.

### ADI-46138 and ADI-58048 provide cross-clade protection in a stringent lethal mouse model of infection

Having demonstrated protective efficacy for ADI-63530 against CCHFV-IbAr10200 challenge (**Figure 7A-C**), we tested its cross-clade protective efficacy against Afg09, Turkey2004, and Oman in STAT1^−/−^ (signal transducer and activator of transcription 1 knockout) mice. STAT1^−/−^ mice are more susceptible to a broad range of CCHFV isolates compared to IFNAR1^−/−^ mice^50^, and were therefore used to assess broad-spectrum efficacy. STAT1^−/−^ mice were challenged with either 100 PFU of Afg09 or with 1000 PFU of Turkey2004 or Oman and subsequently treated with 1 mg per mouse of ADI-63530, ADI-58062, c13G8, or vehicle 1- and 4-days post-challenge (2 mg/mouse total). ADI-58062 was included in these studies to investigate the extent to which protection correlates with binding affinity, as it exhibited the highest binding affinities against Afg09, Oman, and Turkey2004 of all lead mAbs (**Figure 3D**). Overall, survival was relatively poor regardless of the mAb used for treatment (**Supplementary Figure S16**), suggesting that these mAbs cannot provide significant protection under more stringent challenge conditions.

Considering the poor survival observed in the previous study, a third challenge study was conducted utilizing less stringent infection conditions to gain a better understanding of the relationship between cross-clade protective efficacy breadth and GP38 antibody bin. Each of our six lead candidates, in addition to c13G8, was tested in this study. Previous results have shown that 13G8 is 80–100% protective against a CCHFV-Turkey2004 challenge in STAT1^−/−^ mice when given 30 minutes post-exposure at a dose of 0.25 mg^28^. To enhance the stringency, mice were treated with a slightly lower dose of 0.2 mg per mouse. STAT1^−/−^ mice were challenged with either 100 PFU of Afg09 or 1000 PFU of Turkey2004 or Oman and subsequently treated with 0.2 mg per mouse of our six lead mAbs 30 minutes post-challenge.

Although none of the c13G8-treated mice survived challenge with CCHFV-Afg09 (**Figure 7D**), 90% and 100% of the c13G8-treated mice survived challenge against CCHFV-Turkey2004 (**Figure 7G**) and CCHFV-Oman (**Figure 7J**), respectively. Only two antibodies, one targeting bin III+IV+V (ADI-46138) and the other targeting bin II (ADI-58048), were partially protective against all tested viruses; CCHFV-Afg09 (27 and 30%, **Figure 7D–E**), CCHFV-Turkey2004 (∼64% and 27%, respectively; **Figure 7G–H**), and CCHFV-Oman (80% and 60%, respectively; **Figure 7J–K**). While other antibodies from bins III–V, including ADI-63530 (bin III+IV) and ADI-63547 (bin IV+V) were not protective against CCHFV-Afg09 (**Figure 7D–E**), they were broadly protective against CCHFV-Turkey2004 (∼83% and 45%, respectively; **Figure 7G–H**) and CCHFV-Oman (60% and 80%, respectively; **Figure 7J–K**), similar to what was observed for c13G8. Apart from ADI-58048 (bin II), other mAbs from bins I–II (ADI-58062 and ADI-58026) demonstrated minimal-to-no cross-clade protection. Relative to CCHFV-Agf09, overall survival across all mAbs was greater against CCHFV-Turkey2004 and CCHFV-Oman, though a prolonged course of disease for CCHFV-Turkey2004 and CCHFV-Oman was observed whereby animals exhibited clinical signs of disease ranging from days 4 through 15 (**Figure 7I**) and 4 through 11 (**Figure 7L**). Taken together, these data show that antibodies targeting epitope bins III–V (ADI-46138, ADI-63530, ADI-63547, and c13G8) exhibit the best protection across isolates, including CCHFV-IbAr10200, Turkey2004, and Oman. However, antibodies targeting bins I–II (namely ADI-58048 and ADI-58026) elicit some cross-protection, albeit less than that of bins III–V antibodies. Furthermore, although there appears to be an inverse correlation across isolates between protective efficacy and binding affinity (i.e., lower affinity antibodies were more protective), this relationship is not statistically significant (**Supplementary Figure S15**). Overall, ADI-46138 (bin III+IV+V) and ADI-58048 (bin II) emerged as lead GP38 mAbs by providing partial protection against all four CCHFV isolates tested (IbAr10200, Afg09, Turkey2004, and Oman).

## DISCUSSION

GP38 is a validated target for the development of mAb-based therapeutics and vaccines^28,29,51^. Moreover, isolation of protective mAbs from human survivors of infection has been shown to be a promising approach for the development of therapeutics against a number of different viruses^30,52–60^. Herein, we isolated and characterized a large panel of GP38-specific mAbs from human survivors of CCHFV infection in Uganda. Several of these mAbs, particularly ADI-46138 and ADI-58048, were found to be as protective as, or more so than, the previously described murine mAb 13G8 against multiple CCHFV isolates in our animal model systems. Further study of these lead candidates could give insight into regions on the GP38 surface that are important for pathogenesis.

Previous reports have determined the presence of five unique antigenic sites on GP38^27,29,61^. Utilizing our sizable antibody panel, we confirmed, structurally mapped, and characterized each of the five distinct antigenic sites and described the existence of 11 novel overlapping antibody competition “bins” that span the GP38 protein (**Figures 2–3** and **5–6**, **Supplementary Table S1**). Although antibodies bind across GP38, we observed two distinct binding regions: one comprising the N-terminal 3-helix bundle and adjacent loops from the β-sheet (bins III–V), and the other comprising the second variable loop and C-terminal β-hairpin of β12-13 (bins I-II) (**Figure 5C**). Interestingly, we observed variation in protection between epitope bins such that the antibodies targeting bins III–V were overall more protective than the antibodies targeting bins I–II (**Figure 7**). Paired with affinity data (**Figure 3**), these results suggest that higher affinity mAbs are not necessarily the most protective. Similarly, although previously described human mAb CC5-17 has a higher affinity to GP38 than does 13G8, it was poorly protective^27^. Moreover, previous studies have demonstrated that non-neutralizing protective antibodies often function through Fc-mediated mechanisms^62–66^; in fact, reports have characterized a partial contribution of Fc-mediated functions in the protection provided by 13G8^28,29^. One possibility is that mAbs from varied epitope bins differentially engage Fc receptors and complement factors, an observation seen in the studies of filoviruses and influenza viruses^67–69^. Taken together, these data suggest that binding affinity, and even epitope bin, do not exclusively determine protective efficacy provided by GP38 mAbs. However, in this work, only a single antibody from each bin was selected for further *in vitro* and *in vivo* characterization, limiting our ability to draw definitive conclusions regarding the relationship between epitope bin and protection, warranting additional follow-up studies utilizing multiple antibodies from each epitope bin.

CCHFV is the most genetically divergent of the arboviruses^2,6,10^. GP38, in particular, exhibits high diversity among lineages. Sequence diversity of GP38 has been cited as the reason for the poor cross-clade efficacy of 13G8^29^. Along with variable protection between antigenic sites, we observed variable protection within overlapping epitope bins across the divergent isolates (**Figure 7**). Our knowledge regarding GP38 function and its contribution(s) to pathogenesis is limited. Therefore, a plausible explanation for the observed differences in mAb efficacy across isolates *in vivo* is rooted in the unidentified pathogenic functions of GP38 and the ability of these mAbs to limit these functions. Epitope-bin-specific protection could be explained by a potential structural role for GP38. GP38 has been speculated to form a complex with Gn on the virion surface, acting as the head region of the attachment protein, as suggested by an AlphaFold2-predicted model^70^. In this model, the epitopes of bins I and II are near the GP38-Gn interface while those of bins III–V are predicted to be orientated away from Gn, potentially making the bin III–V epitopes more accessible for antibodies to bind and mediate protection. A more thorough investigation into the association of GP38 and Gn is needed to resolve the structural relevance of GP38 on the viral surface and further scrutinize the implications on epitope accessibility for GP38-specific mAbs. Further uncovering the pathogenic functions of GP38 will strengthen our understanding of the mechanisms of protection utilized by our panel of GP38-specific mAbs.

Cocktails of mAbs have shown promise for the broad-spectrum treatment of diverse viral isolates^52,53,71–73^. Earlier work in the context of Ebola virus infection demonstrated that “enabling pairs” of neutralizing and non-neutralizing mAbs can result in potent neutralization and complete protection, even though neither antibody alone was able to provide complete protection^74^. Neutralizing Gc-specific antibodies have been isolated from human survivors of infection and developed into a bi-specific mAb, DVD-121-801, resulting in robust post-exposure protection against a lethal CCHFV-IbAr10200 challenge^30^. Although DVD-121-801 exhibits potent neutralization across multiple clades of CCHFV, weaker neutralization was observed for Clade V isolates Hoti and Turkey2004, suggesting that *in vivo* potency against Clade V isolates may be impacted, although it has not been experimentally tested. In the context of this study, future work should consider combining potent Gc-specific mAbs, such as DVD-121-801, with GP38-specific mAbs (e.g., ADI-46138) to improve potency and maximize cross-clade protective efficacy. Combining multiple GP38 mAbs targeting different epitope regions could also be a useful approach for broadening efficacy and increasing potency. The wealth of structural data and characterization pertaining to antigenic sites across the GP38 protein described in this study should facilitate efforts to identify optimal mAb pairings as well as inform vaccine development.

## ACKNOWLEDGMENTS

All IgGs were sequenced by Adimab’s Molecular Core, and yeast-expressed mAbs and Fabs were produced by Adimab’s High Throughput Expression group. Biolayer interferometry binding experiments with IgGs were performed by Adimab’s Protein Analytics group. We thank Christina Spiropoulu, Eric Bergeron, and Marko Zivcec at the Centers for Disease Control and Prevention for kindly providing the plasmids and protocols necessary to generate tecVLPs. The following reagents were obtained from the Joel M. Dalrymple - Clearance J. Peters USAMRIID Antibody Collection through BEI Resources, NIAID, NIH: Monoclonal Anti-Crimean-Congo Hemorrhagic Fever Virus Pre-Gn Glycoprotein, Clone 13G8 (produced *in vitro)*, NR-40294; Monoclonal Anti-Crimean-Congo Hemorrhagic Fever Virus Pre-Gn Glycoprotein, Clone 7F5 (produced *in vitro*), NR-40281; Monoclonal Anti-Crimean-Congo Hemorrhagic Fever Virus Pre-Gn Glycoprotein, Clone 8F10 (produced *in vitro*), NR-40282; and Monoclonal Anti-Crimean-Congo Hemorrhagic Fever Virus Nucleocapsid Protein, Clone 9D5 (produced *in vitro*), NR-40270. We thank members of all of our groups and the Prometheus consortium for their feedback on preliminary versions of the manuscript. We would like to thank Drs. Axel Brilot and Evan Schwartz at the Sauer Structural Biology Laboratory at UT Austin for assistance with cryo-EM data collection. We would also like to thank Kandis Cogliano for her project management support. Research was supported by NIAID of the National Institutes of Health (NIH) under award number U19AI142777 (Centers of Excellence in Translational Research) to K.C., L.M.W., J.M.D., J.S.M., L.Z., and A.S.H., award number R01AI152246 to K.C. and J.S.M, and The Welch Foundation under award number F-0003-1962064 awarded to J.S.M. We acknowledge the University of Texas College of Natural Sciences and award RR160023 of the Cancer Prevention and Research Institute of Texas for support of the EM facility at the University of Texas at Austin. Results shown in this report are derived from work performed at Argonne National Laboratory, Structural Biology Center (SBC) at the Advanced Photon Source. SBC-CAT is operated by UChicago Argonne, LLC, for the U.S. Department of Energy, Office of Biological and Environmental Research under contract DE-AC02-06CH11357. The content is solely the responsibility of the authors and does not necessarily represent the official views of our institutions or funders. Opinions, interpretations, conclusions, and recommendations are those of the author and are not necessarily endorsed by the U.S. Army.

## AUTHOR CONTRIBUTIONS

Conceptualization, O.S.S., C.K.H., S.R.M., K.C., L.M.W., J.M.D., L.Z., N.T.P., J.S.M., A.S.H.

Methodology, O.S.S., C.K.H., S.R.M.

Formal Analysis, O.S.S., C.K.H., S.R.M.

Investigation, O.S.S., C.K.H., S.R.M., D.A., A.I.K., A.W., R.R.B., A.M., M.M., E.C., L.S., J.L., J.B., JB., V.H., M.D.

Resources, L.L., S.B., J.J.L., A.S.H., J.M.D., S.R.M.

Writing – Original Draft, O.S.S., C.K.H., S.R.M., M.D.

Writing – Reviewing & Editing, all authors

Visualization, O.S.S., C.K.H., S.R.M., N.T.P., J.S.M., A.S.H.

Supervision, N.T.P., L.M.W., J.M.D., A.S.H., J.S.M., L.Z., K.C.

Funding Acquisition, K.C., L.M.W., J.M.D., J.S.M., A.S.H, L.Z.

## METHODS

### Patient recruitment and ethics statement

CCHFV convalescent donors were recruited as described previously^28,30^. Briefly, donors with documented clinical history of CCHF infection between 2013 and 2017 in Agago and Nakaseke districts, Uganda were recruited through the Uganda virus Research Institute, Entebbe, Uganda. The study was approved by the Helsinki committees of Uganda Virus Research Institute (UVRI), Entebbe, Uganda (reference number GC/127/13/01/15); Soroka Hospital, Beer Sheva, Israel (protocol number 0263-13-SOR); and the Ugandan National Council for Science and Technology (UNCST) (registration number HS1332). Written informed consent was obtained and a personal health questionnaire was completed for each donor who participated in this study. Study participants were adults, or minors with parental consent, and were not related. All experiments were performed in accordance with the relevant guidelines and regulations.

### Cell lines

VeroE6 and Vero cells, immortalized epithelial cell lines isolated from the kidney of an adult female African grivet monkey (RRID:CVCL-0574 and CVCL-0059, respectively), were obtained from the American Type Culture Collection (ATCC). SW-13 cells, a cell line isolated from the adrenal gland and cortex of a 55-year-old female patient with carcinoma (RRIDD:CCL-105), were obtained from ATCC. BSR-T7 cells (RRID:CVCL_RW96), generated by stable T7 RNA polymerase expression in BHK-21 cells, were a kind gift from K.-K. Conzelmann. The parent cell line (RRID: CVCL_1915) was isolated from the kidney of a 1-day-old male golden hamster. All cell lines were cultured in Dulbecco’s Modified Eagle Medium (DMEM; ThermoFisher Scientific) enriched with 10% fetal bovine serum (Bio-Techne), 1% GlutaMAX (ThermoFisher Scientific), and 1% penicillin-streptomycin (ThermoFisher Scientific). All cell lines were maintained in a 37 °C incubator supplied with 5% CO_2_. Cell lines were not authenticated following purchase.

### Viruses

The authentic CCHFV isolates CCHFV-IbAr10200, CCHFV-Afg09-2990 (labeled as ‘Afg09’), CCHFV-Turkey2004, and Oman-199809166 (labeled as ‘Oman’) were used in this study.

### Animal models

3–8-week-old male and female B6.129S(Cg)-*Stat1^tm1Dlv^*/J mice (STAT1^−/−^; strain #012606; The Jackson Laboratory)^50,75^ and 5–8-week-old male and female B6(Cg)-*Ifnar1^tm1.2Ees^*/J mice (IFNAR^−/−^; strain #028288; Charles River)^48,49^ were used in animal challenge experiments. Animals were provided with food and water *ad libitum* and housed in individually ventilated cages.

Murine challenge studies were conducted under Institutional Animal Care and Use Committee (IACUC)-approved protocols in compliance with the Animal Welfare Act, PHS Policy, and other applicable federal statutes and regulations. The facilities where these studies were conducted (USAMRIID) are accredited by the Association for Assessment and Accreditation of Laboratory Animal Care, International (AAALAC) and adhere to the principles stated in the Guide for the Care and Use of Laboratory Animals, National Research Council, 2013. IACUC-approved euthanasia criteria were defined as follows: mouse displays severely hunched posture, inability or reluctance to move, appears weak (staggering when moving around cage), or has labored breathing.

### rGP38 serum ELISA

High-binding half-area plates (Greiner Bio-One) were coated with 50 µL of IbAr10200 rGP38 at 5 µg/mL. Plates were incubated overnight at 4 °C. Plates were then blocked with 100 µL of 5% BSA/PBS and flicked to remove liquid. Serum was serially diluted 5-fold in PBS. 50 µL of each dilution was added to plates and incubated for 1 hour at room temperature. Plates were washed 3X with PBS plus 0.05% Tween 20 (PBST). Anti-Human-HRP (Invitrogen) was diluted 1:5000 in 1% BSA/PBS. 50 µL of the diluted solution was added to plates and incubated for 1 hour at room temperature. Plates were again washed 3X with PBST. 50 µL of KPL Blue Sure Substrate (Seracare) was added to plates. Plates were incubated for 5 minutes at room temperature, and the reaction was stopped with 50 µL of 2 N H_2_SO_4._ OD_450_ was measured with a Perkin Elmer EnVision multimode plate reader. Data were plotted and analyzed using GraphPad Prism Software V9.5.1; a Sigmoidal, 4PL curve was fit to interpolate data.

### Single B cell sorting

B cells were eluted from PBMCs using a MACS Human B Cell isolation kit (Miltenyi Biotec). B cells were stained with rGP38 (IbAr10200) that had been tetramerized at 25 nM using Streptactin-PE (IBA Lifesciences) and Streptactin-APC (IBA Lifesciences). B cells were simultaneously stained with rGP38-Streptactin-PE and rGP38-Streptactin-APC tetramers for 1 hour on ice. Cells were washed twice in buffer (PBS, FBS, EDTA). Next, B cells were stained with a panel of antibodies. Donor 1 PBMCs were stained with a cocktail of anti-human CD3 PerCP-Cy5.5 (Biolegend), CD8 PerCP-Cy5.5 (Biolegend), CD14 PerCP-Cy5.5 (Invitrogen), CD16 PerCP-Cy5.5 (Biolegend), propidium iodide (PI) (Invitrogen), CD19 PE-Cy7 (Biolegend), CD27 BV510 (BD Biosciences), IgM BV711 (BD Biosciences), IgD BV421 (Biolegend), IgG BV605 (BD Biosciences), and IgA AF488 (Abcam). Donor 5 and 6 PBMCs were stained with a cocktail of anti-human CD3 PerCP-Cy5.5 (Biolegend), CD8 PerCP-Cy5.5 (Biolegend), CD14 PerCP-Cy5.5 (Invitrogen), CD16 PerCP-Cy5.5 (Biolegend), PI (Invitrogen), CD19 PE-Cy7 (Biolegend), CD20 PE-Cy7 (Biolegend), CD27 BV510 (BD Biosciences), IgM AF488 (Biolegend), and IgD BV421 (Biolegend). B cells were washed twice in buffer and run on a FACS Aria Fusion Cytometer (BD Biosciences). B cells were sorted into Super Script III reaction buffer (ThermoFisher Scientific) in 96-well Costar plates and frozen at −80 °C.

### Amplification of antibody variable genes

cDNA was synthesized using SuperScript III Reverse Transcriptase (ThermoFisher Scientific). Antibody VH and VL genes were amplified following previously designed methods^76^. Gene amplification with HotStartTaq Plus Polymerase (Qiagen) was carried out in two steps. IgG-, IgA-, IgM-specific primers were used in the first reaction. Primers with 5′ and 3′ homology domains, specific to plasmids used for cloning into an engineered strain of *S. cerevisiae,* were used in the second reaction.

Cloning into engineered *S. cerevisiae*

Amplified variable genes were transformed into *S. cerevisiae* through the lithium acetate method^77^. One colony of engineered *S. cerevisiae* was inoculated in yeast extract-peptone-dextrose medium for 14–16 hours. Yeast were washed twice in dH_2_O and resuspended in dH_2_O (67 µL). Resuspended yeast were mixed with variable gene product (10 µL of unpurified VH and 10 µL of unpurified Vκ or Vλ product), digested plasmid (200 ng), 50% w/v polyethylene glycol 3350 (240 µL), 1 M lithium acetate, and boiled salmon sperm DNA (10 µL). Contents of the transformation were incubated at 42 °C. After a 45-minute incubation, yeast were washed twice with dH_2_O, resuspended in selective growth medium, and grown for 48 hours at 30 °C.

### Expression and purification of IgG and Fab

#### Production in yeast

Full length IgG_1_ and Fabs were produced and purified as previously described^78^. Briefly, cultures were grown in 24-well plates for 6 days at 30 °C and 80% relative humidity with shaking at 650 rpm on a Multitron Shaking Incubator (Infors HT). Cultures were centrifuged to obtain supernatants, which were purified by Protein A chromatography. Bound IgGs were eluted with 200 mM acetic acid (pH 3.5), 50 mM NaCl and neutralized with 1/8 v/v 2 M HEPES (pH 8.0). IgGs were buffer exchanged into PBS (pH 7.0) and stored for later use.

To produce Fabs, IgGs were papain-digested for 2 hours at 30 °C. The reaction was quenched with iodoacetamide. The material was passed over a Protein A column to remove undigested IgGs and Fc domains. The flow-through was collected and Fabs were purified using CaptureSelect^TM^ IgG-CH1 affinity resin (ThermoFisher Scientific). 200 mM acetic acid (pH 3.5), 50 mM NaCl was used to elute Fabs, which were neutralized with 1/8 v/v 2 M HEPES (pH 8.0). Fabs were buffer exchanged into PBS (pH 7.0) and stored for later use.

#### Production in mammalian cells

For IgGs used for *in vitro* and *in vivo* studies, and later used to produce Fabs for structural studies (ADI-58048, ADI-46143, ADI-46138, ADI-46158, and 13G8), genes encoding the variable regions were ordered as gBlocks (Integrated DNA Technologies) with a 15-base-pair 5′ overlap to a murine IgKVIII secretion signal and a 15-base-pair 3′ overlap to the appropriate constant region (human kappa, human lambda or human IgG1). The variable regions were cloned into pCDNA 3.4 (ThermoFisher Scientific) vectors previously constructed with a mouse IgKVIII signal sequence and each constant region. In-Fusion enzyme (Takara Bio) was used to insert the gBlocks between the secretion signal and the constant region.

Antibodies were transiently expressed in ExpiCHO cells (ThermoFisher Scientific) following the high-titer protocol for CHO Expifectamine (ThermoFisher Scientific). Cultures were centrifuged 9–10 days after transfection, and the supernatants were filtered and loaded onto a HiTrap MabSelect SuRe affinity column (Cytiva) using an AKTA Pure FPLC system. The column was washed with 10 column volumes of PBS pH 7.2 and antibodies were eluted with Pierce IgG elution buffer (ThermoFisher Scientific). Fractions containing the antibody were combined and neutralized to ∼pH 7 with 1 M Tris pH 7.8.

To produce Fabs of ADI-58048, ADI-46143, ADI-46158, and c13G8 used in structural studies, purified IgG was digested with LysC at a 1:2000 molar ratio of LysC:IgG overnight at 37 °C. A cOmplete™ Protease Inhibitor Cocktail tablet (Sigma-Aldrich) was dissolved into the reaction before loading the digested IgG mixture over a CaptureSelect^TM^ IgG-CH1 affinity resin (ThermoFisher Scientific) to bind the Fabs. The column was washed with 1X PBS followed by elution of the Fabs with 100 mM glycine pH 3.0 into a neutralization buffer of 100 mM Tris pH 8.0.

For IgGs used to produce Fabs for structural studies (ADI-46152, ADI-58026, ADI-58062, ADI-63530, and ADI-63547), the heavy and light chain variable regions were cloned into Igγ1 and either human Igκ or Igλ vectors, respectively. To later generate Fabs from the IgG, a human rhinovirus (HRV) 3C protease site was present at the hinge region of the heavy chain in the Igγ1 vector. Plasmids encoding both the heavy chain and light chain for each antibody were co-transfected into FreeStyle 293-F cells (Invitrogen) using polyethylenimine. Secreted IgG was purified from the culture supernatants via Pierce™ Protein A Plus Agarose resin (ThermoFisher Scientific). The IgG eluent was further purified via SEC with a HiLoad 16/600 Superdex 200 column (GE Healthcare Biosciences) in 2 mM Tris pH 8.0, 200 mM NaCl, and 0.02% NaN_3_.

To produce Fabs for structural studies, purified IgG was bound to Pierce™ Protein A Plus Agarose resin (ThermoFisher Scientific) and washed with 1X PBS. The IgG-bound Protein A resin was removed from the column holder and added to a conical tube with 1X PBS buffer and 10% w/w HRV 3C protease and nutated on a rotating shaker for 2 hours at 23 °C. Following the cleavage reaction, the Fc domains remained bound to the Protein A resin and the Fabs were collected in the nutated flow-through. Purified Fabs were stored for later use.

### Biolayer interferometry binding analysis of antibodies to rGP38

For all experiments, a Fortébio Octet HTX (Sartorius) was used. All steps of the experiments were performed at 25 °C with an orbital shaking speed of 1,000 rpm and all reagents were formulated in PBSF (PBS with 0.1% w/v BSA). For avid binding experiments, biotinylated rGP38 at 100 nM was loaded onto streptavidin biosensors for 10–40 seconds, providing load levels of 0.30–0.40 nm. The sensors were then soaked for 30 minutes in PBSF, dipped in 100 nM IgG for 180 seconds, and dipped into PBSF for 180 seconds to measure dissociation. For monovalent binding, IgGs were loaded onto AHC biosensors (0.6–1.2 nm) at 100 nM for 30 minutes. Considering that antigens contained a twin-strep-tag, the sensors were blocked with 100 µM biocytin for 10 minutes to saturate any remaining streptavidin binding sites. Sensors were incubated for 60 seconds in PBSF to establish a baseline. Next, sensors were dipped in 100 nM antigen for 180 seconds followed by PBSF for 180 seconds to measure dissociation. Data for which binding responses were greater than 0.05 nm were aligned, interstep-corrected to the association step, and subsequently fit to a 1:1 binding model using Fortébio Octet Data Analysis, v 11.1.

### Surface plasmon resonance binding analysis of antibodies to rGP38

For all experiments, the Carterra LSA (Carterra USA) was used. Kinetic analysis was conducted in HBS-ET running buffer (10 mM HEPES pH 7.4, 150 mM NaCl, 3 mM EDTA, 0.01% Tween-20) (Carterra USA) at 25 °C. The standard amine coupling step was conducted in 25 mM MES buffer (Carterra USA) with 0.05% Tween-20. The sample compartment was maintained at a temperature of 20 °C for the duration of the experiment.

Standard amine coupling (1:1 EDC:NHS) was used to covalently couple a goat anti-human Fc antibody (Jackson ImmunoResearch) to the HC30M chip; the chip was then blocked with 1.0 M ethanolamine pH 8.5. Next, antibodies (100 nM in running buffer) were flowed for 5 minutes over discrete regions of interest on the chip surface. Once the antibody samples were captured to the sensor surface, kinetic measurements were collected in cycles. For a given antigen, the loaded biosensor array was first exposed to running buffer (60 s), then three blank buffer injections (300 s association and 300 s dissociation). This was followed by a series of four antigen injections (300 s association and 3000 s dissociation) of increasing concentration (1.56 – 100 nM). At the end of each cycle, all surfaces were regenerated via two 30 seconds injections of 10 mM glycine, pH 1.7.

All kinetic data were reference subtracted using interspot reference surfaces evenly distributed throughout the biosensor surface array. The data were then y-axis aligned, x-axis aligned, corrected for baseline drift using a minimum baseline drift parameter of 4 RU, and blank subtracted from the leading (third of three) blank injection. Sensorgrams were filtered using a minimum spike height of 5 RU and width of 3 points before being cropped, beginning just after the start of the association and ending just before the end of the dissociation. The processed sensorgrams were then fit to a 1:1 binding model with floating T_0_ using the Carterra LSA Kinetics Software version 1.7.1.3055 (Carterra, USA).

### Antibody competition assays

#### Biolayer interferometry

For all experiments, a Fortébio Octet HTX (Sartorius) was used. All steps of the experiments were performed at 25 °C with an orbital shaking speed of 1,000 rpm and all reagents were formulated in PBSF (PBS with 0.1% w/v BSA). IgGs were loaded onto AHC biosensors (0.7–1.5 nm) at 100 nM for 30–600 seconds, providing load levels of 1.0–1.3 nm. The sensors were blocked for 10 minutes with an inert human antibody at 0.5 mg/mL to fill unoccupied binding sites and then were equilibrated for 30 minutes in PBSF. To check for cross-interactions on the protein surface, prior to binding analysis, the sensors were dipped in 300 nM control antibody for 90 seconds. After a baseline step in PBSF for 60 seconds, the sensors were exposed first to antigen (100 nM) for 180 seconds, then to control antibody (300 nM) for an additional 180 seconds. Data were then y-axis normalized and interstep-corrected using Fortébio Octet Data Analysis, v 11.1. Binding of the secondary antibody indicates a non-competitor (unoccupied epitope), whereas no binding indicates a competitor antibody (epitope blocking).

#### Yeast presentation

Biotinylated CCHFV GP38 (IbAr10200; 50 nM) was incubated with a 20-fold excess of anti-CCHFV-GP38 Fab (1 µM) for 30 minutes at room temperature. Pre-complexed biotinylated CCHFV GP38 and Fab mixtures were incubated with yeast expressing full-length anti-CCHFV-GP38 IgG for 5 minutes at room temperature. Yeast were washed two times with PBSF (PBS with 0.1% w/v BSA) to remove any unbound GP38-Fab complexes. Samples were incubated for 30 minutes on ice with a cocktail of streptavidin Alexa Fluor 633 (Invitrogen; to detect bound GP38), goat F(ab′)2 anti-human kappa FITC and goat F(ab′)2 anti-human lambda FITC (SouthernBiotech; to detect antibody expression), and PI (Invitrogen; to detect cell viability). After staining, samples were run on a FACSCanto II flow cytometer (BD Biosciences). Competition levels were assessed by calculating the fold reduction between a known non-competitive isotype control IgG and an IgG of interest; bound GP38 levels were normalized to light chain expression. The following equation was used to calculate the fold reduction with mean fluorescence intensity (MFI); Fold Reduction = (AF633 MFI/FITC MFI)_No-competition_/(AF633 MFI/FITC MFI)_Competition_. Antibodies with a calculated fold reduction greater than 10 were considered competitive with the pre-complexed Fab.

### CCHFV GP38 yeast display and epitope mapping

#### Display of CCHFV GP38 on the surface of yeast

The sequence encoding GP38 from the CCHFV-IbAr10200 *GP* gene (GenBank Accession: NC_005300.2) was inserted into a plasmid containing an N-terminal HA tag-G_4_S linker and a G_4_S-HA tag C-terminal linker. The plasmid was transformed and expressed as previously described^30^.

#### CCHFV GP38 library construction

PCR was carried out with an error-prone polymerase (Agilent, GeneMorph II Random Mutagenesis Kit) to create a randomly mutagenized GP38 library as previously described^79^.

#### Titration of anti-GP38 mAbs on yeast displayed GP38

Antibodies used in epitope mapping studies were titrated against yeast displaying GP38 to adequately calculate EC_50_s and EC_80_s for each antibody. Yeast were induced to express non-mutagenized GP38 as noted above. Antibodies were titrated from 100 nM in two-fold, 12-point serial dilutions. Once an OD_600_ of 0.1 was achieved, the non-mutagenized GP38-expressing yeast were mixed with each antibody dilution and incubated on ice for one hour. Yeast cells were washed two times with PBSF and subsequently stained for 30 minutes on ice with a cocktail of anti-HA APC antibody (Biolegend, Clone: 16B12, dilution 1:100), goat F(ab′)_2_ anti-human IgG PE (SouthernBiotech, dilution 1:100), and PI (Invitrogen, 1:100 dilution). After staining, samples were run on a FACSCanto II flow cytometer (BD Biosciences). PE MFIs were plotted against antibody concentrations; EC_50_ and EC_80_ concentrations were calculated using GraphPad Prism 9.

#### Flow cytometric sorting of mutant GP38 libraries

The mutant GP38 library and non-mutagenized GP38-expressing yeast were induced as noted above. Both the mutant GP38 library and non-mutagenized GP38-expressing yeast were incubated with a solution of each mAb at its respective EC_80_ for one hour on ice. Cells were washed two times in PBSF and further stained with anti-HA APC, anti-human IgG PE, and PI (as described above). Cells were washed and run on a FACSAria (BD Biosciences). Mutagenized GP38 clones that showed reduced binding to each antibody of interest were sorted and cultured in synthetic complete (SC) media minus tryptophan (4% dextrose, 0.1 M sodium phosphate, pH 6.3) for further rounds of selection. The same selection strategy was applied to cultured cells from the first round of selection to carry out a second round of selection. A third and final round of selection occurred; the final selection was a positive selection used to remove any mutagenized clones that were global knock-outs. Cultured cells from the second round of selection were stained with a panel of anti-GP38 antibodies of non-overlapping epitopes to the antibody used in the first round of selection. Cells that bound the non-competing anti-GP38 antibodies were sorted and plated on complete minimal media glucose agar plates minus tryptophan (Teknova). For each library, 100 clones were picked and sequenced.

#### Flow cytometric analysis of single GP38 mutants

Unique clones that came out of selections were induced as described above. GP38 wild-type control clones were induced alongside the clones from selections. Clones were stained with each antibody of interest as well as with an isotype control antibody. Next, clones were stained with each antibody at its respective EC_50_ for one hour on ice. Yeast were washed twice with PBSF. Cells were washed two times in PBSF and further stained with anti-HA (hemagglutinin) APC, anti-human IgG PE, and PI (as described above). Samples were run on a FACSCanto II flow cytometer (BD Biosciences). Percent loss of binding was calculated utilizing the following equation; % of WT Binding = [((IgG MFI/HA MFI)_MUT_ – (IgG MFI/HA MFI)_BACK_)/ ((IgG MFI/HA MFI)_WT_ – (IgG MFI/HA MFI)_BACK_)] × 100. Clones with less than 25% of wild-type binding for a specific antibody were considered to have a mutation critical for binding.

### Cloning, expression, and purification of CCHFV GP38

Recombinant CCHFV GP38 proteins were produced from the following isolates: Oman-199809166 (UniProt: A0A0U3C6Q7), Kosova-Hoti (UniProt: B2BSL7), 200406546-Turkey (UniProt: A0A0U2SQZ0), Afg09-2990 (UniProt: E5FEZ4), and 79121M18 (UniProt: D4NYK3). Gene fragments (Integrated DNA Technologies) of each isolate’s MLD-GP38 sequence encoding for residues 1–515, as denoted by CCHFV IbAr10200 strain GPC numbering, were codon-optimized for human cell expression (GenScript Codon Optimization Tool). Gene fragments were each cloned into a pαH eukaryotic expression vector with a C-terminal HRV 3C protease cleavage site, an 8x HisTag, and a Twin-Strep-tag. The plasmid for CCHFV strain IbAr10200 GP38 was previously reported^28^. To ensure cleavage of the MLD from GP38, a pCDNA3.1 plasmid encoding for furin was co-transfected with each clinical GP38 plasmid at a mass ratio of 1:9 furin:GP38. The two plasmids were transiently transfected into FreeStyle 293 cells (Invitrogen) using polyethylenimine followed by treatment with 5 μM kifunensine to ensure uniform high-mannose glycosylation. Soluble GP38 was secreted from the harvested medium and purified via Ni-NTA resin (Thermo Scientific HisPur™ Ni-NTA Resin). GP38 proteins were further purified via SEC using a Superdex 200 Increase 10/300 GL (GE Healthcare Biosciences) in 2 mM Tris pH 8.0, 200 mM NaCl, and 0.02% NaN_3_.

### Crystallization and data collection

#### GP38 + ADI-46143 Fab

GP38 (from CCHFV-IbAr10200) was incubated at room temperature for 20 minutes with a 1.2-fold molar excess of ADI-46143 Fab and the complex was purified by SEC on a Superdex 200 Increase 10/300 GL (GE Healthcare Biosciences) in 2 mM Tris pH 8.0, 50 mM NaCl, and 0.02% NaN_3_. The GP38-ADI-46143 Fab complex (4.1 mg/mL) underwent crystallization trials via the sitting-drop vapor diffusion method. The crystal from which the diffraction data were obtained was grown in 9.3% w/v PEG 3350, 12.2% v/v isopropanol, 0.2 M ammonium citrate pH 7.5 at a protein:buffer ratio of 1:1. The crystal was looped with 20% ethylene glycol as a cryoprotectant, and flash frozen in liquid nitrogen. The 19-ID beamline (Advanced Photon Source; Argonne National Laboratories) was used to collect the X-ray diffraction data to 2.6 Å resolution.

#### GP38 + c13G8 Fab

GP38 (from CCHFV-IbAr10200) was incubated at room temperature for 20 minutes with a slight molar excess of c13G8 Fab and the complex was purified by SEC on a HiLoad 16/600 Superdex 200 column (GE Healthcare Biosciences) in 2 mM Tris pH 8.0, 200 mM NaCl, and 0.02% NaN_3_. The GP38-c13G8 Fab complex (9.8 mg/mL) underwent crystallization trials through the sitting-drop vapor diffusion method. The crystal used to obtain the diffraction data was grown in 2 M ammonium sulfate, 0.1 M Bis-Tris pH 5.5, 0.01 M cobalt chloride hexahydrate at a protein:buffer ratio of 2:1. The crystal was looped with 20% ethylene glycol as a cryoprotectant and flash frozen in liquid nitrogen. The 19-ID beamline (Advanced Photon Source; Argonne National Laboratories) was used to collect the X-ray diffraction data to 1.8 Å resolution.

### Crystal structure determination, model building, and refinement

Diffraction data from the 19-ID beamline were processed using the CCP4 software^80^, indexed and integrated in iMOSFLM^81^, and scaled and merged in Aimless^82^. Both crystal structures were phased using PhaserMR^83^ and refined and built using COOT^84^ and Phenix^85^. The GP38+13G8 crystal structure was refined to a final R_work_/R_free_ of 20.0%/21.5% (**Supplementary Table S6**). The GP38+ADI-46143 crystal structure was refined to a final R_work_/R_free_ of 17.7%/21.7% (**Supplementary Table S6**). The crystal structures were displayed in PyMOL^86^.

### Cryo-EM sample preparation and data collection

#### GP38+ADI-58026+ADI-63547 Fabs and GP38+ADI-58062+ADI-63530 Fabs

For the GP38+ADI-58026 Fab+ADI-63547 Fab complex, a 0.4 mg/mL complex was prepared by combining purified IbAr10200 GP38^28^ with a 1.8-fold molar excess of each Fab followed by incubation for 30 minutes at room temperature in 2 mM Tris pH 8.0, 200 mM NaCl, 0.02% NaN_3_, and 0.03% amphipol A8-35. For the GP38+ADI-58062 Fab+ADI-63530 Fab complex, a 0.4 mg/mL complex was prepared by combining purified IbAr10200 GP38^28^ with a 1.8-fold molar excess of each Fab followed by incubation for 30 minutes at room temperature in 2 mM Tris pH 8.0, 200 mM NaCl, 0.02% NaN_3_, and 0.03% amphipol A8-35.

#### GP38+ADI-46152+ADI-58048 Fabs and GP38+ADI-46143+ADI-46158 Fabs

Two complexes were prepared by using purified IbAr10200 GP38^28^ complexed with a 1.2– 1.5-fold molar excess of ADI-46152 Fab and ADI-58048 Fab or ADI-46143 Fab and ADI-46158 Fab. Complexes were incubated for 20 minutes at 23 ℃ before further purification via SEC on a Superdex 200 Increase 10/300 GL (GE Healthcare Biosciences) in 2 mM Tris pH 8.0, 200 mM NaCl, and 0.02% NaN_3_. The GP38+ADI-46152+ADI-58048 complex was used at a concentration of 0.5 mg/mL and the GP38+ADI-46143+ADI-46158 was at a concentration of 0.4 mg/mL.

#### Cryo-EM Data Collection

A 3 µL aliquot of each complex was applied to a Quantifoil 1.2/1.3 Cu300 grid that was glow discharged for 25 seconds at 15 mAmps (PELCO easiGlow™ Glow Discharge Cleaning System). A Vitrobot Mark IV (ThermoFisher Scientific) was used to plunge freeze the grids at 10 ℃ and 100% humidity with a blot time of 3.5 seconds, blot force of −4, blot total of 1, and wait time of 2 seconds. 2,504 micrographs for the GP38+ADI-58026 Fab+ADI-63547 Fab complex, 3,647 micrographs for the GP38+ADI-46152 Fab+ADI-58048 Fab complex, 2,962 micrographs for the GP38+ADI-58062 Fab+ADI-63530 Fab complex, and 1,485 micrographs for the GP38+ADI-46143 Fab+ADI-46158 Fab complex, were collected using a FEI Titan Krios equipped with a K3 detector (Gatan). Data were collected with a 30° tilt at a magnification of 105,000x, corresponding to a calibrated pixel size of 0.81 Å/pixel and a total electron dose of 80 e^−^/Å^2^. Statistics for each data collection are in **Supplementary Table S5**.

### Cryo-EM data processing, model building, and refinement

On-the-fly data processing was performed in cryoSPARC Live^87^, and included motion correction, defocus estimation, micrograph curation, particle picking, particle extraction, and particle curation through iterative streaming 2D classification. Data processing and refinement of all datasets were performed using cryoSPARC v3.2 and subsequent versions. Statistics for each dataset are in **Supplementary Table S5**.

For the GP38+ADI-58026 Fab+ADI-63547 Fab complex, several rounds of 2D classification and *ab initio* reconstruction were performed to refine the particle stack for the complex with two Fabs bound to GP38, as the lower binding affinity for ADI-63547 led to heterogeneity in the Fab occupancy. After volumes were refined for the complex bound with two Fabs, the volume underwent homogeneous and non-uniform refinement before another round of non-uniform refinement using particles from the extracted particle stack. The dataset underwent two rounds of heterogeneous, homogeneous, and non-uniform refinements. Duplicate particles were then removed followed by a non-uniform refinement. The final map was sharpened using DeepEMhancer^88^. The EM processing pipeline is summarized in **Supplemental Figure S10**.

For the GP38+ADI-46152 Fab+ADI-58048 Fab complex, selected particles underwent *ab initio* 3D reconstruction followed by heterogeneous refinement. For the best class, homogeneous and non-uniform refinements were performed, then curated particles were further refined using another round of heterogeneous refinement. The best class underwent homogeneous and non-uniform refinement, followed by extracting the curated particles without Fourier cropping and removing duplicate particles with non-uniform refinements between each step. The final volume was sharpened using DeepEMhancer^88^. The model was built iteratively using PHENIX^85^, COOT^84^, and ISOLDE^89^. The EM processing pipeline is summarized in **Supplemental Figure S11**.

For the GP38+ADI-58062 Fab+ADI-63530 Fab complex, extracted particles underwent two rounds of 2D classification to generate a curated particle stack. Particles were further processed using *ab initio* 3D reconstruction and heterogeneous refinement. From the best class, a non-uniform refinement was conducted before extracting the particles without Fourier crop followed by another round of non-uniform and heterogeneous refinements. Next, the best class underwent homogeneous refinement and non-uniform refinement before duplicate particles were removed. Lastly, a non-uniform refinement was performed on the resulting map before the map was sharpened using DeepEMhancer^88^. The EM processing pipeline is summarized in **Supplemental Figure S12**.

For the GP38+ADI-46143 Fab+ADI-46158 Fab complex, extracted particles were curated via 2D classification followed by iterative rounds of *ab initio* reconstruction, heterogeneous refinement, homogeneous refinement, and non-uniform refinement. In some steps, volumes obtained from the processing of a smaller initial particle stack were used. After a final non-uniform refinement, the maps were processed with DeepEMhancer^88^. The EM processing pipeline is summarized in **Supplemental Figure S13**.

### Polyreactivity assay

A polyreactivity assay was carried out as previously described^90^. Briefly, soluble cytosolic protein (SCP) and soluble membrane protein (SMP) preps were extracted from Chinese hamster ovary (CHO) cells and were biotinylated using an NHS-LC-Biotin kit (ThermoFisher Scientific). Yeast displaying IgGs on their surface were incubated with biotinylated SCP and SMP preps at a 1:10 dilution in PBSF (PBS with 0.1% w/v BSA) and incubated on ice for 20 min. Yeast cells were then washed two times in PBSF and further stained with a cocktail of ExtraAvidin-R-PE (Sigma Aldrich, dilution 1:50), anti-human kappa FITC (Southern Biotech, dilution 1:100), anti-human lambda FITC (Southern Biotech, dilution 1:100), and PI (Invitrogen, dilution 1:100) for 20 minutes on ice. Yeast were again washed two times and samples were analyzed on a BD FACSCanto II flow cytometer (BD Biosciences).

### Hydrophobic interaction chromatography (HIC)

HIC assays were carried out as previously described^91^. Briefly, antibodies were diluted in a solution of 1.8 M ammonium sulfate and 0.1 M sodium phosphate pH 6.5 (phase A solution) to achieve a final concentration of 1.0 M ammonium sulfate. A linear salt gradient from phase A solution to the same solution without ammonium sulfate (phase B solution) was set up on a Sepax Proteomix HIC butyl-NP5 column; the gradient was run for 20 minutes at a flow rate of 1.0 mL/min. The UV absorbance at 280 nm was monitored to obtain peak retention times.

### Thermostability assay by differential scanning fluorescence (DSF)

Thermal melting (T_m_) measurements of the Fabs were carried out as previously described^92^. Briefly, 20 µL of 1 mg/mL antibody sample was mixed with 10 µL of 20X SYPRO orange. The CFX Real-Time System (BioRad) was used to scan the plate from 40–95 °C at a rate of 0.25 °C/min. Subsequently, BioRad analysis software was used to calculate T ^App^ from the primary derivative of the raw data.

### Generation of tecVLPs bearing CCHFV IbAr10200 GPC

The amino acid sequence for the IbAr10200 GPC was derived from GenBank M-segment sequences with an accession number NC_005300. Transcription- and entry-competent virus-like particles (tecVLPs) were generated as described previously^30,45^. Briefly, BSR-T7 cells were transfected with plasmids encoding the T7 polymerase, a minigenome expressing Nano-Glo Luciferase, and the CCHFV nucleoprotein (NP), glycoprotein complex (GPC), and polymerase (L). 15 hours post-transfection, transfection medium was removed and replaced with fresh DMEM growth media. 48 hours post-transfection, tecVLP-containing supernatants were collected, clarified by low-speed centrifugation, and pelleted by ultracentrifugation at 25,000 x *g* for 2.5 hours. Pelleted tecVLPs were resuspended in DMEM overnight and stored at −80 °C overnight prior to use.

### Neutralization assays against IbAr10200 tecVLPs

Neutralization by candidate mAbs against CCHFV IbAr10200 tecVLPs were assessed in Vero cells, maintained as described above and previously^30^. In brief, antibodies were diluted to starting concentrations of 350 nM (anti-GP38 mAbs) or 100 nM (anti-Gc mAbs) and subsequently serially diluted 3-fold in complete DMEM. TecVLPs, at an amount empirically determined such that the luciferase signal in target cells was approximately 500-fold over background, were then incubated with antibodies for one hour at 4 °C. After one hour, antibody/tecVLP mixtures were added to Vero cells in triplicate and incubated for 16 hours. Following infection, luciferase signal was assayed using Nano-Glo Luciferase assay system (Promega) and the signal for each mAb tested was normalized to a no-antibody control.

### Neutralization assays against authentic CCHFV

Neutralization assays were conducted similarly to what was described previously, with modifications^28,30^. Briefly, CCHFV-IbAr10200, CCHFV-Afg09, CCHFV-Turkey2004, or CCHFV-Oman were incubated with serial 3-fold dilutions of mAbs (at a starting concentration of 500 nM) for 1 hour at 37 °C. The antibody-virus mixture was added to monolayers of VeroE6 or SW-13 cells in a 96-well plate at a final multiplicity of infection of 1 (IbAr10200 and Afg09) or 0.3 (Turkey2004 and Oman) and incubated for one hour at 37 °C. Infection medium was then removed, and fresh cell culture medium without mAb was added. 24 (IbAr10200 and Afg09) or 48 hours (Turkey2004 and Oman) post infection, culture medium was removed, and plates were submerged in 10% formalin and plates were fixed for at least 24 hours at 4 °C. Plates were removed from formalin and permeabilized with 0.2% Triton-X for 10 minutes at room temperature and treated with blocking buffer (5% milk). Infected cells were detected by consecutive incubation with CCHFV-specific antibody 9D5 (3 µg/ml; BEI NR-40270) and secondary detection antibody (goat anti-mouse) conjugated to AlexaFluor 488 (1:2000 dilution; Invitrogen). Percent infection was determined using the Cytation5 high-content imaging instrument and data analysis was performed using the or Gen5.11 software (BioTek).

### Murine challenge studies

#### Therapeutic IbAr10200 study

5–8-week-old male and female IFNAR^−/−^ mice (Charles River) were exposed intraperitoneally (IP) to 100 PFU of CCHFV-IbAr10200. Mice were treated IP with 1 mg of indicated mAb, or an equivalent volume (200 µl) of phosphate-buffered saline (PBS) vehicle 24 hours (+1 day) and 96 hours (+4 day) post-exposure, for a total of 2 mg of mAb per mouse. Animals were observed daily for clinical signs of disease and morbidity for 28 days. Mice were scored on a 4-point grading scale, where a 1 was defined by decreased grooming and ruffled fur, a 2 defined by subdued behavior when un-stimulated, a 3 defined by lethargy, hunched posture, and subdued behavior even when stimulated, and a 4 defined by bleeding, unresponsiveness, severe weakness, or inability to walk. Mice scoring a 4 were considered moribund and were humanely euthanized based on IACUC-approved criteria. Daily observations were increased to a minimum of twice daily while mice were exhibiting clinical signs of disease (clinical score = 3).

#### Therapeutic Afg09, Oman, and Turkey2004 study

3–8-week-old male and female STAT1^−/−^ mice (The Jackson Laboratory) were exposed IP to 100 PFU of CCHFV-Afg09 or 1000 PFU of CCHFV-Turkey2004 or CCHFV-Oman. For the second challenge study (**Supplementary Figure S16**), mice were either treated IP with 1 mg of indicated mAb, or an equivalent volume (200 µl) of PBS vehicle 24 hours (+1 day) and 96 hours (+4 day) post-exposure, for a total of 2 mg of mAb per mouse. For the third challenge study (**Figure 7**), mice were treated IP with 0.2 mg of indicated mAb or an equivalent volume (200 µl) of PBS vehicle 30 minutes post-exposure. Animals were observed daily for clinical signs of disease and morbidity for 28 days. Mice were scored on a 4-point grading scale as described above. Daily observations were increased to a minimum of twice daily while mice were exhibiting signs of disease (clinical score = 3). Mice scoring a 4 were considered moribund and were humanely euthanized based on IACUC-approved criteria.

### Quantification and statistical analysis

Statistical details, including the number of replicates (n), measures of precision, and the statistical test used for each experiment can be found in the corresponding figure legends and in the results section. All statistical analyses were conducted in GraphPad Prism.

---

**Supplementary Figure S1.**
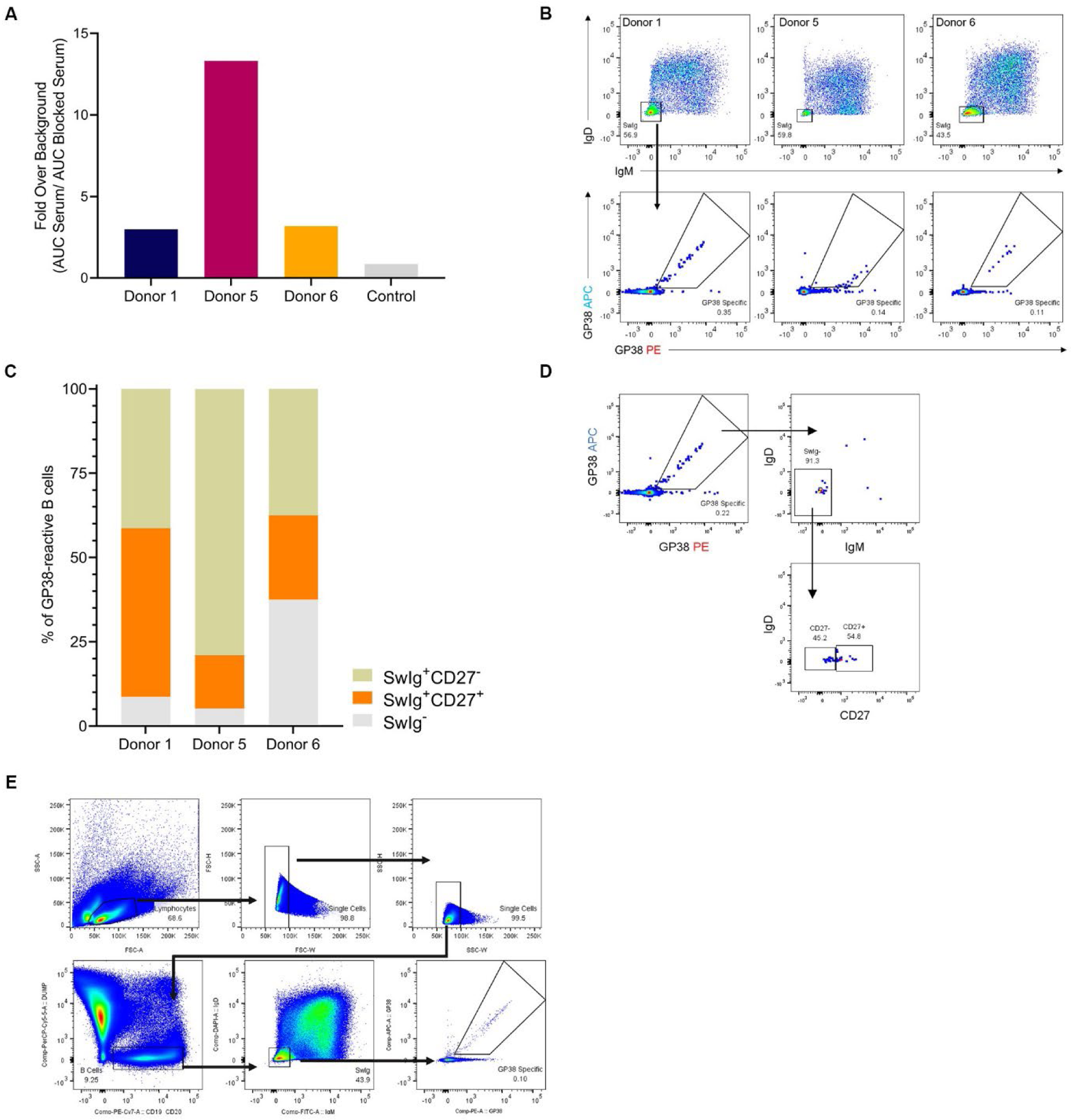
Serum analysis and flow cytometry. (A) ELISA data from donor and control serum, reported as fold-over-background of area under the curve (AUC) of the serum/blocked serum. (B) Flow cytometric analysis of IgM and IgD surface expression of donor B cells (top panel) and avid-rGP38 binding of SwIg B cells (bottom panel). Donor 1 PBMCs were gated on CD3^−^CD8^−^CD14^−^CD16^−^PI^−^CD19^+^ lymphocytes; Donors 5 and 6 PMBCs were gated on CD3^−^CD8^−^CD14^−^CD16^−^PI^−^CD19^+^CD20^+^ lymphocytes. (C) Bar chart of CD27 surface expression on rGP38-reactive SwIg B cells broken down by donor. (D) Representative gating strategy used for the calculations in panel C. Upstream of the first flow plot, PBMCs were gated on CD3^−^CD8^−^ CD14^−^CD16^−^PI^−^CD19^+^ lymphocytes. (E) Example of gating strategy used to sort rGP38-reactive, SwIg B cells.

**Supplementary Figure S2.**
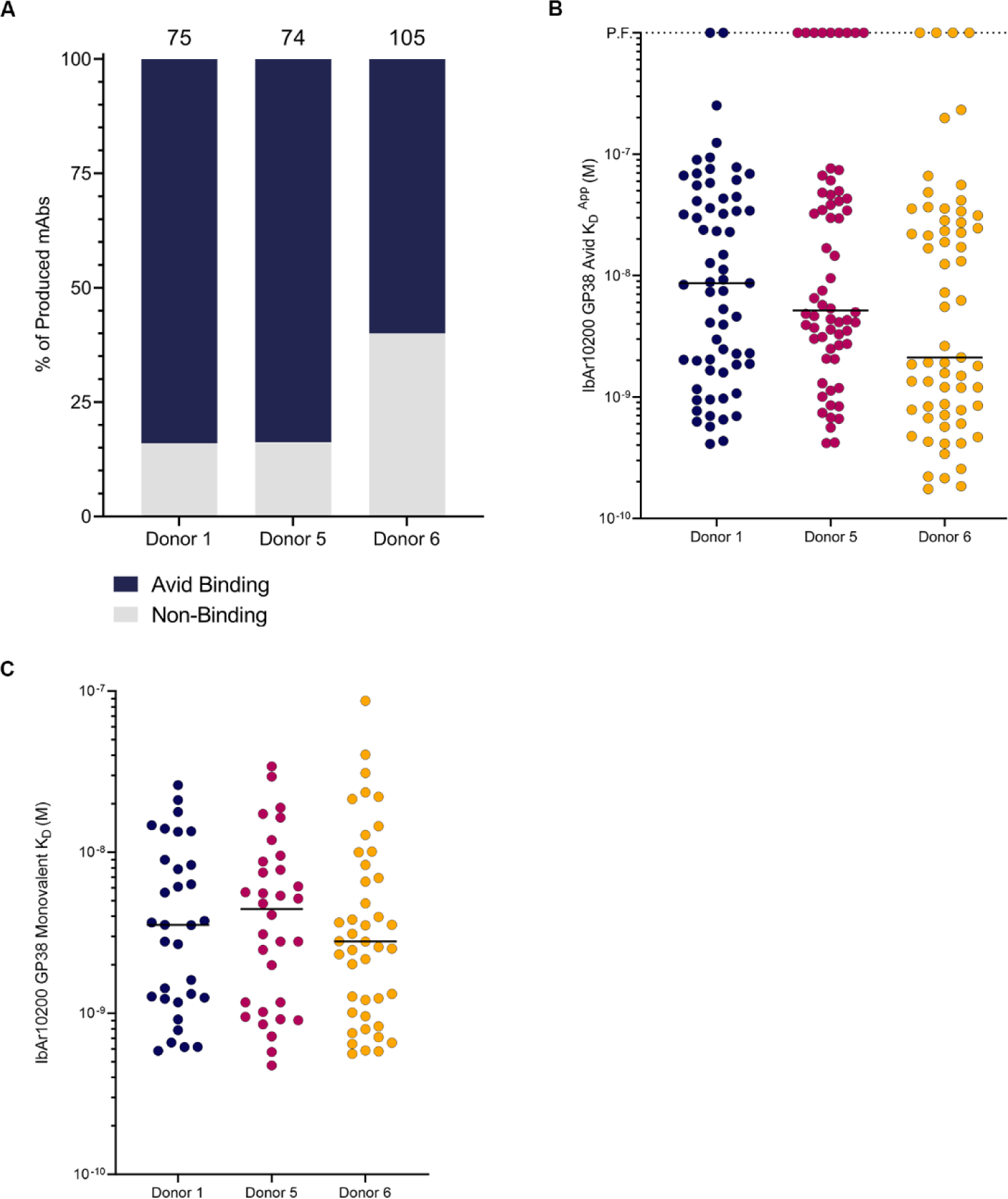
Avid binding analysis of antibodies from the three donors. (A) Avid binding of the produced mAbs to IbAr10200 rGP38 protein. Total number of mAbs from each donor are indicated above each representative bar. (B) Avid K_D_apparent (K_D_^App^) of the mAbs that bound avidly to IbAr10200 rGP38 in the assay shown in panel A. P.F. indicates poor fit of the BLI curve. (C) Single concentration monovalent K_D_ of the mAbs that bound monovalently to IbAR10200 rGP38 shown in **Figure 1B** with poor fitting samples removed for analysis. Black horizontal line defines median.

**Supplementary Figure S3.**
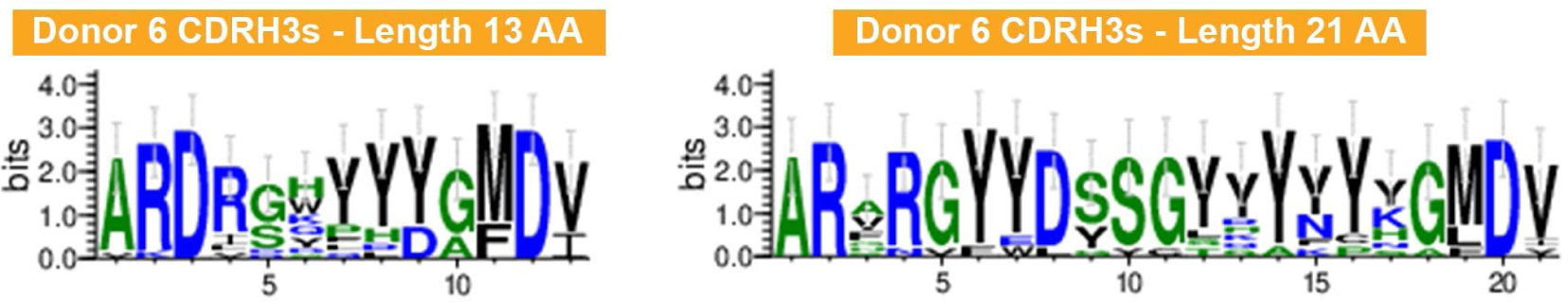
Logo plot. Logo plot representing CDRH3 sequences of mAbs cloned from Donor 6 B cells that have a length of 13 and 21 amino acids (AA), respectively. Hydrophilic amino acids (R, K, D, E, N, Q) are colored in blue; neutral amino acids (S, G, H, T, A, P) are colored in green; hydrophobic amino acids (Y, V, M, C, L, F, I, W) are colored in black. Logo plots were created using WebLogo software v3.5.0.

**Supplementary Figure S4.**
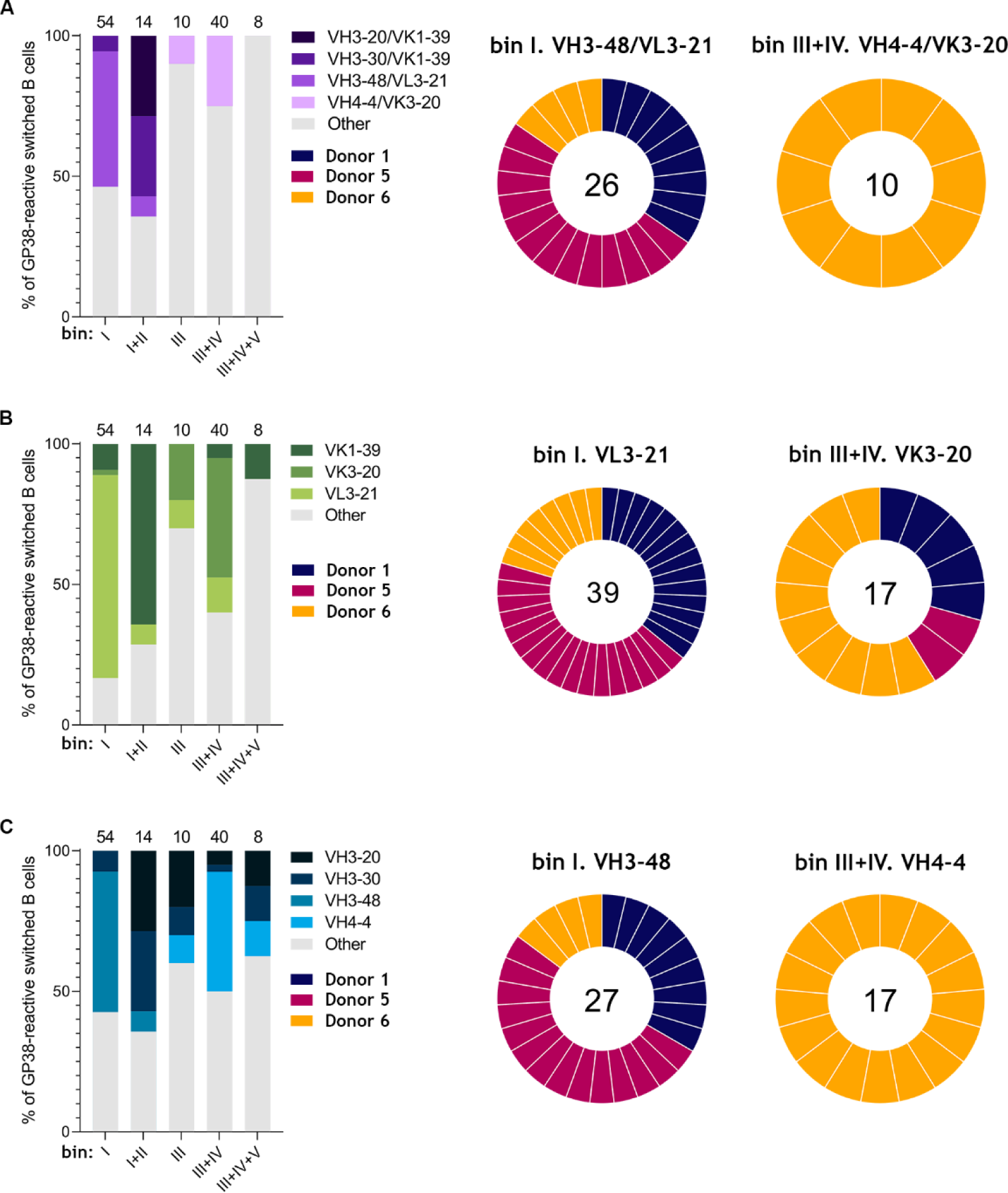
Germline gene usage per bin. (A) Germline gene pairing usage per bin (left); bin I VH3-48/VL3-21 and bin III+IV VH4-4/VK3-20 germline gene usage per donor (right). (B) Variable light chain germline gene usage per bin (left); bin I VL3-21 and bin III+IV VK3-20 germline gene usage per donor (right). (C) Variable heavy chain germline gene usage per bin (left); bin I VH3-48 and bin III+IV VH4-4 germline gene usage per donor (right). For all panels, the total number of mAbs per germline gene usage is indicated within each circular diagram.

**Supplementary Figure S5.**
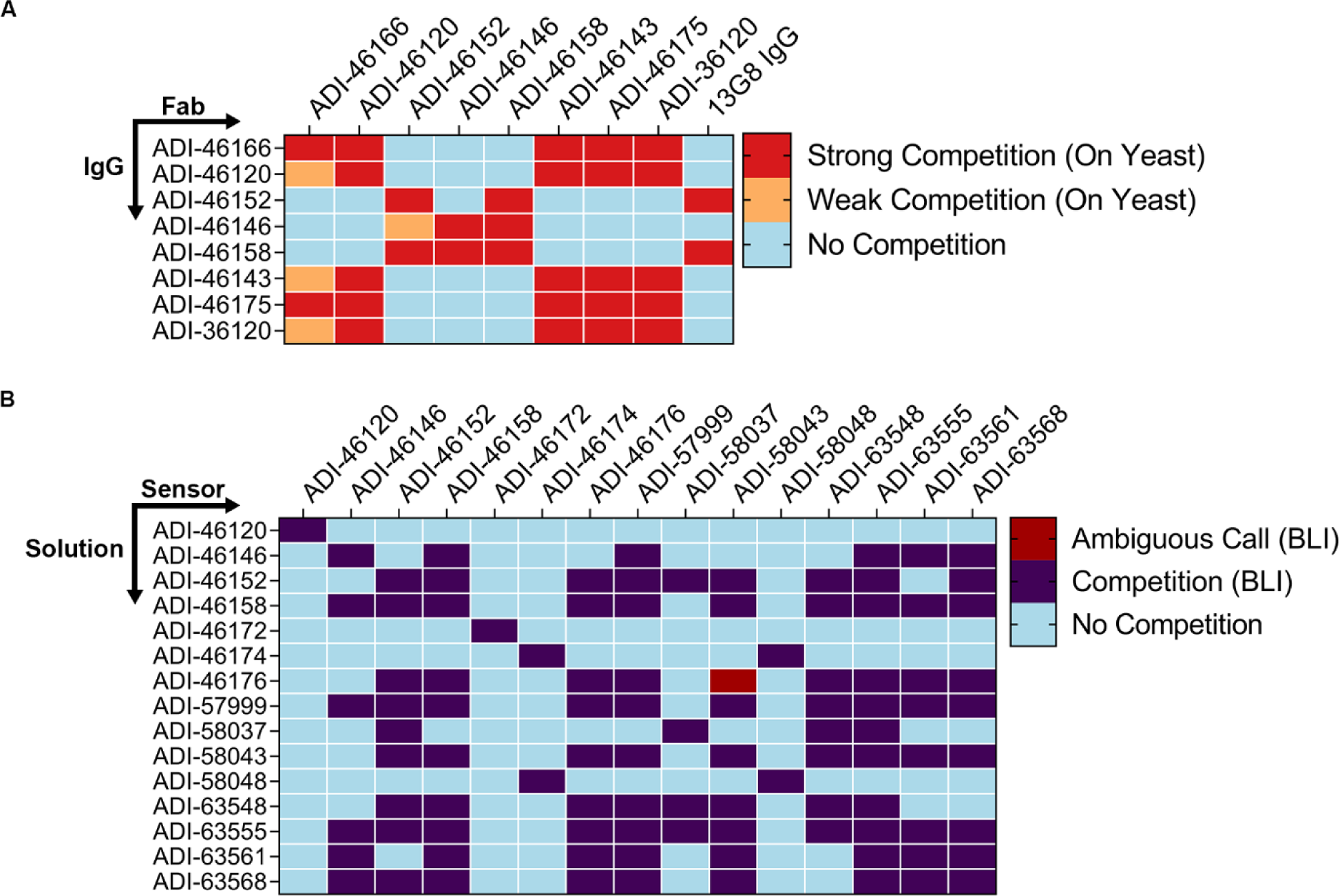
Preliminary cross-binning experiments. (A) Matrix of first preliminary cross-binning experiment. Results are displayed with surface-presented IgGs on the y-axis and competitive pre-complexed Fabs on the x-axis, unless otherwise noted. (B) Matrix of second preliminary cross-binning experiment. Results were run in an IgG vs. IgG format.

**Supplementary Figure S6.**
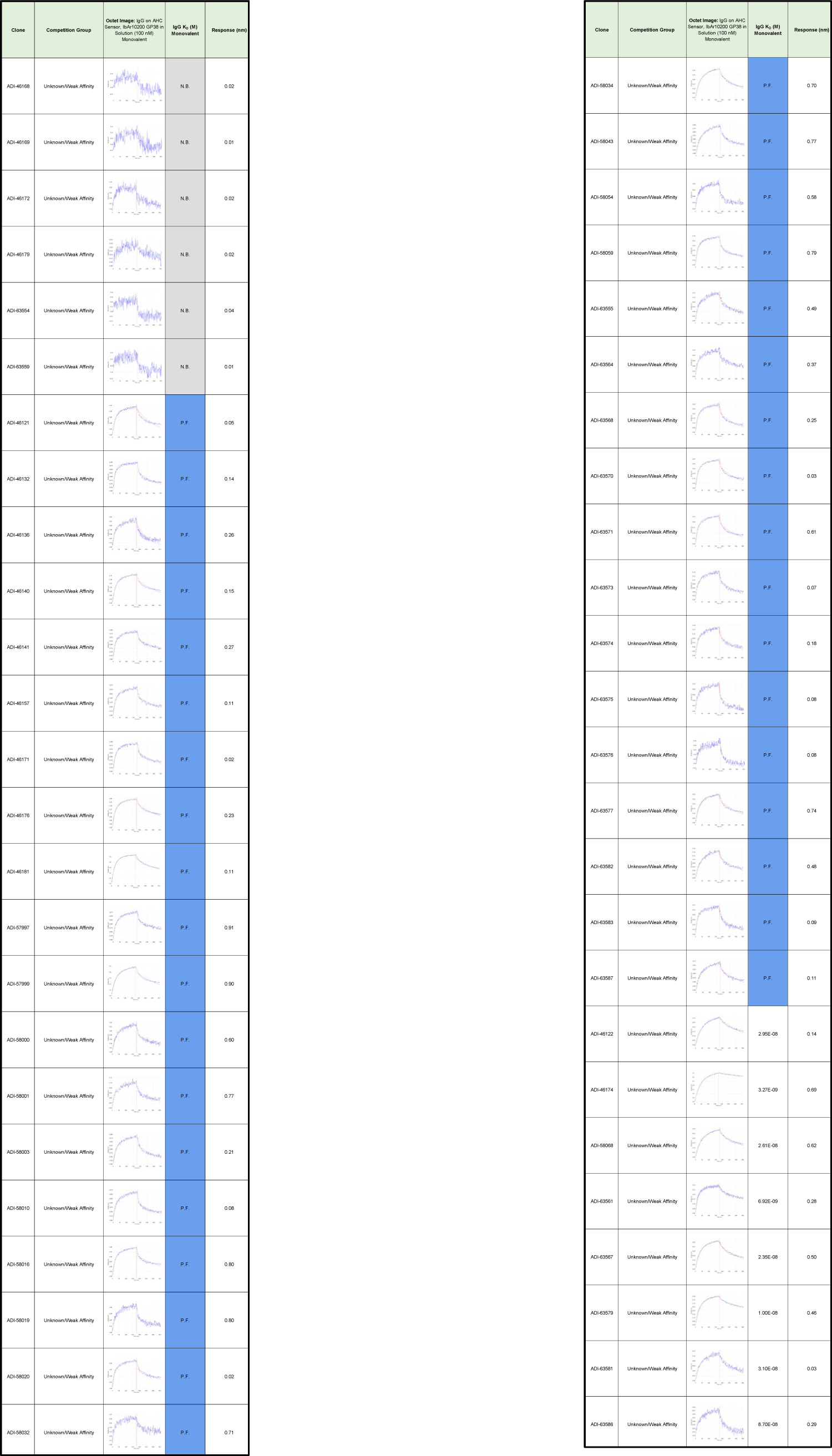
Antibody binding kinetics of the Unknown/Weak Affinity bin. Single concentration monovalent binding kinetics data for the antibodies in the Unknown/Weak Affinity bin. Columns from left to right: antibody name, competition group, ForteBio Octet trace, calculated monovalent K_D_ and response. For mAbs whose binding did not list a K_D_ the curves are denoted as either “P.F.” for poor fit for data to the binding model or “N.B.” for no binding.

**Supplementary Figure S7.**
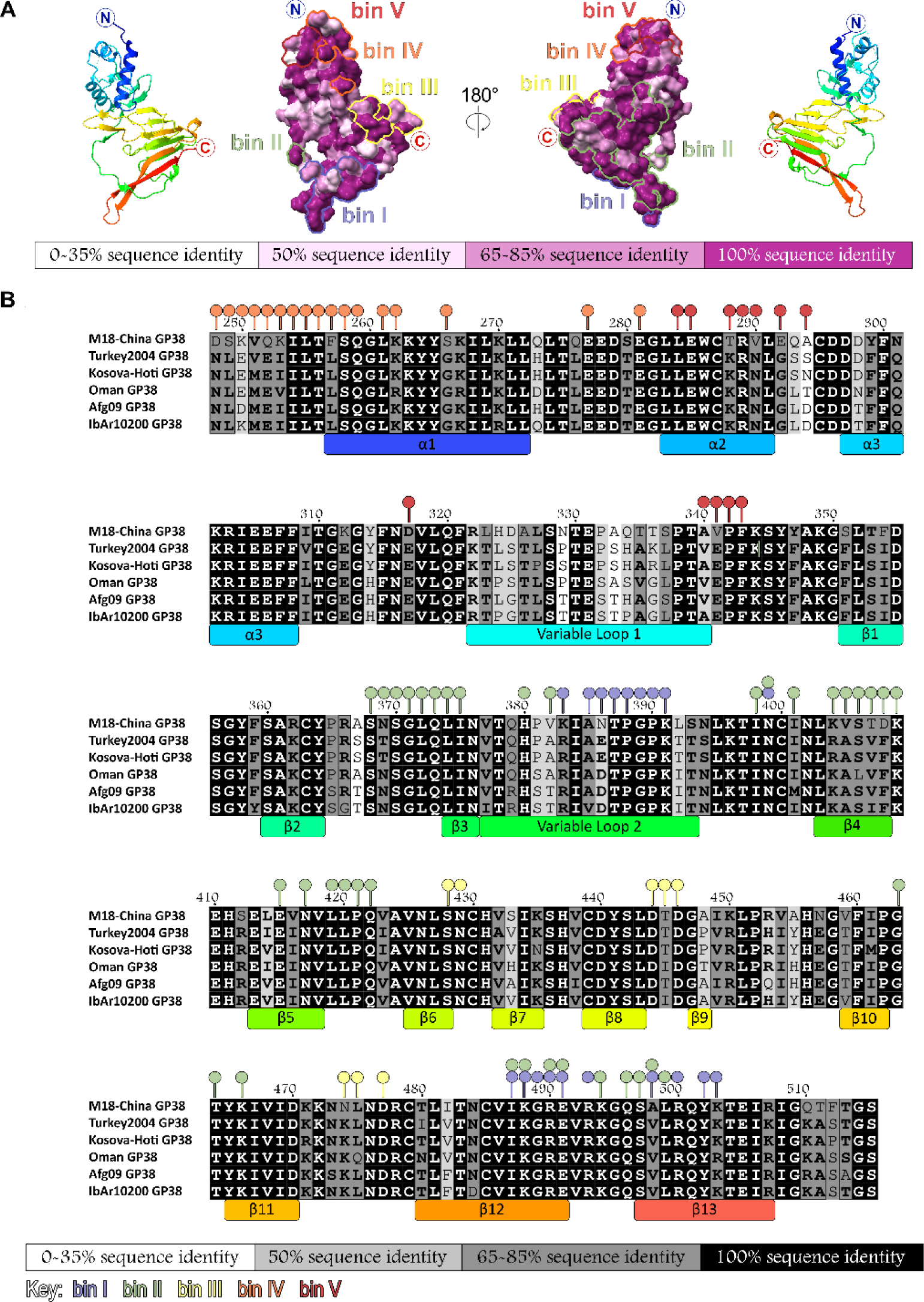
Conservation of CCHFV GP38 and interactions of GP38-specific antibodies. Regions of interactions for GP38-specific antibodies as identified from yeast surface display-based mapping (YSD residues that disrupted antibody binding by >75%) and high-resolution antibody structures (c13G8, ADI-46152, ADI-58048, ADI-46143) are displayed within the five main antigenic sites in corresponding colors for each bin: bin I (blue), bin II (green), bin III (yellow), bin IV (orange), and bin V (red). (A) Surface representation of IbAr10200 GP38 (PDB ID: 6VKF) colored by sequence identity for the six isolates using ChimeraX Color by Conservation: most variable residues (white) to most conserved residues (purple). (B) Sequence alignment generated by ClustalOmega for 79121M18 (UniProt: D4NYK3), 200406546-Turkey (UniProt: A0A0U2SQZ0), Kosova-Hoti (UniProt: B2BSL7), Oman-199809166 (UniProt: A0A0U3C6Q7), Afg09-2990 (UniProt: E5FEZ4), and IbAr10200 (UniProt: Q8JSZ3). Percent sequence identity indicated by box color: white (0-35%), light gray (50%), dark gray (65-85%), and black (100%). Sequence consensus or strong conservation among sequences is indicated by bold lettering. Secondary structure assigned for IbAr10200 GP38 (PDB 6VKF) from ESPript server with colored boxes from N-terminal (blue) to C-terminal (red).

**Supplementary Figure S8.**
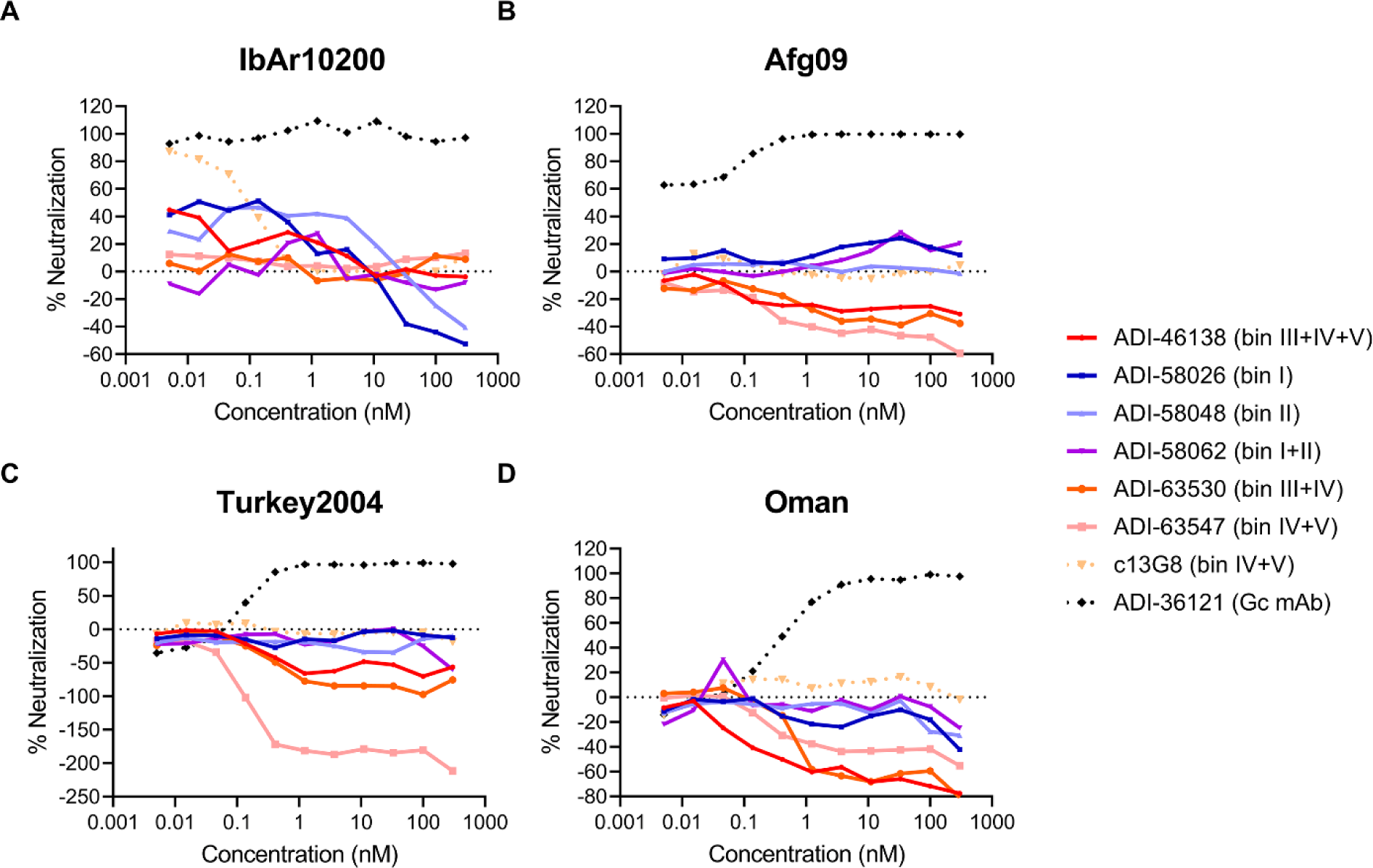
Authentic virus neutralization assay of GP38 mAb panel. (A-D) Neutralization curves of the indicated mAbs against authentic (A) CCHFV IbAr10200, (B) CCHFV Afg09, (C) CCHFV Turkey2004, and (D) CCHFV Oman. Neutralization assays were conducted in VeroE6 cells. The average of n=3 each from two independent experiments (n=6 total) is shown for all neutralization curves

**Supplementary Figure S9.**
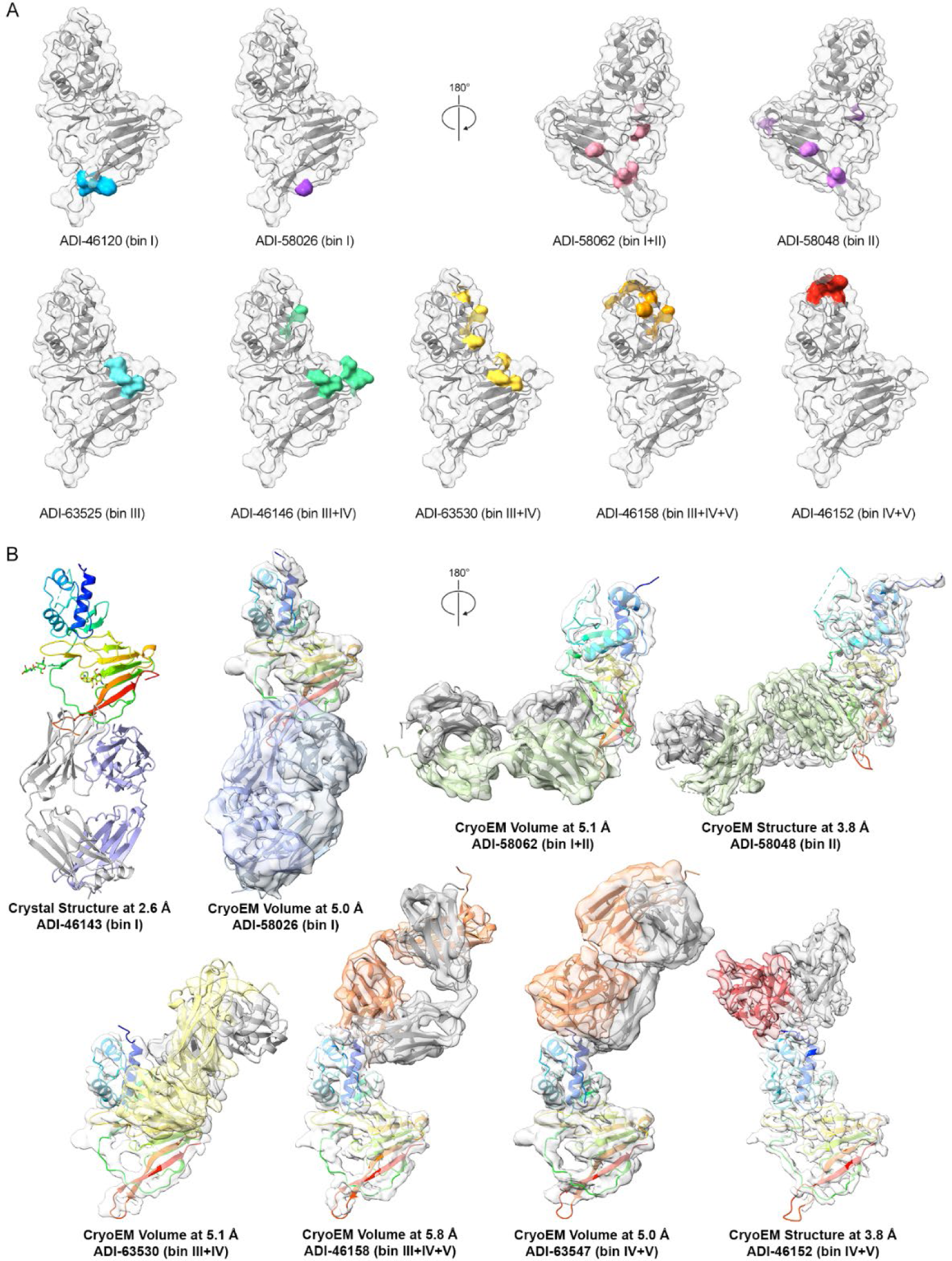
Structures and identified critical residues on GP38 required for antibody binding mapped across the surface of GP38. (A) Related to **Figure 5A**, yeast-based critical-residue mapping strategy revealed one to nine critical residues necessary for an antibody to bind to GP38. YSD residues are colored on the surface representation of CCHFV IbAr10200 GP38 (PBD ID: 6VKF, white surface). (B) GP38 bound ADI-46143 Fab crystal structure shown as ribbons. For the high-resolution cryo-EM structure of GP38+ADI-46152+ADI-58048, the refined model is docked into the cryo-EM map and displayed by each Fab. For the remaining medium-resolution cryo-EM structures, CCHFV IbAr10200 GP38 (PDB ID: 6VKF) and AlphaFold2 models are docked into the corresponding cryo-EM maps and displayed by each Fab. Full cryo-EM complexes (GP38 bound by both Fabs) are displayed in Supplementary Figures S10-S13.

**Supplementary Figure S10.**
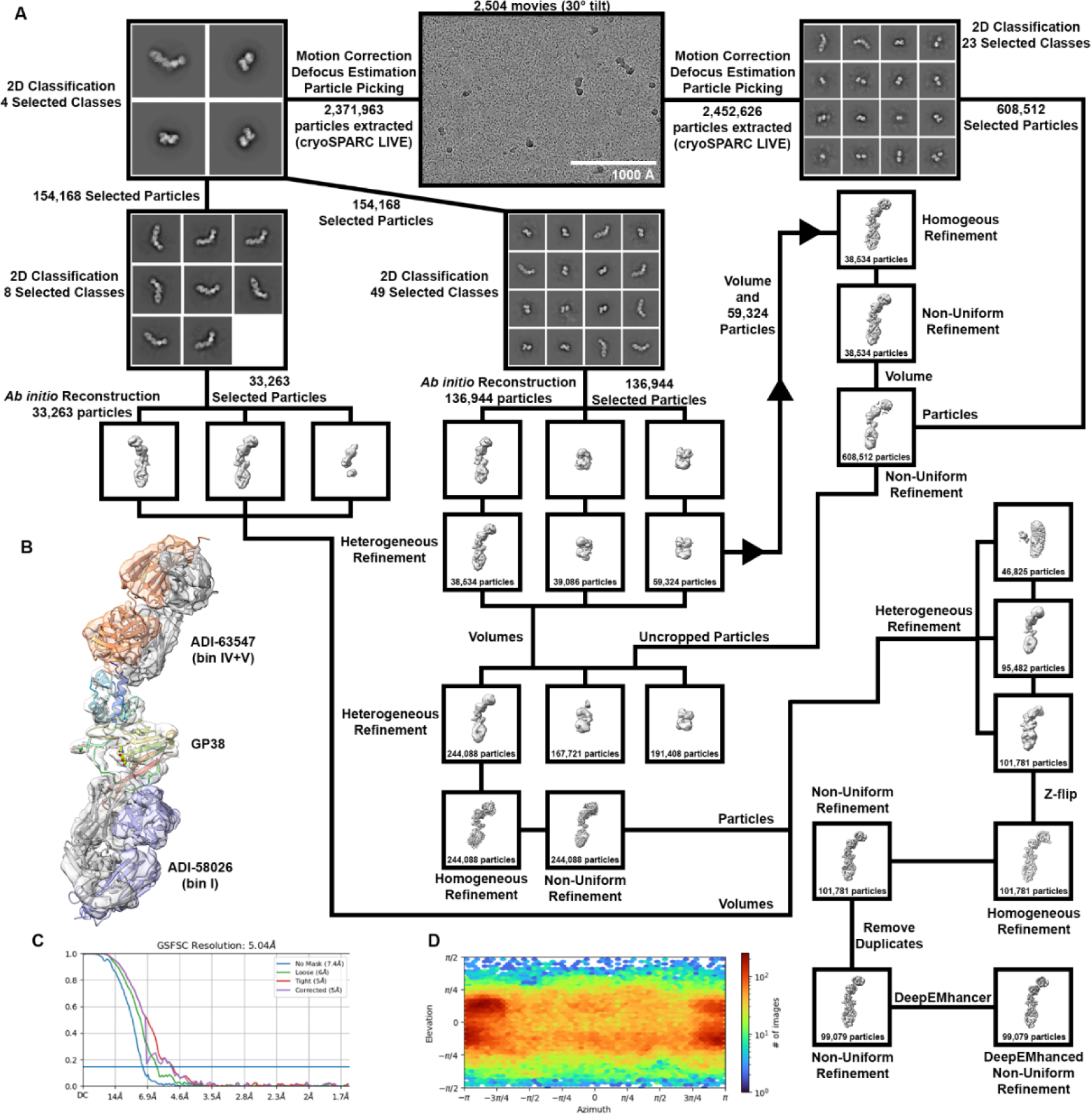
Structural characterization of ADI-58026 and ADI-63547 Fabs bound to CCHFV IbAr10200 GP38. (A) Data processing and refinement pipeline for the complex. Unless otherwise noted, processing was done using cryoSPARC v3.2 and subsequent versions. (B) Cryo-EM volume of the complex with docked models of GP38 (PDB:6VKF, rainbow) with ADI-58026 Fab (AlphaFold2 model, heavy chain in blue and light chain in dark gray) and ADI-63547 (AlphaFold2 model, heavy chain in orange and light chain in dark gray). (C) Gold standard FSC curve. (D) Viewing distribution plot.

**Supplementary Figure S11.**
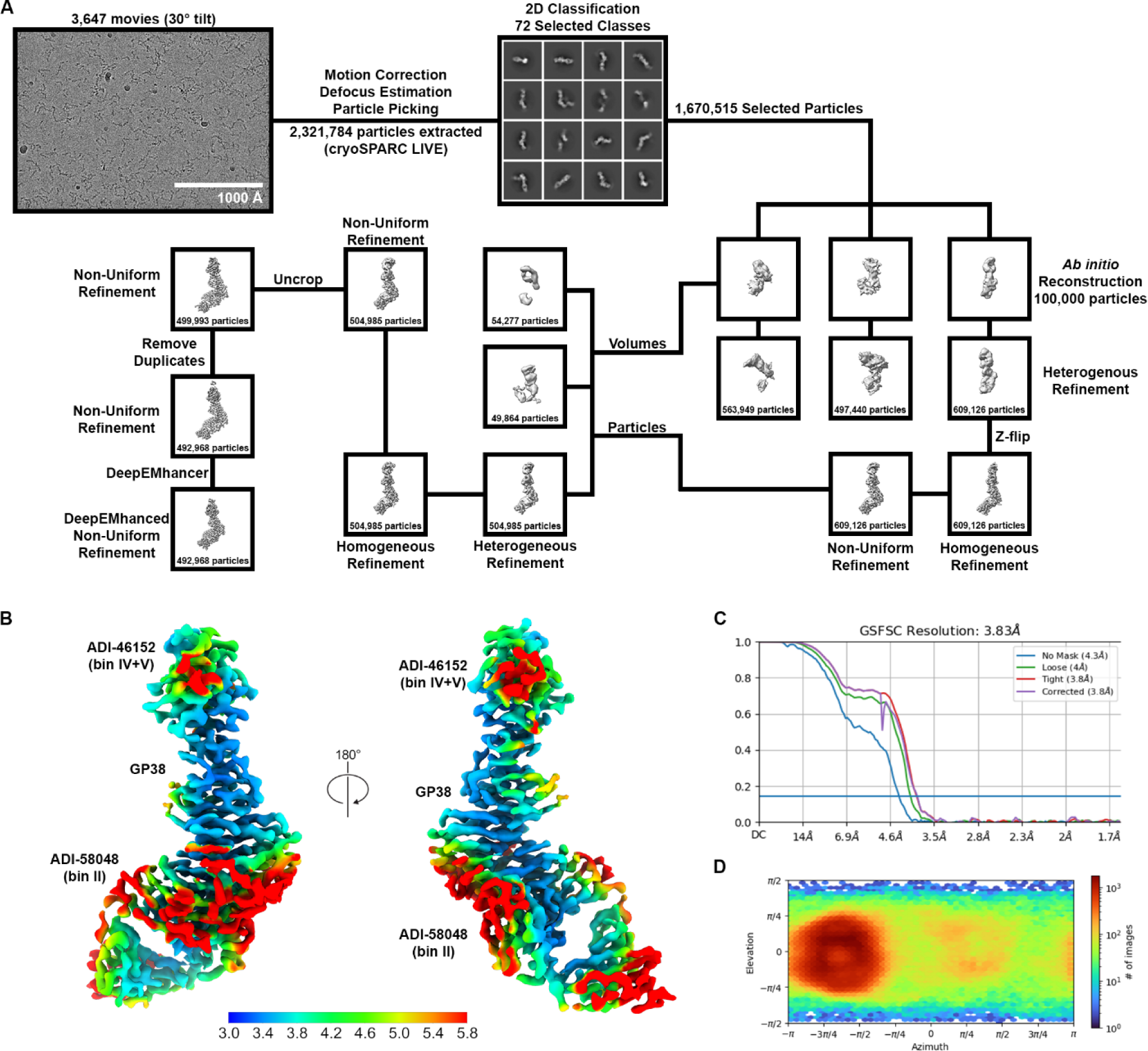
Structural characterization of ADI-46152 and ADI-58048 Fabs bound to CCHFV IbAr10200 GP38. (A) Data processing and refinement pipeline for the complex. Unless otherwise noted, processing was performed using cryoSPARC v3.2 and subsequent versions. (B) Local resolution estimation of the cryo-EM structure colored as a rainbow from blue (3.0 Å) to red (5.8 Å). (C) Gold standard FSC curve. (D) Viewing distribution plot.

**Supplementary Figure S12.**
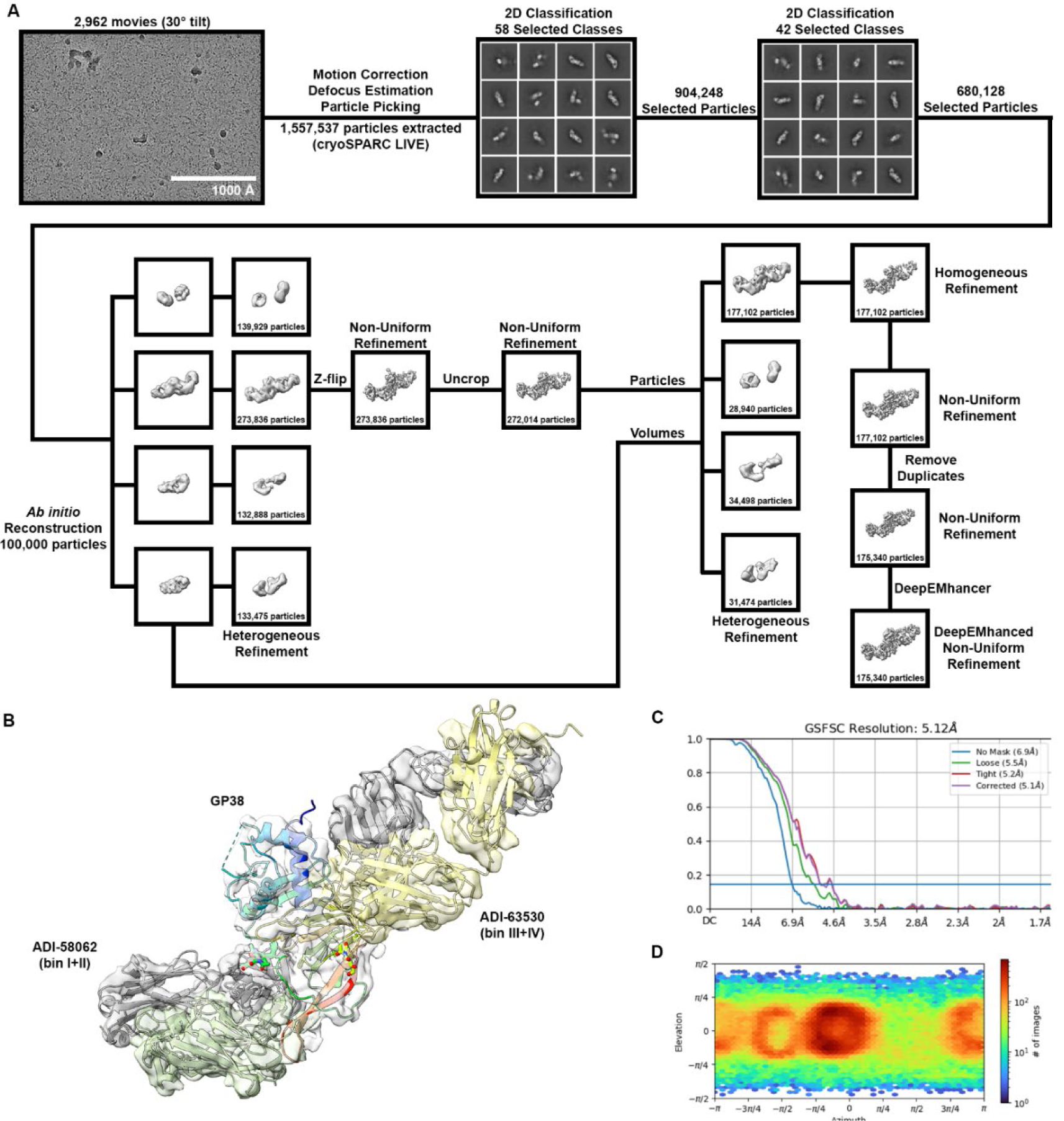
Structural characterization of ADI-58062 and ADI-63530 Fabs bound to CCHFV IbAr10200 GP38. (A) Data processing and refinement pipeline for the complex. Unless otherwise noted, processing was done using cryoSPARC v3.2 and subsequent versions. (B) Cryo-EM volume of the complex with docked models of GP38 (PDB ID: 6VKF, rainbow) with ADI-58062 Fab (AlphaFold2 model, heavy chain in green and light chain in dark gray) and ADI-63530 (AlphaFold2 model, heavy chain in yellow and light chain in dark gray). (C) Gold standard FSC curve. (D) Viewing distribution plot.

**Supplementary Figure S13.**
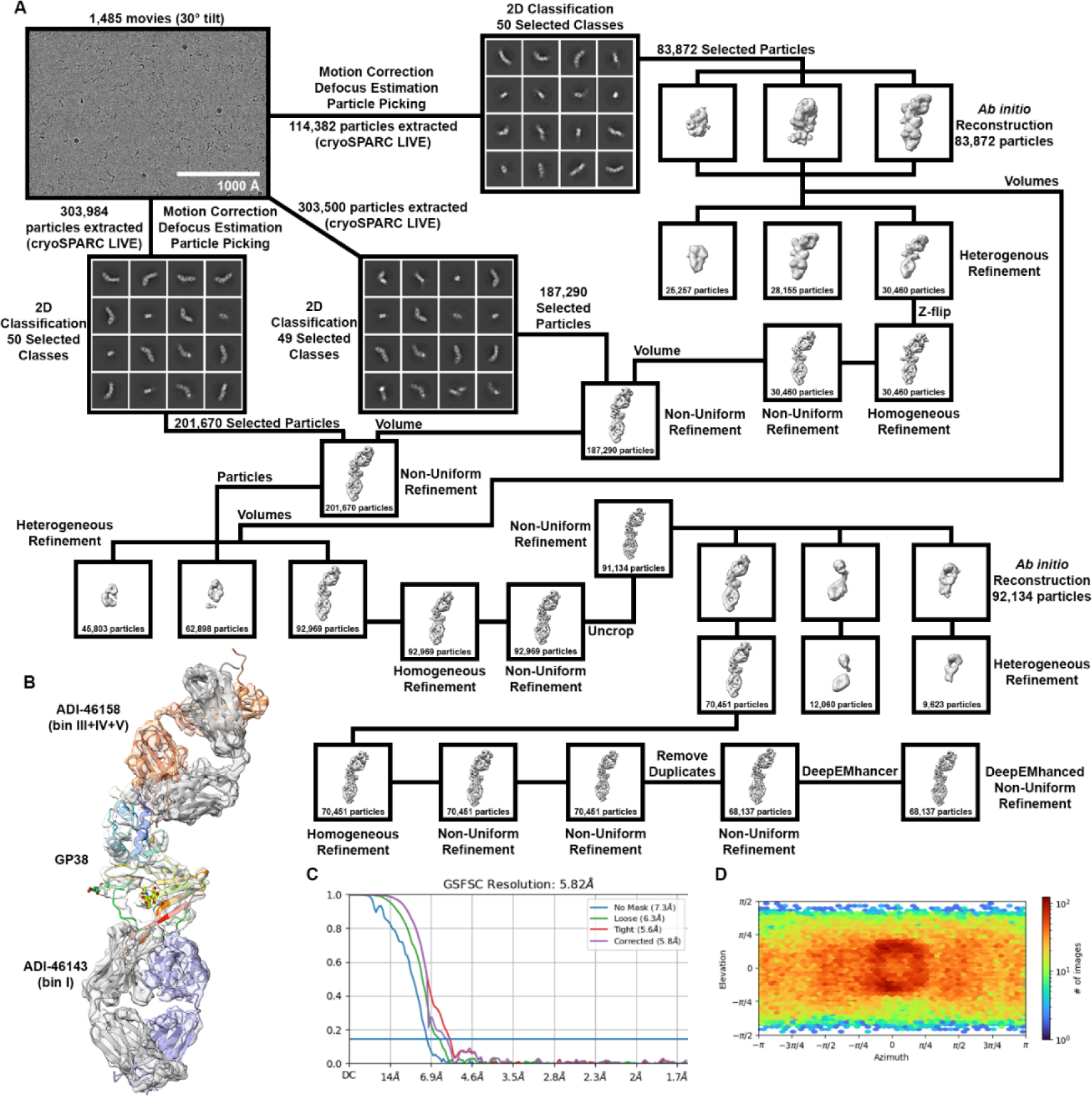
Structural characterization of ADI-46143 and ADI-46158 Fabs bound to CCHFV IbAr10200 GP38. (A) Data processing and refinement pipeline for the complex. Unless otherwise noted, processing was done using cryoSPARC v3.2 and subsequent versions. (B) Cryo-EM volume of the complex with docked models of GP38 (PDB ID:6VKF, rainbow) with ADI-46143 Fab (crystal structure, heavy chain in blue and light chain in dark gray) and ADI-46158 (AlphaFold2 model, heavy chain in orange and light chain in dark gray). (C) Gold standard FSC curve. (D) Viewing distribution plot.

**Supplementary Figure S14.**
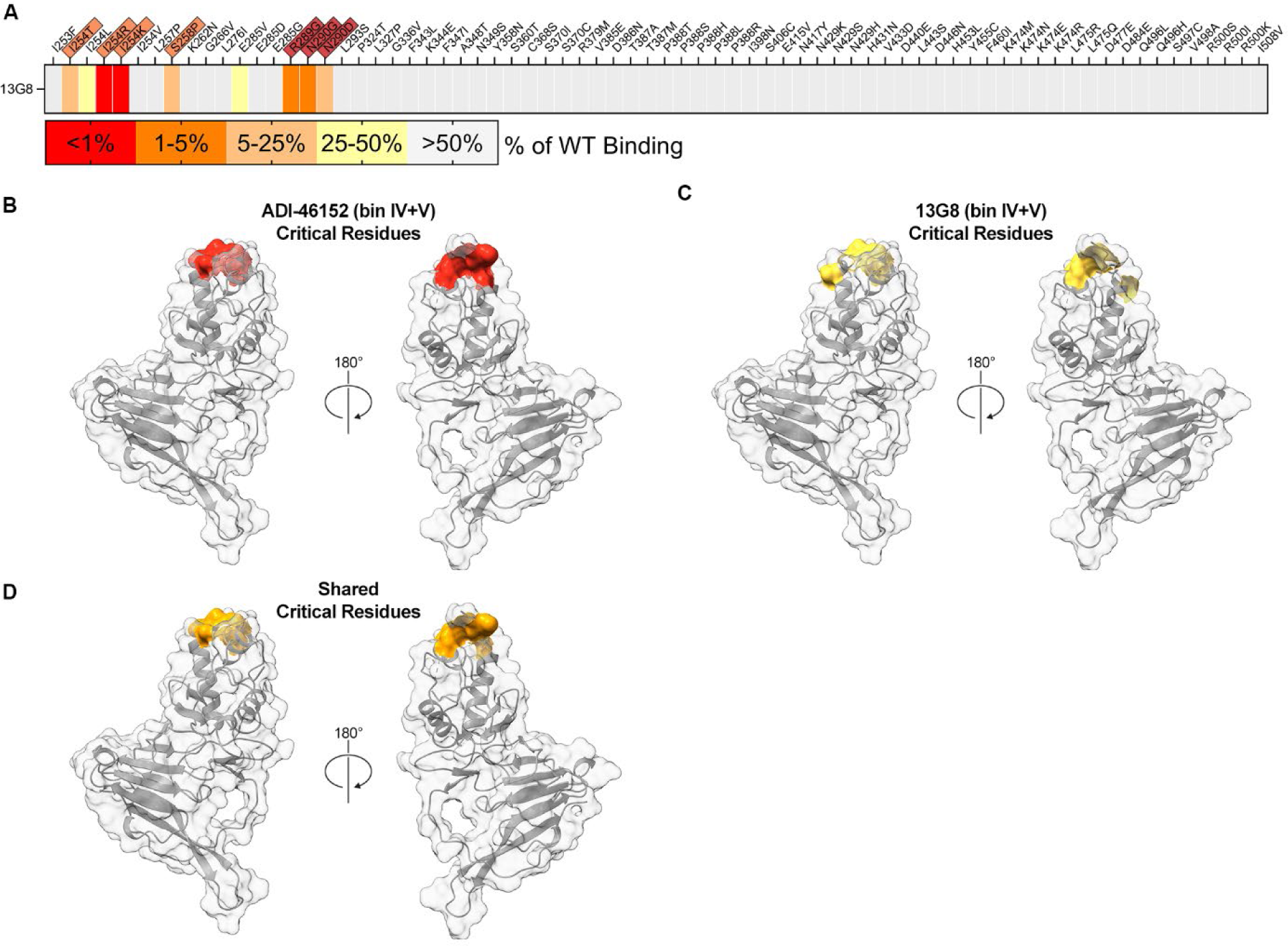
Identified critical residues for 13G8 binding to GP38. (A) Yeast-based critical-residue mapping strategy for 13G8. Total loss of binding with an identified residue (red), disruption of binding (orange to yellow), and majority of binding retained (gray). Identified critical residues on GP38 required for antibody binding overlaid onto the surface representation of GP38 (PDB ID: 6VKF) for (B) ADI-46152, (C) 13G8, and (D) shared critical residues between ADI-46152 and 13G8.

**Supplementary Figure S15.**
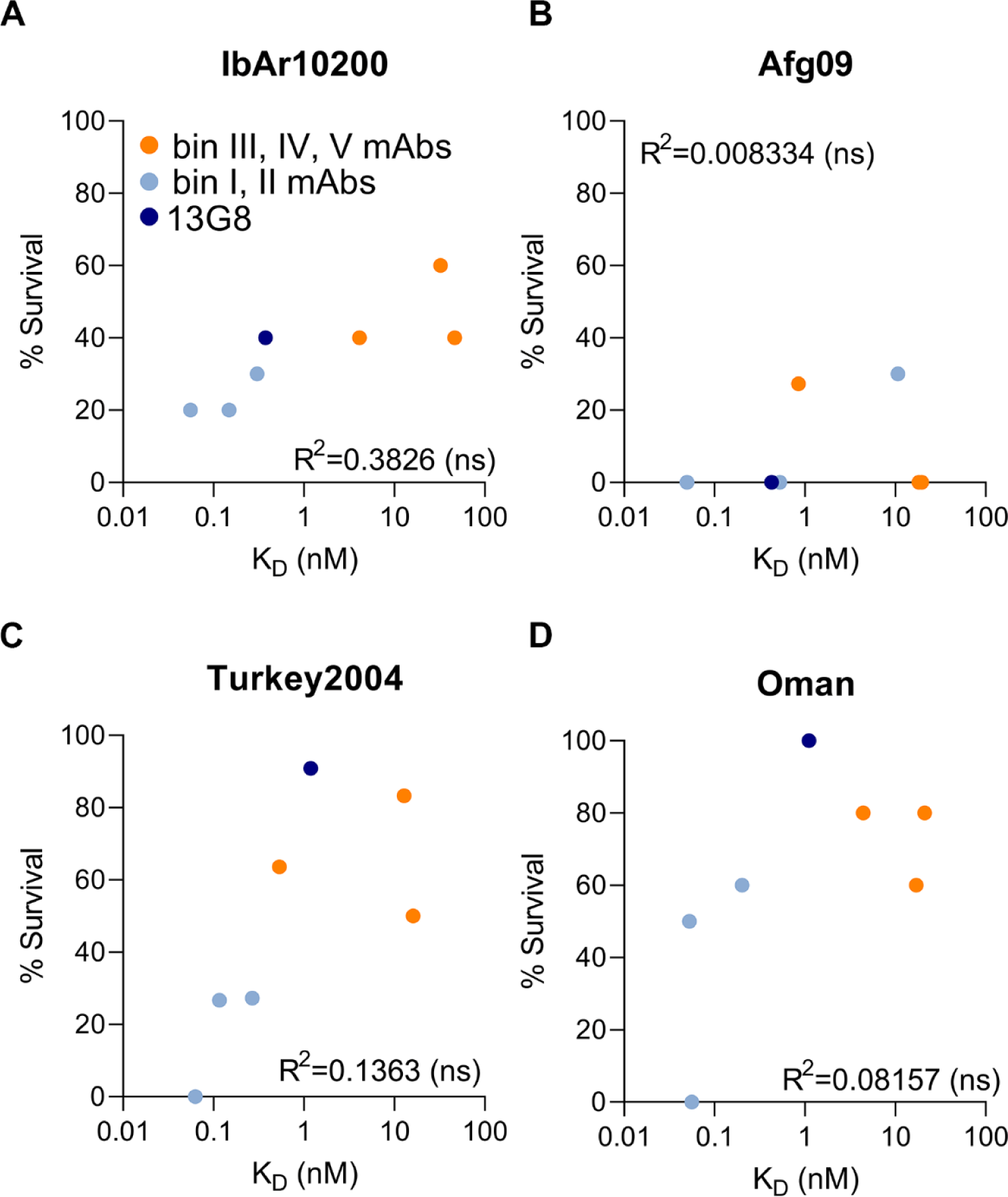
Percent survival of mice from experiments in **Figure 7** are plotted versus K_D_ determinations from SPR experiments in **Figure 3C** for each respective isolate (A, IbAr10200; B, Afg09-2990; C, Turkey2004; D, Oman). mAbs that are bin III and/or IV and/or V competitors are colored orange and mAbs that are bin I and/or II competitors are colored light blue. 13G8 is colored navy. R-squared calculated by Spearman’s correlation coefficient. ns is non-significant.

**Supplementary Figure S16.**
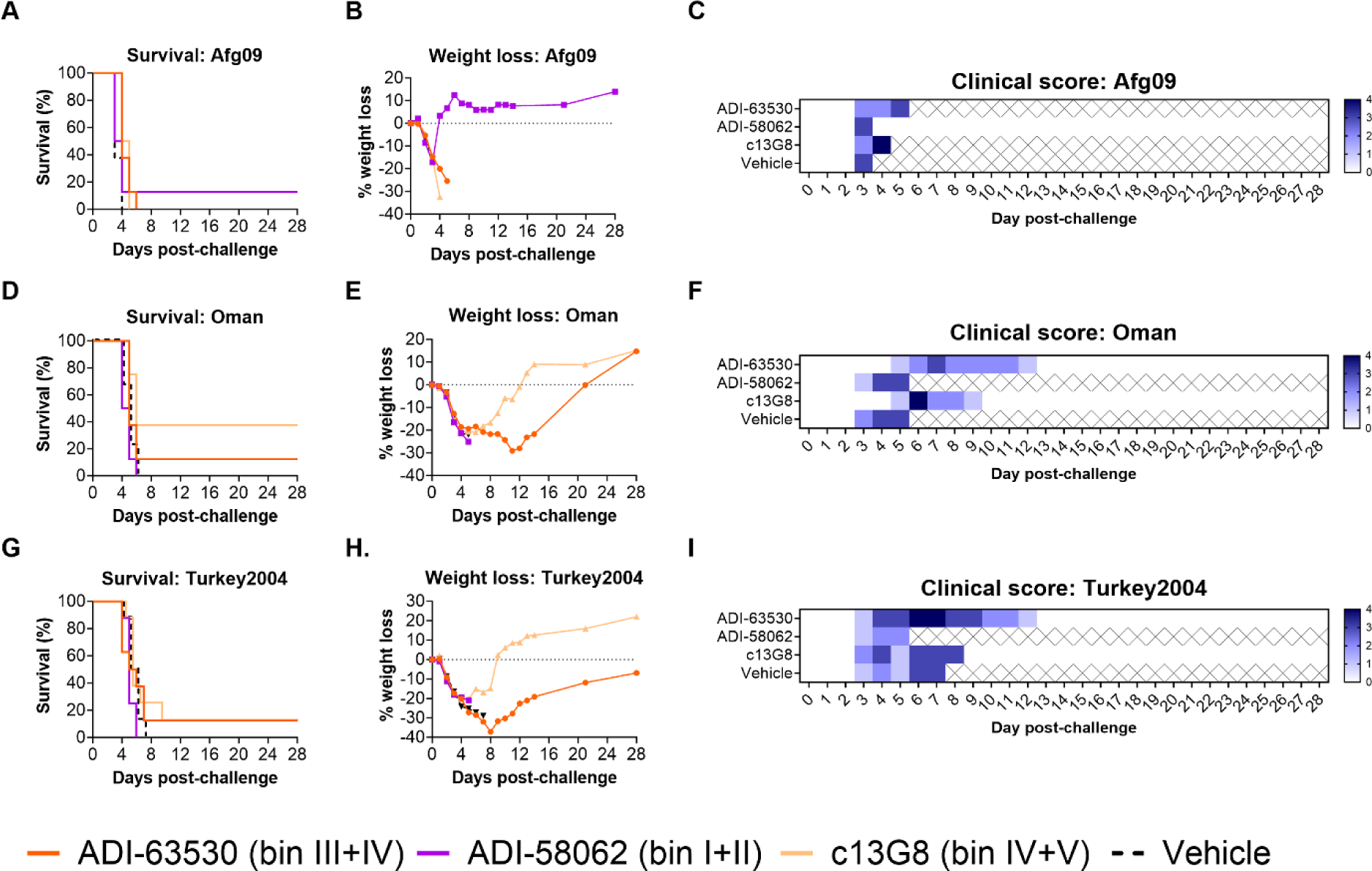
Cross-clade therapeutic efficacy of lead GP38 mAbs. STAT1^−/−^ mice were challenged with (A-C) CCHFV-Afg09, (D-F) CCHFV-Oman, or (G-I) CCHFV-Turkey2004 and then treated with 1 mg/mouse of mAb or vehicle 1- and 4-days post-challenge (2 mg total; n=8 mice per group). (A, D, G) Survival curves, (B, E, H) associated mean weight loss, and (C, F, I) clinical score data are shown.

**Supplementary Table S1.** Bin code and representative antibody table. Antibodies representative of each of the discrete antigenic sites (bold font) along with the overlapping bins (non-bold font).

| **Bin Code** | **Competing Antibodies** |
| --- | --- |
| **I** | **ADI-46120** |
| I + II | ADI-46120 & ADI-58048 |
| I + III | ADI-46120 & ADI-46146 |
| I + IV | ADI-46120 & ADI-46158 |
| **II** | **ADI-58048** |
| **III** | **ADI-46146** |
| III + IV | ADI-46146 & ADI-46158 |
| III + IV + V | ADI-46146 & ADI-46158 & ADI-46152 |
| **IV** | **ADI-46158** |
| IV + V | ADI-46158 & ADI-46152 |
| **V** | **ADI-46152** |
| Unknown/Weak Affinity | NA |

**Supplementary Table S2.**
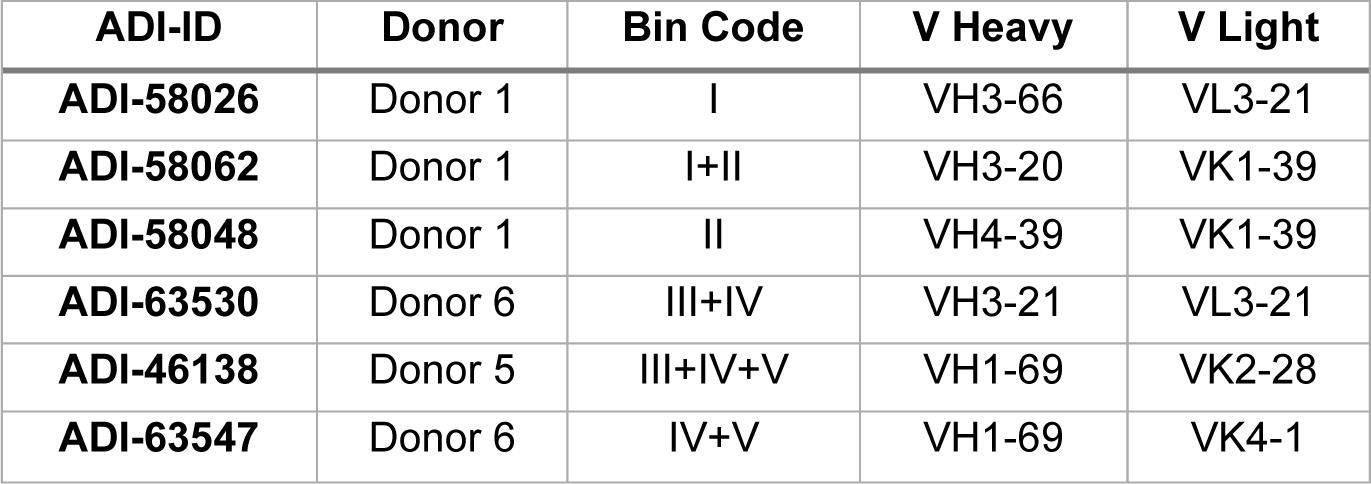
Germline gene usage of mAbs used in protection studies. Variable heavy chain and variable light chain gene information for mAbs selected for protection studies.

| **ADI-ID** | **Donor** | **Bin Code** | **V Heavy** | **V Light** |
| --- | --- | --- | --- | --- |
| **ADI-58026** | Donor 1 | I | VH3-66 | VL3-21 |
| **ADI-58062** | Donor 1 | I+II | VH3-20 | VK1-39 |
| **ADI-58048** | Donor 1 | II | VH4-39 | VK1-39 |
| **ADI-63530** | Donor 6 | III+IV | VH3-21 | VL3-21 |
| **ADI-46138** | Donor 5 | III+IV+V | VH1-69 | VK2-28 |
| **ADI-63547** | Donor 6 | IV+V | VH1-69 | VK4-1 |

**Supplementary Table S3.**
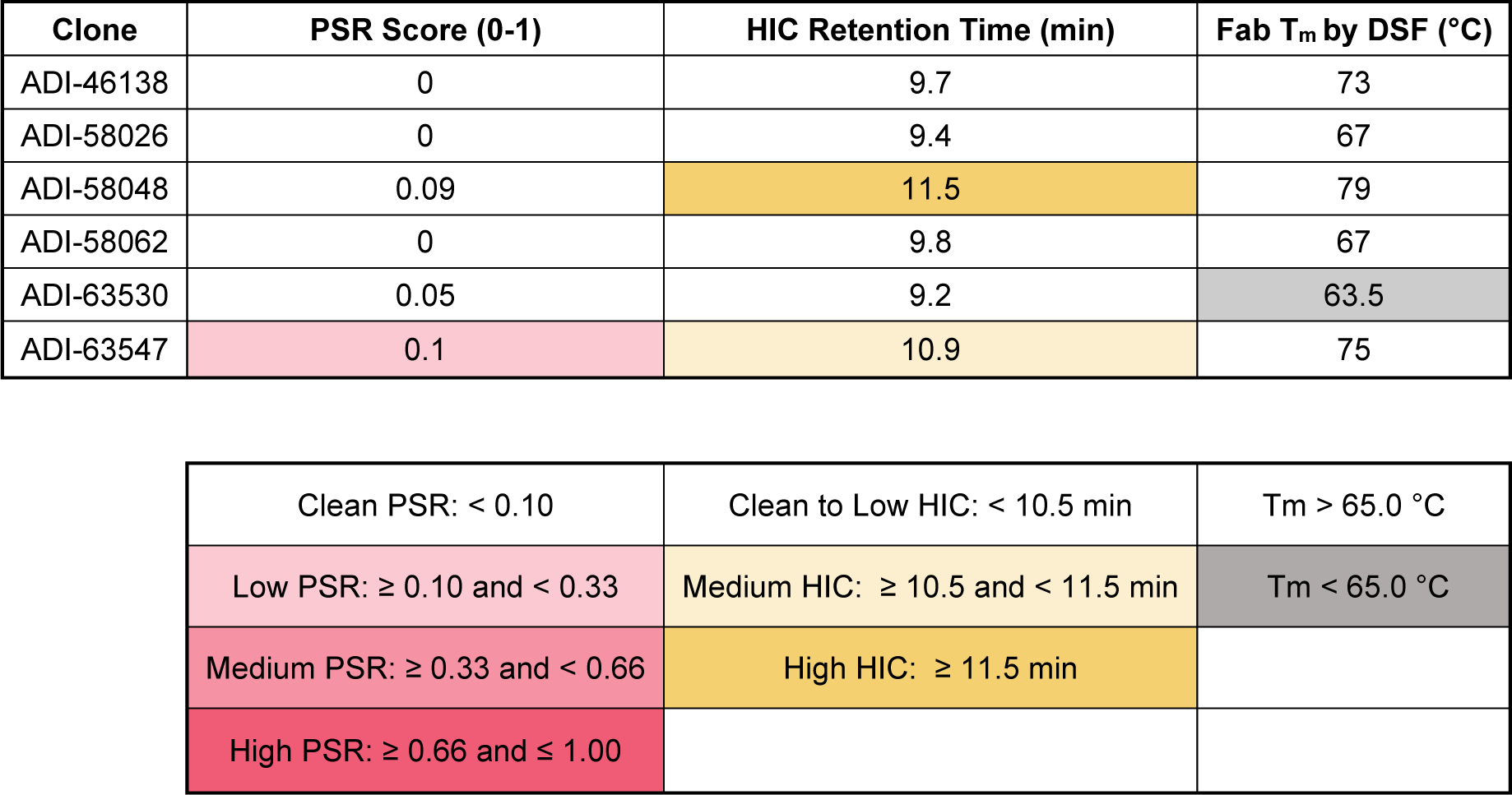
Developability metrics for the six mAbs used in protection studies. Table describing the developability properties of lead mAbs^1^. Poly-Specificity Reagent (PSR) indicates relative level of poly-specificity in each mAb normalized against standard control IgGs. Hydrophobicity Interaction Chromatography (HIC) measures mAb interaction with a HIC column as a normalized time to elution off the column. Fab Tm provides a measure of antibody thermostability using differential scanning fluorimetry (DSF) and is reported as the lowest temperature event distinct from a constant-heavy-2 (CH2) signal.

| Clone | PSR Score (0-1) | HIC Retention Time (min) | Fab T<sub>m</sub> by DSF (°C) |
| --- | --- | --- | --- |
| ADI-46138 | 0 | 9.7 | 73 |
| ADI-58026 | 0 | 9.4 | 67 |
| ADI-58048 | 0.09 | 11.5 | 79 |
| ADI-58062 | 0 | 9.8 | 67 |
| ADI-63530 | 0.05 | 9.2 | 63.5 |
| ADI-63547 | 0.1 | 10.9 | 75 |

|  |  |  |
| --- | --- | --- |
| Clean PSR: < 0.10 | Clean to Low HIC: < 10.5 min | T<sub>m</sub> > 65.0 °C |
| Low PSR: ≥ 0.10 and < 0.33 | Medium HIC: ≥ 10.5 and < 11.5 min | T<sub>m</sub> < 65.0 °C |
| Medium PSR: ≥ 0.33 and < 0.66 | High HIC: ≥ 11.5 min |  |
| High PSR: ≥ 0.66 and ≤ 1.00 |  |  |

**Supplementary Table S4.**
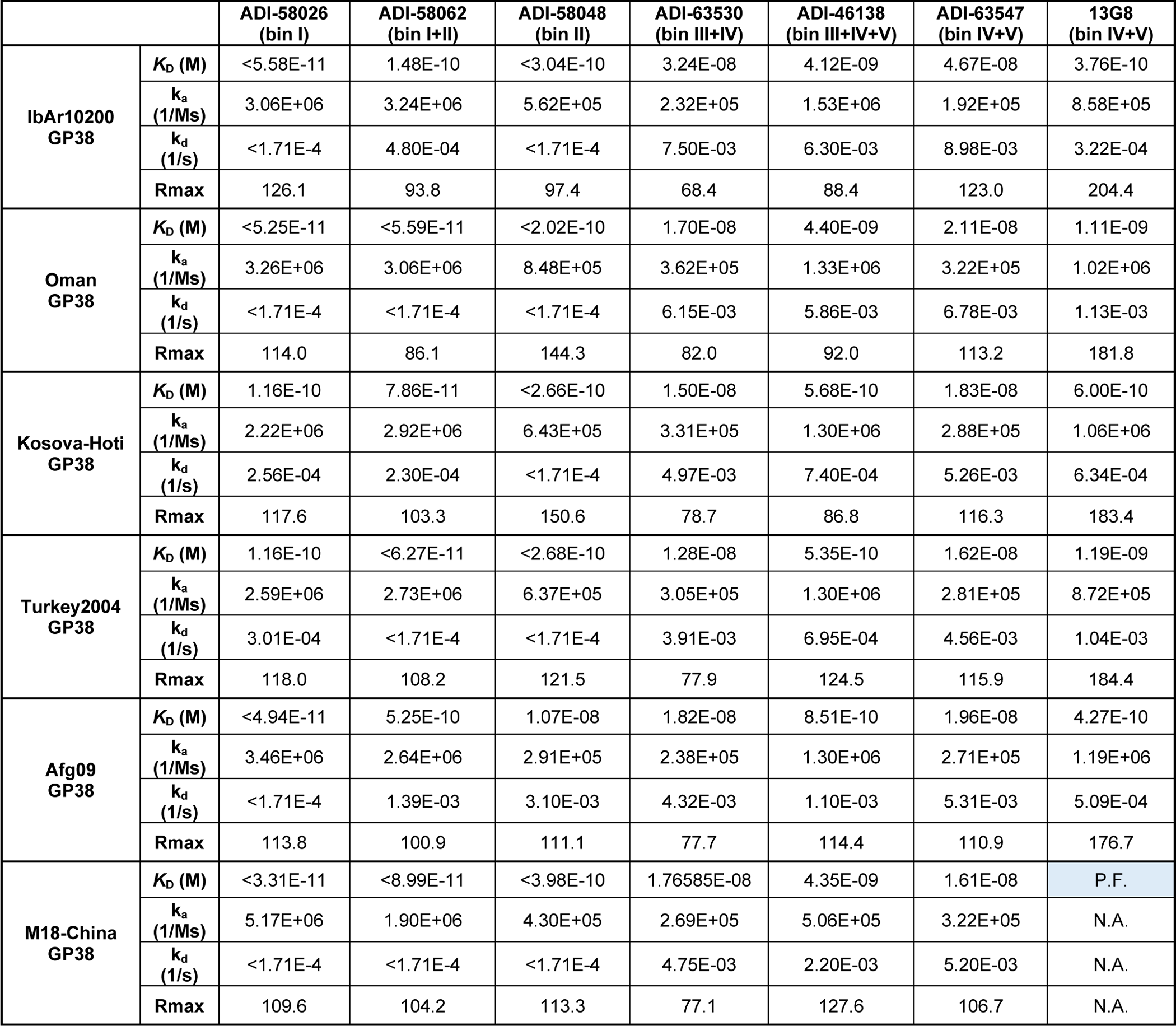
Carterra kinetics of GP38 antibodies. Multi-concentration Carterra kinetics data of the six mAbs used in protection studies against rGP38 protein of six tested clinical isolates. Samples for which the off-rate was limited are denoted as < the k_d_ and calculated K_D_. The sample for which a curve could not be fit is denoted as P.F and shaded in blue.

**Supplementary Table S5.**
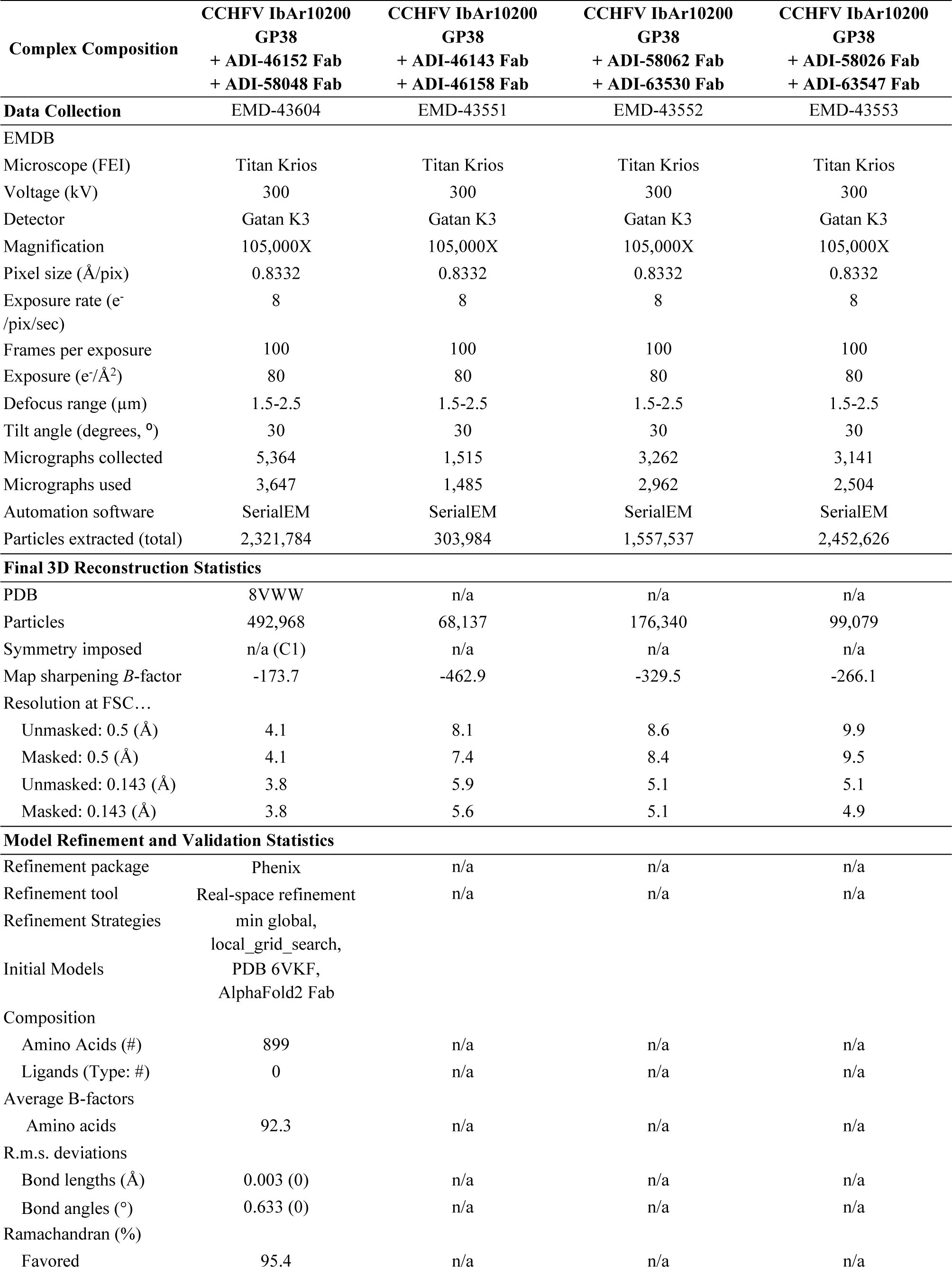
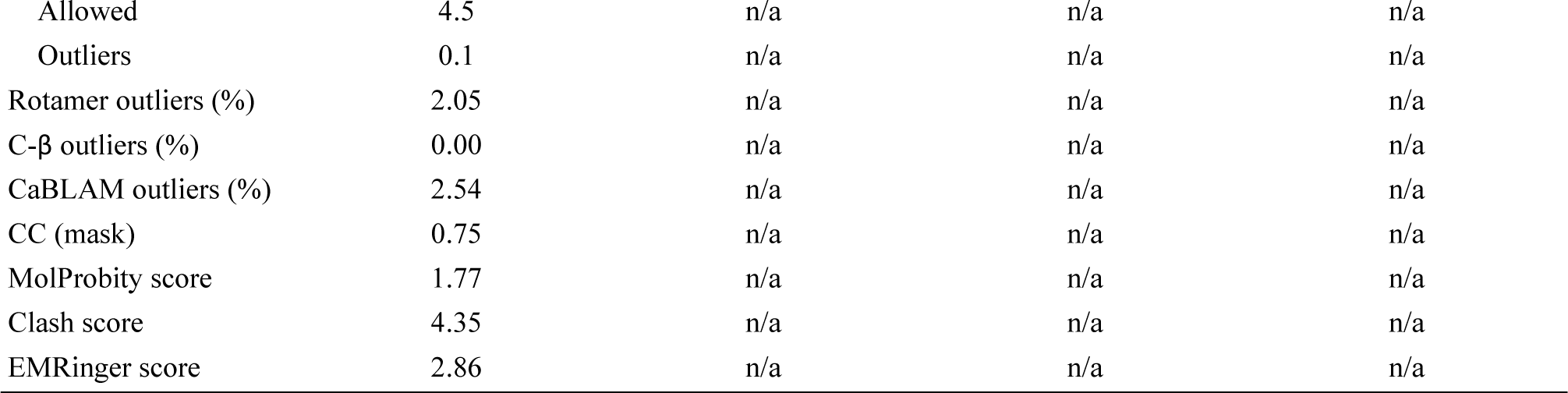
Cryo-EM data collection, reconstruction, and model validation statistics.

| **Complex Composition** | **CCHFV IbAr10200**<br>**GP38**<br>**+ ADI-46152 Fab**<br>**+ ADI-58048 Fab** | **CCHFV IbAr10200**<br>**GP38**<br>**+ ADI-46143 Fab**<br>**+ ADI-46158 Fab** | **CCHFV IbAr10200**<br>**GP38**<br>**+ ADI-58062 Fab**<br>**+ ADI-63530 Fab** | **CCHFV IbAr10200**<br>**GP38**<br>**+ ADI-58026 Fab**<br>**+ ADI-63547 Fab** |
| --- | --- | --- | --- | --- |
| **Data Collection** | EMD-43604 | EMD-43551 | EMD-43552 | EMD-43553 |
| EMDB |  |  |  |  |
| Microscope (FEI) | Titan Krios | Titan Krios | Titan Krios | Titan Krios |
| Voltage (kV) | 300 | 300 | 300 | 300 |
| Detector | Gatan K3 | Gatan K3 | Gatan K3 | Gatan K3 |
| Magnification | 105,000X | 105,000X | 105,000X | 105,000X |
| Pixel size (Å/pix) | 0.8332 | 0.8332 | 0.8332 | 0.8332 |
| Exposure rate (e<sup>-</sup><br>/pix/sec) | 8 | 8 | 8 | 8 |
| Frames per exposure | 100 | 100 | 100 | 100 |
| Exposure (e<sup>-</sup>/Å<sup>2</sup>) | 80 | 80 | 80 | 80 |
| Defocus range (µm) | 1.5-2.5 | 1.5-2.5 | 1.5-2.5 | 1.5-2.5 |
| Tilt angle (degrees, °) | 30 | 30 | 30 | 30 |
| Micrographs collected | 5,364 | 1,515 | 3,262 | 3,141 |
| Micrographs used | 3,647 | 1,485 | 2,962 | 2,504 |
| Automation software | SerialEM | SerialEM | SerialEM | SerialEM |
| Particles extracted (total) | 2,321,784 | 303,984 | 1,557,537 | 2,452,626 |
| **Final 3D Reconstruction Statistics** |  |  |  |  |
| PDB | 8VWW | n/a | n/a | n/a |
| Particles | 492,968 | 68,137 | 176,340 | 99,079 |
| Symmetry imposed | n/a (C1) | n/a | n/a | n/a |
| Map sharpening *B*-factor | -173.7 | -462.9 | -329.5 | -266.1 |
| Resolution at FSC... |  |  |  |  |
| Unmasked: 0.5 (Å) | 4.1 | 8.1 | 8.6 | 9.9 |
| Masked: 0.5 (Å) | 4.1 | 7.4 | 8.4 | 9.5 |
| Unmasked: 0.143 (Å) | 3.8 | 5.9 | 5.1 | 5.1 |
| Masked: 0.143 (Å) | 3.8 | 5.6 | 5.1 | 4.9 |
| **Model Refinement and Validation Statistics** |  |  |  |  |
| Refinement package | Phenix | n/a | n/a | n/a |
| Refinement tool | Real-space refinement | n/a | n/a | n/a |
| Refinement Strategies | min global,<br>local_grid_search, |  |  |  |
| Initial Models | PDB 6VKF,<br>AlphaFold2 Fab |  |  |  |
| Composition |  |  |  |  |
| Amino Acids (#) | 899 | n/a | n/a | n/a |
| Ligands (Type: #) | 0 | n/a | n/a | n/a |
| Average B-factors |  |  |  |  |
| Amino acids | 92.3 | n/a | n/a | n/a |
| R.m.s. deviations |  |  |  |  |
| Bond lengths (Å) | 0.003 (0) | n/a | n/a | n/a |
| Bond angles (°) | 0.633 (0) | n/a | n/a | n/a |
| Ramachandran (%) |  |  |  |  |
| Favored | 95.4 | n/a | n/a | n/a |
| Allowed | 4.5 | n/a | n/a | n/a |
| Outliers | 0.1 | n/a | n/a | n/a |
| Rotamer outliers (%) | 2.05 | n/a | n/a | n/a |
| C-β outliers (%) | 0.00 | n/a | n/a | n/a |
| CaBLAM outliers (%) | 2.54 | n/a | n/a | n/a |
| CC (mask) | 0.75 | n/a | n/a | n/a |
| MolProbity score | 1.77 | n/a | n/a | n/a |
| Clash score | 4.35 | n/a | n/a | n/a |
| EMRinger score | 2.86 | n/a | n/a | n/a |

**Supplementary Table S6.** Crystallographic data collection and refinement statistics.

| **Complex Composition** | **CCHFV IBAr10200 GP38 and ADI-46143 Fab** | **CCHFV IBAr10200 GP38 and c13G8 Fab** |
| --- | --- | --- |
| PDB ID | 8VVK | 8VVL |
| Reservoir solution for crystallization | 0.2 M (NH<sub>4</sub>)-Citrate pH 7.5<br>9.3% (w/v) PEG 3350<br>12.6% (v/v) 2-PropOH | 2 M ammonium sulfate<br>0.1 M Bis-Tris pH 5.5<br>0.01 M cobalt chloride hexahydrate |
| **Data collection** |  |  |
| Space group | *P* 6<sub>1</sub> 2 2 | *I* 1 2 1 |
| Wavelength (Å) | 0.979 | 0.979 |
| Cell dimensions |  |  |
| *a*, *b*, *c* (Å) | 149.7, 149.7, 315.7 | 100.4, 67.6, 141.0 |
| $\alpha$, $\beta$, $\gamma$ (°) | 90, 90, 120 | 90, 96.5, 90 |
| Resolution range (Å) | 64.83-2.61 (2.67-2.61) | 45.16-1.80 (1.84-1.80) |
| *R*<sub>merge</sub> | 0.027 (0.262) | 0.033 (0.269) |
| *I*/$\sigma$(*I*) | 15.6 (2.8) | 9.5 (2.5) |
| *CC*<sub>1/2</sub> | 0.999 (0.814) | 0.998 (0.915) |
| Completeness (%) | 99.93 (99.86) | 94.50 (92.89) |
| Redundancy | 2.0 (2.0) | 1.9 (1.9) |
| Total reflections | 128,324 (12,566) | 157,627 (15,257) |
| Unique reflections | 64,162 (6,283) | 81,877 (8,008) |
| **Refinement** |  |  |
| Resolution range (Å) | 59.95-2.61 (2.70-2.61) | 45.16-1.80 (1.87-1.80) |
| Unique reflections | 64,131 (6,274) | 81,792 (7,990) |
| *R*<sub>work</sub> / *R*<sub>free</sub> (%) | 17.71/21.71 (24.66/33.15) | 19.97/21.54 (30.17/33.95) |
| Number of atoms | 10,654 | 5,823 |
| Protein | 10,294 | 5,121 |
| Solvent | 237 | 623 |
| Ligands | 123 | 79 |
| Average *B*-factor (Å<sup>2</sup>) | 59.3 | 41.0 |
| Protein | 59.2 | 40.0 |
| Solvent | 50.2 | 47.0 |
| Ligands | 89.4 | 56.5 |
| R.m.s. deviations |  |  |
| Bond lengths (Å) | 0.005 | 0.004 |
| Bond angles (°) | 0.75 | 0.76 |
| Ramachandran (%) |  |  |
| Favored | 96.5 | 98.1 |
| Allowed | 3.3 | 1.9 |
| Outliers | 0.2 | 0.0 |

Data in parentheses are for the highest resolution shell.

## REFERENCES

1. Spengler, J.R., Bente, D.A., Bray, M., Burt, F., Hewson, R., Korukluoglu, G., Mirazimi, A., Weber, F., and Papa, A. (2018). Second International Conference on Crimean-Congo Hemorrhagic Fever. Antiviral Res 150, 137–147. 10.1016/j.antiviral.2017.11.019.

2. Spengler, J.R., Bergeron, É., and Spiropoulou, C.F. (2019). Crimean-Congo hemorrhagic fever and expansion from endemic regions. Curr Opin Virol 34, 70–78. 10.1016/j.coviro.2018.12.002.

3. Spengler, J.R., Estrada-Peña, A., Garrison, A.R., Schmaljohn, C., Spiropoulou, C.F., Bergeron, É., and Bente, D.A. (2016). A chronological review of experimental infection studies of the role of wild animals and livestock in the maintenance and transmission of Crimean-Congo hemorrhagic fever virus. Antiviral Res 135, 31–47. 10.1016/j.antiviral.2016.09.013.

4. Spengler, J.R., Bergeron, É., and Rollin, P.E. (2016). Seroepidemiological Studies of Crimean-Congo Hemorrhagic Fever Virus in Domestic and Wild Animals. PLoS Negl Trop Dis 10, e0004210. 10.1371/journal.pntd.0004210.

5. Nurettin, C., Engin, B., Sukru, T., Munir, A., Zati, V., and Aykut, O. (2022). The Seroprevalence of Crimean-Congo Hemorrhagic Fever in Wild and Domestic Animals: An Epidemiological Update for Domestic Animals and First Seroevidence in Wild Animals from Turkiye. Vet Sci 9. 10.3390/vetsci9090462.

6. Bente, D.A., Forrester, N.L., Watts, D.M., McAuley, A.J., Whitehouse, C.A., and Bray, M. (2013). Crimean-Congo hemorrhagic fever: history, epidemiology, pathogenesis, clinical syndrome and genetic diversity. Antiviral Res 100, 159–189. 10.1016/j.antiviral.2013.07.006.

7. Atim, S.A., Niebel, M., Ashraf, S., Vudriko, P., Odongo, S., Balinandi, S., Aber, P., Bameka, R., Ademun, A.R., Masembe, C., et al. (2023). Prevalence of Crimean-Congo haemorrhagic fever in livestock following a confirmed human case in Lyantonde district, Uganda. Parasit Vectors 16, 7. 10.1186/s13071-022-05588-x.

8. Smego, R.A., Jr., Sarwari, A.R., and Siddiqui, A.R. (2004). Crimean-Congo Hemorrhagic Fever: Prevention and Control Limitations in a Resource-Poor Country. Clinical Infectious Diseases 38, 1731–1735. 10.1086/421093.

9. Conger, N.G., Paolino, K.M., Osborn, E.C., Rusnak, J.M., Günther, S., Pool, J., Rollin, P.E., Allan, P.F., Schmidt-Chanasit, J., Rieger, T., and Kortepeter, M.G. (2015). Health care response to CCHF in US soldier and nosocomial transmission to health care providers, Germany, 2009. Emerg Infect Dis 21, 23–31. 10.3201/eid2101.141413.

10. Belobo, J.T.E., Kenmoe, S., Kengne-Nde, C., Emoh, C.P.D., Bowo-Ngandji, A., Tchatchouang, S., Sowe Wobessi, J.N., Mbongue Mikangue, C.A., Tazokong, H.R., Kingue Bebey, S.R., et al. (2021). Worldwide epidemiology of Crimean-Congo Hemorrhagic Fever Virus in humans, ticks and other animal species, a systematic review and meta-analysis. PLoS Negl Trop Dis 15, e0009299. 10.1371/journal.pntd.0009299.

11. Messina, J.P., Pigott, D.M., Golding, N., Duda, K.A., Brownstein, J.S., Weiss, D.J., Gibson, H., Robinson, T.P., Gilbert, M., William Wint, G.R., et al. (2015). The global distribution of Crimean-Congo hemorrhagic fever. Trans R Soc Trop Med Hyg 109, 503–513. 10.1093/trstmh/trv050.

12. Deyde, V.M., Khristova, M.L., Rollin, P.E., Ksiazek, T.G., and Nichol, S.T. (2006). Crimean-Congo hemorrhagic fever virus genomics and global diversity. J Virol 80, 8834–8842. 10.1128/jvi.00752-06.

13. Anagnostou, V., and Papa, A. (2009). Evolution of Crimean-Congo Hemorrhagic Fever virus. Infect Genet Evol 9, 948–954. 10.1016/j.meegid.2009.06.018.

14. Carroll, S.A., Bird, B.H., Rollin, P.E., and Nichol, S.T. (2010). Ancient common ancestry of Crimean-Congo hemorrhagic fever virus. Mol Phylogenet Evol 55, 1103–1110. 10.1016/j.ympev.2010.01.006.

15. Chamberlain, J., Cook, N., Lloyd, G., Mioulet, V., Tolley, H., and Hewson, R. (2005). Co-evolutionary patterns of variation in small and large RNA segments of Crimean-Congo hemorrhagic fever virus. J Gen Virol 86, 3337–3341. 10.1099/vir.0.81213-0.

16. Grard, G., Drexler, J.F., Fair, J., Muyembe, J.J., Wolfe, N.D., Drosten, C., and Leroy, E.M. (2011). Re-emergence of Crimean-Congo hemorrhagic fever virus in Central Africa. PLoS Negl Trop Dis 5, e1350. 10.1371/journal.pntd.0001350.

17. Hewson, R., Chamberlain, J., Mioulet, V., Lloyd, G., Jamil, B., Hasan, R., Gmyl, A., Gmyl, L., Smirnova, S.E., Lukashev, A., et al. (2004). Crimean-Congo haemorrhagic fever virus: sequence analysis of the small RNA segments from a collection of viruses world wide. Virus Res 102, 185–189. 10.1016/j.virusres.2003.12.035.

18. Lukashev, A.N., Klimentov, A.S., Smirnova, S.E., Dzagurova, T.K., Drexler, J.F., and Gmyl, A.P. (2016). Phylogeography of Crimean Congo Hemorrhagic Fever Virus. PLOS ONE 11, e0166744. 10.1371/journal.pone.0166744.

19. Sherifi, K., Cadar, D., Muji, S., Robaj, A., Ahmeti, S., Jakupi, X., Emmerich, P., and Krüger, A. (2014). Crimean-Congo Hemorrhagic Fever Virus Clades V and VI (Europe 1 and 2) in Ticks in Kosovo, 2012. PLOS Neglected Tropical Diseases 8, e3168. 10.1371/journal.pntd.0003168.

20. Papa, A., Marklewitz, M., Paraskevopoulou, S., Garrison, A.R., Alkhovsky, S.V., Avšič-Županc, T., Bente, D.A., Bergeron, É., Burt, F., Di Paola, N., et al. (2022). History and classification of Aigai virus (formerly Crimean–Congo haemorrhagic fever virus genotype VI). Journal of General Virology 103. 10.1099/jgv.0.001734.

21. Mehand, M.S., Al-Shorbaji, F., Millett, P., and Murgue, B. (2018). The WHO R&D Blueprint: 2018 review of emerging infectious diseases requiring urgent research and development efforts. Antiviral research 159, 63–67.

22. Johnson, S., Henschke, N., Maayan, N., Mills, I., Buckley, B.S., Kakourou, A., and Marshall, R. (2018). Ribavirin for treating Crimean Congo haemorrhagic fever. Cochrane Database of Systematic Reviews. 10.1002/14651858.CD012713.pub2.

23. Zivcec, M., Scholte, F.E., Spiropoulou, C.F., Spengler, J.R., and Bergeron, É. (2016). Molecular Insights into Crimean-Congo Hemorrhagic Fever Virus. Viruses 8, 106. 10.3390/v8040106.

24. Sanchez, A.J., Vincent, M.J., Erickson, B.R., and Nichol, S.T. (2006). Crimean-congo hemorrhagic fever virus glycoprotein precursor is cleaved by Furin-like and SKI-1 proteases to generate a novel 38-kilodalton glycoprotein. J Virol 80, 514–525. 10.1128/jvi.80.1.514-525.2006.

25. Sanchez, A.J., Vincent, M.J., and Nichol, S.T. (2002). Characterization of the glycoproteins of Crimean-Congo hemorrhagic fever virus. J Virol 76, 7263–7275. 10.1128/jvi.76.14.7263-7275.2002.

26. Freitas, N., Enguehard, M., Denolly, S., Levy, C., Neveu, G., Lerolle, S., Devignot, S., Weber, F., Bergeron, E., Legros, V., and Cosset, F.L. (2020). The interplays between Crimean-Congo hemorrhagic fever virus (CCHFV) M segment-encoded accessory proteins and structural proteins promote virus assembly and infectivity. PLoS Pathog 16, e1008850. 10.1371/journal.ppat.1008850.

27. Durie, I.A., Tehrani, Z.R., Karaaslan, E., Sorvillo, T.E., McGuire, J., Golden, J.W., Welch, S.R., Kainulainen, M.H., Harmon, J.R., Mousa, J.J., et al. (2022). Structural characterization of protective non-neutralizing antibodies targeting Crimean-Congo hemorrhagic fever virus. Nat Commun 13, 7298. 10.1038/s41467-022-34923-0.

28. Mishra, A.K., Moyer, C.L., Abelson, D.M., Deer, D.J., El Omari, K., Duman, R., Lobel, L., Lutwama, J.J., Dye, J.M., Wagner, A., et al. (2020). Structure and Characterization of Crimean-Congo Hemorrhagic Fever Virus GP38. J Virol 94. 10.1128/jvi.02005-19.

29. Golden, J.W., Shoemaker, C.J., Lindquist, M.E., Zeng, X., Daye, S.P., Williams, J.A., Liu, J., Coffin, K.M., Olschner, S., Flusin, O., et al. (2019). GP38-targeting monoclonal antibodies protect adult mice against lethal Crimean-Congo hemorrhagic fever virus infection. Sci Adv 5, eaaw9535. 10.1126/sciadv.aaw9535.

30. Fels, J.M., Maurer, D.P., Herbert, A.S., Wirchnianski, A.S., Vergnolle, O., Cross, R.W., Abelson, D.M., Moyer, C.L., Mishra, A.K., and Aguilan, J.T. (2021). Protective neutralizing antibodies from human survivors of Crimean-Congo hemorrhagic fever. Cell 184, 3486–3501. e3421.

31. Andrews, S.F., Chambers, M.J., Schramm, C.A., Plyler, J., Raab, J.E., Kanekiyo, M., Gillespie, R.A., Ransier, A., Darko, S., Hu, J., et al. (2019). Activation Dynamics and Immunoglobulin Evolution of Pre-existing and Newly Generated Human Memory B cell Responses to Influenza Hemagglutinin. Immunity 51, 398–410.e395. 10.1016/j.immuni.2019.06.024.

32. Glass, D.R., Tsai, A.G., Oliveria, J.P., Hartmann, F.J., Kimmey, S.C., Calderon, A.A., Borges, L., Glass, M.C., Wagar, L.E., Davis, M.M., and Bendall, S.C. (2020). An Integrated Multi-omic Single-Cell Atlas of Human B Cell Identity. Immunity 53, 217–232.e215. 10.1016/j.immuni.2020.06.013.

33. Wec, A.Z., Haslwanter, D., Abdiche, Y.N., Shehata, L., Pedreño-Lopez, N., Moyer, C.L., Bornholdt, Z.A., Lilov, A., Nett, J.H., Jangra, R.K., et al. (2020). Longitudinal dynamics of the human B cell response to the yellow fever 17D vaccine. Proc Natl Acad Sci U S A 117, 6675–6685. 10.1073/pnas.1921388117.

34. Sakharkar, M., Rappazzo, C.G., Wieland-Alter, W.F., Hsieh, C.-L., Wrapp, D., Esterman, E.S., Kaku, C.I., Wec, A.Z., Geoghegan, J.C., McLellan, J.S., et al. (2021). Prolonged evolution of the human B cell response to SARS-CoV-2 infection. Science Immunology 6, eabg6916. doi:10.1126/sciimmunol.abg6916.

35. Rogers, T.F., Goodwin, E.C., Briney, B., Sok, D., Beutler, N., Strubel, A., Nedellec, R., Le, K., Brown, M.E., Burton, D.R., and Walker, L.M. (2017). Zika virus activates de novo and cross-reactive memory B cell responses in dengue-experienced donors. Sci Immunol 2. 10.1126/sciimmunol.aan6809.

36. Bornholdt, Z.A., Turner, H.L., Murin, C.D., Li, W., Sok, D., Souders, C.A., Piper, A.E., Goff, A., Shamblin, J.D., Wollen, S.E., et al. (2016). Isolation of potent neutralizing antibodies from a survivor of the 2014 Ebola virus outbreak. Science 351, 1078–1083. doi:10.1126/science.aad5788.

37. Wrammert, J., Smith, K., Miller, J., Langley, W.A., Kokko, K., Larsen, C., Zheng, N.Y., Mays, I., Garman, L., Helms, C., et al. (2008). Rapid cloning of high-affinity human monoclonal antibodies against influenza virus. Nature 453, 667–671. 10.1038/nature06890.

38. Schramm, C.A., and Douek, D.C. (2018). Beyond Hot Spots: Biases in Antibody Somatic Hypermutation and Implications for Vaccine Design. Front Immunol 9, 1876. 10.3389/fimmu.2018.01876.

39. Elsner, R.A., and Shlomchik, M.J. (2020). Germinal Center and Extrafollicular B Cell Responses in Vaccination, Immunity, and Autoimmunity. Immunity 53, 1136–1150. 10.1016/j.immuni.2020.11.006.

40. Briney, B., Inderbitzin, A., Joyce, C., and Burton, D.R. (2019). Commonality despite exceptional diversity in the baseline human antibody repertoire. Nature 566, 393–397. 10.1038/s41586-019-0879-y.

41. Hoogstraal, H. (1979). The epidemiology of tick-borne Crimean-Congo hemorrhagic fever in Asia, Europe, and Africa. J Med Entomol 15, 307–417. 10.1093/jmedent/15.4.307.

42. Kong, Y., Yan, C., Liu, D., Jiang, L., Zhang, G., He, B., and Li, Y. (2022). Phylogenetic analysis of Crimean-Congo hemorrhagic fever virus in inner Mongolia, China. Ticks and Tick-borne Diseases 13, 101856. 10.1016/j.ttbdis.2021.101856.

43. Guo, R., Shen, S., Zhang, Y., Shi, J., Su, Z., Liu, D., Liu, J., Yang, J., Wang, Q., Hu, Z., et al. (2017). A new strain of Crimean-Congo hemorrhagic fever virus isolated from Xinjiang, China. Virologica Sinica 32, 80–88. 10.1007/s12250-016-3936-9.

44. Jain, T., Sun, T., Durand, S., Hall, A., Houston, N.R., Nett, J.H., Sharkey, B., Bobrowicz, B., Caffry, I., Yu, Y., et al. (2017). Biophysical properties of the clinical-stage antibody landscape. Proc Natl Acad Sci U S A 114, 944–949. 10.1073/pnas.1616408114.

45. Zivcec, M., Metcalfe, M.G., Albariño, C.G., Guerrero, L.W., Pegan, S.D., Spiropoulou, C.F., and Bergeron, É. (2015). Assessment of Inhibitors of Pathogenic Crimean-Congo Hemorrhagic Fever Virus Strains Using Virus-Like Particles. PLoS Negl Trop Dis 9, e0004259. 10.1371/journal.pntd.0004259.

46. Dai, S., Deng, F., Wang, H., and Ning, Y. (2021). Crimean-Congo Hemorrhagic Fever Virus: Current Advances and Future Prospects of Antiviral Strategies. Viruses 13. 10.3390/v13071195.

47. Zivcec, M., Guerrero, L.I.W., Albariño, C.G., Bergeron, É., Nichol, S.T., and Spiropoulou, C.F. (2017). Identification of broadly neutralizing monoclonal antibodies against Crimean-Congo hemorrhagic fever virus. Antiviral Res 146, 112–120. 10.1016/j.antiviral.2017.08.014.

48. Bereczky, S., Lindegren, G., Karlberg, H., Akerström, S., Klingström, J., and Mirazimi, A. (2010). Crimean-Congo hemorrhagic fever virus infection is lethal for adult type I interferon receptor-knockout mice. J Gen Virol 91, 1473–1477. 10.1099/vir.0.019034-0.

49. Zivcec, M., Safronetz, D., Scott, D., Robertson, S., Ebihara, H., and Feldmann, H. (2013). Lethal Crimean-Congo hemorrhagic fever virus infection in interferon α/β receptor knockout mice is associated with high viral loads, proinflammatory responses, and coagulopathy. J Infect Dis 207, 1909–1921. 10.1093/infdis/jit061.

50. Bente, D.A., Alimonti, J.B., Shieh, W.J., Camus, G., Ströher, U., Zaki, S., and Jones, S.M. (2010). Pathogenesis and immune response of Crimean-Congo hemorrhagic fever virus in a STAT-1 knockout mouse model. J Virol 84, 11089–11100. 10.1128/jvi.01383-10.

51. Suschak, J.J., Golden, J.W., Fitzpatrick, C.J., Shoemaker, C.J., Badger, C.V., Schmaljohn, C.S., and Garrison, A.R. (2021). A CCHFV DNA vaccine protects against heterologous challenge and establishes GP38 as immunorelevant in mice. npj Vaccines 6, 31. 10.1038/s41541-021-00293-9.

52. Bornholdt, Z.A., Herbert, A.S., Mire, C.E., He, S., Cross, R.W., Wec, A.Z., Abelson, D.M., Geisbert, J.B., James, R.M., Rahim, M.N., et al. (2019). A Two-Antibody Pan-Ebolavirus Cocktail Confers Broad Therapeutic Protection in Ferrets and Nonhuman Primates. Cell Host Microbe 25, 49–58.e45. 10.1016/j.chom.2018.12.005.

53. Wec, A.Z., Bornholdt, Z.A., He, S., Herbert, A.S., Goodwin, E., Wirchnianski, A.S., Gunn, B.M., Zhang, Z., Zhu, W., Liu, G., et al. (2019). Development of a Human Antibody Cocktail that Deploys Multiple Functions to Confer Pan-Ebolavirus Protection. Cell Host Microbe 25, 39–48.e35. 10.1016/j.chom.2018.12.004.

54. Bornholdt, Z.A., Ndungo, E., Fusco, M.L., Bale, S., Flyak, A.I., Crowe, J.E., Jr., Chandran, K., and Saphire, E.O. (2016). Host-Primed Ebola Virus GP Exposes a Hydrophobic NPC1 Receptor-Binding Pocket, Revealing a Target for Broadly Neutralizing Antibodies. mBio 7, e02154–02115. 10.1128/mBio.02154-15.

55. Chapman, N.S., Zhao, H., Kose, N., Westover, J.B., Kalveram, B., Bombardi, R., Rodriguez, J., Sutton, R., Genualdi, J., LaBeaud, A.D., et al. (2021). Potent neutralization of Rift Valley fever virus by human monoclonal antibodies through fusion inhibition. Proceedings of the National Academy of Sciences 118, e2025642118. doi:10.1073/pnas.2025642118.

56. Corti, D., Misasi, J., Mulangu, S., Stanley, D.A., Kanekiyo, M., Wollen, S., Ploquin, A., Doria-Rose, N.A., Staupe, R.P., Bailey, M., et al. (2016). Protective monotherapy against lethal Ebola virus infection by a potently neutralizing antibody. Science 351, 1339–1342. doi:10.1126/science.aad5224.

57. Engdahl, T.B., Kuzmina, N.A., Ronk, A.J., Mire, C.E., Hyde, M.A., Kose, N., Josleyn, M.D., Sutton, R.E., Mehta, A., Wolters, R.M., et al. (2021). Broad and potently neutralizing monoclonal antibodies isolated from human survivors of New World hantavirus infection. Cell Rep 35, 109086. 10.1016/j.celrep.2021.109086.

58. Mittler, E., Wec, A.Z., Tynell, J., Guardado-Calvo, P., Wigren-Byström, J., Polanco, L.C., O’Brien, C.M., Slough, M.M., Abelson, D.M., Serris, A., et al. (2022). Human antibody recognizing a quaternary epitope in the Puumala virus glycoprotein provides broad protection against orthohantaviruses. Sci Transl Med 14, eabl5399. 10.1126/scitranslmed.abl5399.

59. Quiroz, J.A., Malonis, R.J., Thackray, L.B., Cohen, C.A., Pallesen, J., Jangra, R.K., Brown, R.S., Hofmann, D., Holtsberg, F.W., Shulenin, S., et al. (2019). Human monoclonal antibodies against chikungunya virus target multiple distinct epitopes in the E1 and E2 glycoproteins. PLOS Pathogens 15, e1008061. 10.1371/journal.ppat.1008061.

60. West, B.R., Wec, A.Z., Moyer, C.L., Fusco, M.L., Ilinykh, P.A., Huang, K., Wirchnianski, A.S., James, R.M., Herbert, A.S., Hui, S., et al. (2019). Structural basis of broad ebolavirus neutralization by a human survivor antibody. Nat Struct Mol Biol 26, 204–212. 10.1038/s41594-019-0191-4.

61. Bertolotti-Ciarlet, A., Smith, J., Strecker, K., Paragas, J., Altamura, L.A., McFalls, J.M., Frias-Stäheli, N., García-Sastre, A., Schmaljohn, C.S., and Doms, R.W. (2005). Cellular localization and antigenic characterization of crimean-congo hemorrhagic fever virus glycoproteins. J Virol 79, 6152–6161. 10.1128/jvi.79.10.6152-6161.2005.

62. Earnest, J.T., Holmes, A.C., Basore, K., Mack, M., Fremont, D.H., and Diamond, M.S. (2021). The mechanistic basis of protection by non-neutralizing anti-alphavirus antibodies. Cell Reports 35, 108962. 10.1016/j.celrep.2021.108962.

63. Gunn, B.M., Yu, W.H., Karim, M.M., Brannan, J.M., Herbert, A.S., Wec, A.Z., Halfmann, P.J., Fusco, M.L., Schendel, S.L., Gangavarapu, K., et al. (2018). A Role for Fc Function in Therapeutic Monoclonal Antibody-Mediated Protection against Ebola Virus. Cell Host Microbe 24, 221–233.e225. 10.1016/j.chom.2018.07.009.

64. Ilinykh, P.A., Huang, K., Santos, R.I., Gilchuk, P., Gunn, B.M., Karim, M.M., Liang, J., Fouch, M.E., Davidson, E., Parekh, D.V., et al. (2020). Non-neutralizing Antibodies from a Marburg Infection Survivor Mediate Protection by Fc-Effector Functions and by Enhancing Efficacy of Other Antibodies. Cell Host Microbe 27, 976–991.e911. 10.1016/j.chom.2020.03.025.

65. Lewis, G.K., Pazgier, M., Evans, D.T., Ferrari, G., Bournazos, S., Parsons, M.S., Bernard, N.F., and Finzi, A. (2017). Beyond Viral Neutralization. AIDS Res Hum Retroviruses 33, 760–764. 10.1089/aid.2016.0299.

66. Sicca, F., Neppelenbroek, S., and Huckriede, A. (2018). Effector mechanisms of influenza-specific antibodies: neutralization and beyond. Expert Rev Vaccines 17, 785–795. 10.1080/14760584.2018.1516553.

67. Krammer, F., and Palese, P. (2015). Advances in the development of influenza virus vaccines. Nat Rev Drug Discov 14, 167–182. 10.1038/nrd4529.

68. Saphire, E.O., Schendel, S.L., Fusco, M.L., Gangavarapu, K., Gunn, B.M., Wec, A.Z., Halfmann, P.J., Brannan, J.M., Herbert, A.S., Qiu, X., et al. (2018). Systematic Analysis of Monoclonal Antibodies against Ebola Virus GP Defines Features that Contribute to Protection. Cell 174, 938–952.e913. 10.1016/j.cell.2018.07.033.

69. Lux, A., Yu, X., Scanlan, C.N., and Nimmerjahn, F. (2013). Impact of immune complex size and glycosylation on IgG binding to human FcγRs. J Immunol 190, 4315–4323. 10.4049/jimmunol.1200501.

70. Guardado-Calvo, P., and Rey, F.A. (2021). The Viral Class II Membrane Fusion Machinery: Divergent Evolution from an Ancestral Heterodimer. Viruses 13. 10.3390/v13122368.

71. Dyer, O. (2019). Two Ebola treatments halve deaths in trial in DRC outbreak. Bmj 366, l5140. 10.1136/bmj.l5140.

72. Herbert, A.S., Froude, J.W., Ortiz, R.A., Kuehne, A.I., Dorosky, D.E., Bakken, R.R., Zak, S.E., Josleyn, N.M., Musiychuk, K., Jones, R.M., et al. (2020). Development of an antibody cocktail for treatment of Sudan virus infection. Proceedings of the National Academy of Sciences 117, 3768–3778. doi:10.1073/pnas.1914985117.

73. Pascal, K.E., Dudgeon, D., Trefry, J.C., Anantpadma, M., Sakurai, Y., Murin, C.D., Turner, H.L., Fairhurst, J., Torres, M., Rafique, A., et al. (2018). Development of Clinical-Stage Human Monoclonal Antibodies That Treat Advanced Ebola Virus Disease in Nonhuman Primates. J Infect Dis 218, S612–s626. 10.1093/infdis/jiy285.

74. Howell, K.A., Brannan, J.M., Bryan, C., McNeal, A., Davidson, E., Turner, H.L., Vu, H., Shulenin, S., He, S., Kuehne, A., et al. (2017). Cooperativity Enables Non-neutralizing Antibodies to Neutralize Ebolavirus. Cell Rep 19, 413–424. 10.1016/j.celrep.2017.03.049.

75. Bowick, G.C., Airo, A.M., and Bente, D.A. (2012). Expression of interferon-induced antiviral genes is delayed in a STAT1 knockout mouse model of Crimean-Congo hemorrhagic fever. Virol J 9, 122. 10.1186/1743-422x-9-122.

76. Tiller, T., Meffre, E., Yurasov, S., Tsuiji, M., Nussenzweig, M.C., and Wardemann, H. (2008). Efficient generation of monoclonal antibodies from single human B cells by single cell RT-PCR and expression vector cloning. J Immunol Methods 329, 112–124. 10.1016/j.jim.2007.09.017.

77. Gietz, R.D., and Woods, R.A. (2006). Yeast transformation by the LiAc/SS Carrier DNA/PEG method. Methods Mol Biol 313, 107–120. 10.1385/1-59259-958-3:107.

78. Wec, A.Z., Wrapp, D., Herbert, A.S., Maurer, D.P., Haslwanter, D., Sakharkar, M., Jangra, R.K., Dieterle, M.E., Lilov, A., Huang, D., et al. (2020). Broad neutralization of SARS-related viruses by human monoclonal antibodies. Science 369, 731–736. 10.1126/science.abc7424.

79. Mata-Fink, J., Kriegsman, B., Yu, H.X., Zhu, H., Hanson, M.C., Irvine, D.J., and Wittrup, K.D. (2013). Rapid conformational epitope mapping of anti-gp120 antibodies with a designed mutant panel displayed on yeast. J Mol Biol 425, 444–456. 10.1016/j.jmb.2012.11.010.

80. Winn, M.D., Ballard, C.C., Cowtan, K.D., Dodson, E.J., Emsley, P., Evans, P.R., Keegan, R.M., Krissinel, E.B., Leslie, A.G., McCoy, A., et al. (2011). Overview of the CCP4 suite and current developments. Acta Crystallogr D Biol Crystallogr 67, 235–242. 10.1107/s0907444910045749.

81. Battye, T.G., Kontogiannis, L., Johnson, O., Powell, H.R., and Leslie, A.G. (2011). iMOSFLM: a new graphical interface for diffraction-image processing with MOSFLM. Acta Crystallogr D Biol Crystallogr 67, 271–281. 10.1107/s0907444910048675.

82. Evans, P.R., and Murshudov, G.N. (2013). How good are my data and what is the resolution? Acta Crystallogr D Biol Crystallogr 69, 1204–1214. 10.1107/s0907444913000061.

83. McCoy, A.J. (2007). Solving structures of protein complexes by molecular replacement with Phaser. Acta Crystallogr D Biol Crystallogr 63, 32–41. 10.1107/s0907444906045975.

84. Emsley, P., and Cowtan, K. (2004). Coot: model-building tools for molecular graphics. Acta Crystallogr D Biol Crystallogr 60, 2126–2132. 10.1107/S0907444904019158.

85. Adams, P.D., Afonine, P.V., Bunkoczi, G., Chen, V.B., Davis, I.W., Echols, N., Headd, J.J., Hung, L.W., Kapral, G.J., Grosse-Kunstleve, R.W., et al. (2010). PHENIX: a comprehensive Python-based system for macromolecular structure solution. Acta Crystallogr D Biol Crystallogr 66, 213–221. 10.1107/S0907444909052925.

86. DeLano, W.L. (2002). Pymol: An open-source molecular graphics tool. CCP4 Newsl. Protein Crystallogr 40, 82–92.

87. Punjani, A., Rubinstein, J.L., Fleet, D.J., and Brubaker, M.A. (2017). cryoSPARC: algorithms for rapid unsupervised cryo-EM structure determination. Nature Methods 14, 290–296. 10.1038/nmeth.4169.

88. Sanchez-Garcia, R., Gomez-Blanco, J., Cuervo, A., Carazo, J.M., Sorzano, C.O.S., and Vargas, J. (2021). DeepEMhancer: a deep learning solution for cryo-EM volume post-processing. Communications Biology 4, 874. 10.1038/s42003-021-02399-1.

89. Croll, T.I. (2018). ISOLDE: a physically realistic environment for model building into low-resolution electron-density maps. Acta Crystallographica Section D: Structural Biology 74, 519–530.

90. Xu, Y., Roach, W., Sun, T., Jain, T., Prinz, B., Yu, T.-Y., Torrey, J., Thomas, J., Bobrowicz, P., Vásquez, M., et al. (2013). Addressing polyspecificity of antibodies selected from an in vitro yeast presentation system: a FACS-based, high-throughput selection and analytical tool. Protein Engineering, Design and Selection 26, 663–670. 10.1093/protein/gzt047.

91. Estep, P., Caffry, I., Yu, Y., Sun, T., Cao, Y., Lynaugh, H., Jain, T., Vásquez, M., Tessier, P.M., and Xu, Y. (2015). An alternative assay to hydrophobic interaction chromatography for high-throughput characterization of monoclonal antibodies. MAbs 7, 553–561. 10.1080/19420862.2015.1016694.

92. He, F., Woods, C.E., Becker, G.W., Narhi, L.O., and Razinkov, V.I. (2011). High-throughput assessment of thermal and colloidal stability parameters for monoclonal antibody formulations. J Pharm Sci 100, 5126–5141. 10.1002/jps.22712.

## Supplemental References

1. Jain, T., Sun, T., Durand, S., Hall, A., Houston, N.R., Nett, J.H., Sharkey, B., Bobrowicz, B., Caffry, I., Yu, Y., et al. (2017). Biophysical properties of the clinical-stage antibody landscape. Proc Natl Acad Sci U S A 114, 944–949. 10.1073/pnas.1616408114.

